# A new antibiotic from an uncultured bacterium binds to an immutable target

**DOI:** 10.1101/2023.05.15.540765

**Authors:** Rhythm Shukla, Aaron J. Peoples, Kevin C. Ludwig, Sourav Maity, Maik G.N. Derks, Stefania de Benedetti, Annika M Krueger, Bram J.A. Vermeulen, Francesca Lavore, Rodrigo V. Honorato, Fabian Grein, Alexandre Bonvin, Ulrich Kubitscheck, Eefjan Breukink, Catherine Achorn, Anthony Nitti, Christopher J. Schwalen, Amy L. Spoering, Losee Lucy Ling, Dallas Hughes, Moreno Lelli, Wouter H. Roos, Kim Lewis, Tanja Schneider, Markus Weingarth

**Affiliations:** NMR Spectroscopy, Department of Chemistry, Utrecht University, Padualaan 8, 3584 CH Utrecht, The Netherlands; Membrane Biochemistry and Biophysics, Department of Chemistry, Utrecht University, Padualaan 8, 3584 CH, Utrecht, The Netherlands; NovoBiotic Pharmaceuticals, Cambridge, Massachusetts 02138, USA; Institute for Pharmaceutical Microbiology, University Hospital Bonn, University of Bonn, Bonn, Germany; Moleculaire Biofysica, Zernike Instituut, Rijksuniversiteit Groningen, Nijenborgh 4, 9747 AG Groningen, The Netherlands; Institute for Physical and Theoretical Chemistry, University of Bonn, Bonn, Germany; German Center for Infection Research (DZIF), partner site Bonn-Cologne, Bonn, Germany; Novartis Institutes for Biomedical Research, Cambridge, MA 02139, USA; Magnetic Resonance Center (CERM) and Department of Chemistry “Ugo Schiff”, University of Florence, via Sacconi 6, Sesto Fiorentino, 50019 Italy; Consorzio Interuniversitario Risonanze Magnetiche MetalloProteine (CIRMMP), via Sacconi 6, Sesto Fiorentino, 50019 Italy; Antimicrobial Discovery Center, Northeastern University, Department of Biology, Boston, Massachusetts 02115, USA

**Keywords:** Antibiotics, uncultured bacteria, peptidoglycan, cell wall, lipid II, mechanism of action, solid state NMR, atomic force microscopy, antibiotic resistance, infection, animal models

## Abstract

Antimicrobial resistance is a leading mortality factor worldwide. Here we report the discovery of clovibactin, a new antibiotic, isolated from uncultured soil bacteria. Clovibactin efficiently kills drug-resistant bacterial pathogens without detectable resistance. Using biochemical assays, solid-state NMR, and atomic force microscopy, we dissect its mode of action. Clovibactin blocks cell wall synthesis by targeting pyrophosphate of multiple essential peptidoglycan precursors (C_55_PP, Lipid II, Lipid_WTA_). Clovibactin uses an unusual hydrophobic interface to tightly wrap around pyrophosphate, but bypasses the variable structural elements of precursors, accounting for the lack of resistance. Selective and efficient target binding is achieved by the irreversible sequestration of precursors into supramolecular fibrils that only form on bacterial membranes that contain lipid-anchored pyrophosphate groups. Uncultured bacteria offer a rich reservoir of antibiotics with new mechanisms of action that could replenish the antimicrobial discovery pipeline.

## Introduction

The introduction of antibiotics has revolutionized medicine, providing effective treatment for infectious diseases that were once fatal, and enabling modern medicine such as surgery and organ transplantation. Widespread resistance development thwarts the effectiveness and lifespan of antibiotics, calling for the discovery of new drugs in the perpetual standoff against human pathogens (Brown and Wright, 2016; Cook and Wright, 2022; Miethke et al., 2021; Mordoch et al., 1999).

Most antibiotics used in the clinic originate from natural product scaffolds that were discovered by screening soil dwelling bacteria (Lewis, 2020). This approach, ushered in by the ‘Waksman platform’, was vastly successful in the 1940s to 1960s, considered the Golden Age of antibiotics discovery, and eventually led to the introduction of compounds such as streptomycin, vancomycin, or tetracycline. However, to date, traditional screening sources seem overmined as they tend to yield previously-known compounds. As the drug discovery pipeline has considerably thinned, it is prudent to search for new antibiotics with unprecedented mechanism of action among untapped groups of bacterial producers.

Uncultured bacteria represent a vast (~99% of all species), unexploited source of new natural product scaffolds (Lewis, 2013; Wilson et al., 2014). Recently, the development of the iChip technique (Nichols et al., 2010) provided access to a broad diversity of uncultured bacterial species, leading to the discovery of teixobactin (Ling et al., 2015), isolated from the soil bacterium *Eleftheria terrae*. Teixobactin shows excellent antibacterial activity and has a unique chemical structure. Teixobactin blocks cell wall biosynthesis by specific binding to highly conserved lipid precursors (Ling et al., 2015), leading to the formation of supramolecular structures that perturb membrane stability (Shukla et al., 2022b). Additional antibiotics with novel modes of action that came out of screening uncultured bacteria are lassomycin, an inhibitor of the ClpP1P2C1 protease and amycobactin, an inhibitor of the SecY protein exporter, both acting selectively against mycobacteria (Gavrish et al., 2014; Quigley et al., 2020). Uncultured bacteria hence appear to offer a rich source of compounds with new chemical and mechanistic characteristics, which bodes well for the sustained discovery of effective leads to develop next-generation antibiotics.

Here we report the discovery and mode of action of clovibactin, a new antibiotic without detectable resistance, identified from a screen of uncultured bacteria.

## Results

### Discovery of clovibactin

Some environmental bacteria, as well as spores, may require a prolonged incubation to initiate growth in vitro (Buerger et al., 2012a, b). This allows access to microorganisms that would be missed by standard cultivation techniques. To favor growth of spore-forming actinomycetes, we first incubated soil at 65°C for 30 minutes, then diluted it to extinction into a growth medium in microtiter plates to achieve ≤1 cell per well. Some non-spore formers can grow after this mild heating as well. Growth was followed by observing the wells under a dissecting microscope (50X magnification) at 1, 2, 4, 8, 12, and 16 weeks. Colonies were then sub-cultured and screened for antimicrobial activity on nutrient agar plates overlaid with *Staphylococcus aureus*. One of the producers was detected at 12 weeks of incubation and based on 16s rDNA sequence, is 99% identical to *Eleftheria terrae*. We refer to this isolate as *E. terrae ssp. carolina*; the sandy soil it was isolated from comes from North Carolina. The genus *Eleftheria* belongs to β-proteobacteria, is very rare, and the original *E. terrae* we isolated several years ago by growth in situ is the producer of teixobactin (Ling et al., 2015). We therefore decided to identify the antimicrobial compound produced by *E. terrae ssp. carolina*.

The first antibiotic identified from the producing organism was kalimantacin. However, extracts of the fermented broth showed activity against *B. subtilis*, while kalimantacin is not active against this species (Fage et al., 2020). Upon an initial separation of the crude extract by HPLC, we identified another fraction of the extract that showed activity against both *B. subtilis* and *S. aureus*. In an attempt to increase metabolic flux to novel compound production and simplify isolation, we set out to disrupt genes responsible for kalimantacin production.

Whole genome sequencing led to the identification of a biosynthetic gene cluster (BGC) with 55% identity to the kalimantacin/batumin operon (Mattheus et al., 2010). Production of kalimantacin was reduced below detectable levels by interrupting the first gene in the operon, *bat1*, using homologous recombination of a suicide vector (Fernández-Martínez and Bibb, 2014).

Fermentation broth of *E. terrae ssp. carolina Δbat1* was separated by HPLC, and bioassay-guided isolation produced a fraction with a compound having a unique mass of 903.5291 [M+H]^+^ as analysed by Antibase. According to Antibase, this mass is unique. A combination of mass spectrometry and solution NMR resolved the structure of this compound, which is a novel depsipeptide that we named clovibactin **(Figure 1 and Supplementary Tables 1 and 2)**. Stereochemistry was confirmed by Marfey’s analysis. Clovibactin features two D-amino acids in its linear N-terminus and an uncommon residue D-3-hydroxyasparagine in the depsi-cycle. The compound’s molecular scaffold bears some resemblance to the depsipeptide teixobactin, as reflected by a Tanimoto coefficient of 0.8761. However, clovibactin has a considerably shorter linear N-terminus (4 residues in clovibactin, 7 residues in teixobactin) that harbors the two positive charges present in the compound. Additionally, teixobactin contains one of its two positively charged amino acids in the macrolactone that is further represented by the presence of the unusual enduracididine residue, missing in clovibactin.

**Figure 1.**
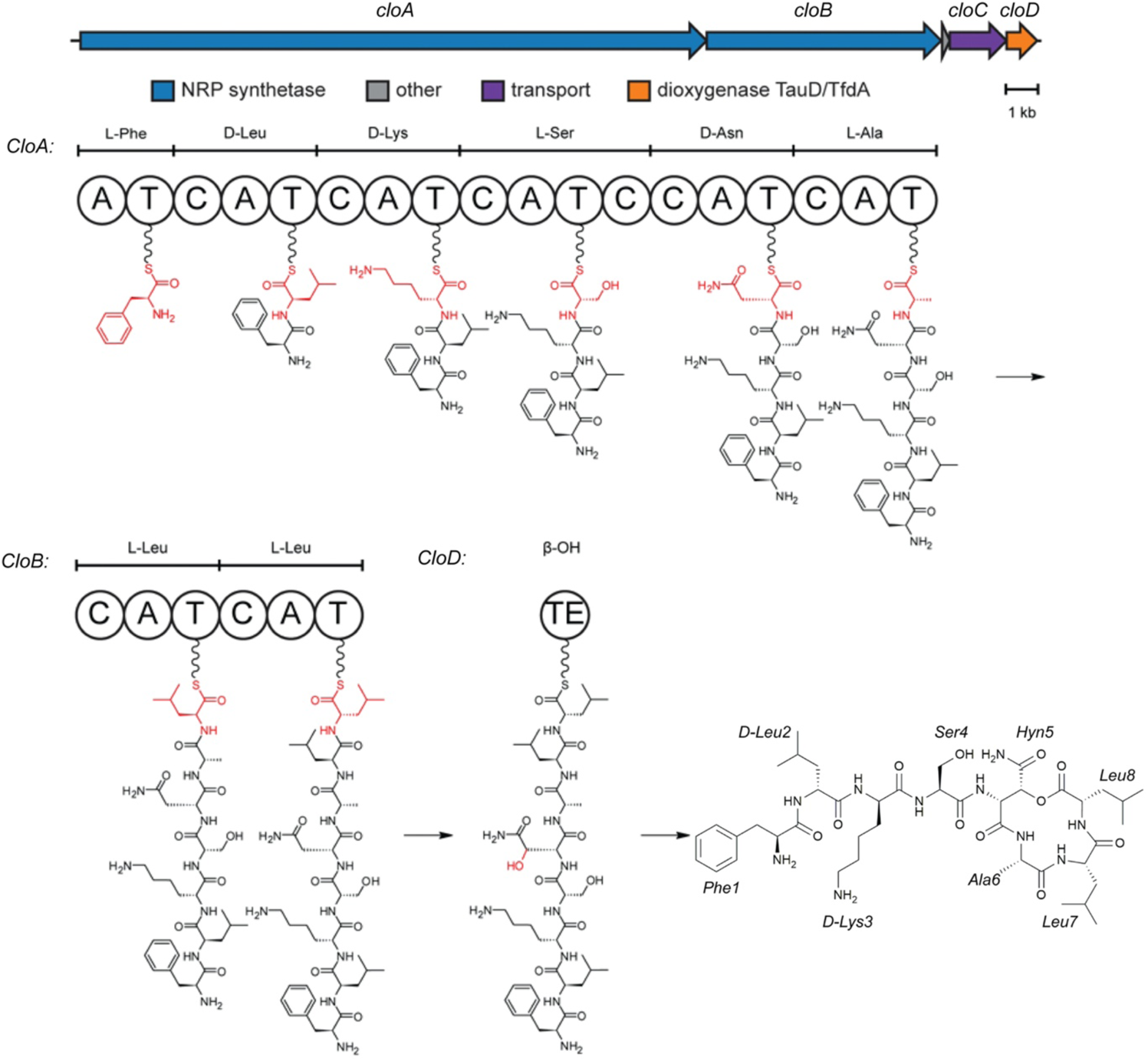
Biosynthetic gene cluster and proposed biosynthesis of clovibactin. The gene cluster associated with the biosynthesis of clovibactin was identified via whole genome sequencing and contained two nonribosomal peptide synthetases (NRPS) genes (*cloA* and *cloB*), a transporter gene (*cloC*) and a tailoring enzyme (*cloD*). Proposed biosynthetic pathway of clovibactin involves the assembly-line condensation of 8 canonical amino acids with 3 epimerizations carried out by dual-function condensation domains and a β-hydroxylation on Asn5 by the CloD, a TauD/TfdA dioxygenase. This hydroxylation provides the cyclization point for release from the NRPS and formation of the macrocyclic lactone.

The genome of *E. terrae ssp. carolina* was sequenced using PacBio. *E. terrae ssp. carolina* contains 19 predicted BGCs overall, of which 14 have NRPS-like elements (either pure NRPS or NRPS/PKS hybrid). The gene cluster associated with the biosynthesis of clovibactin was identified by antiSMASH (Hsu et al., 2004) version 5.1.1. It contains two NRPS genes (*cloA* and *cloB*), a transporter gene (*cloC*) and a tailoring enzyme (*cloD*). The proposed biosynthetic pathway of clovibactin is shown in **Figure 1**.

When comparing the clovibactin and teixobactin BGCs, the identity is 72% by BLASTN alignment **(Figure 1)**.

Clovibactin exhibited antibacterial activity against a broad range of Gram-positive pathogens, including methicillin-resistant *S. aureus* (MRSA), daptomycin-resistant as well as vancomycin-intermediate resistant (VISA) strains, and difficult to treat vancomycin-resistant *Enterococcus faecalis* (Lebreton et al., 2017) and *E. faecium* (VRE) **(Table 1, Supplementary Table 3)**. *Escherichia coli* was only marginally affected compared to an outer membrane deficient *E. coli* WO153 strain, probably reflecting insufficient penetration of the compound.

**Table 1.**
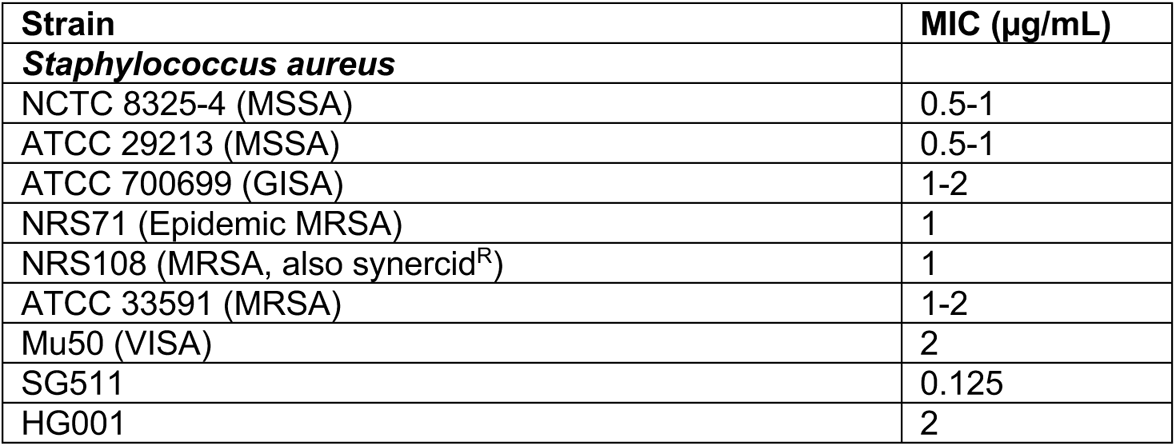
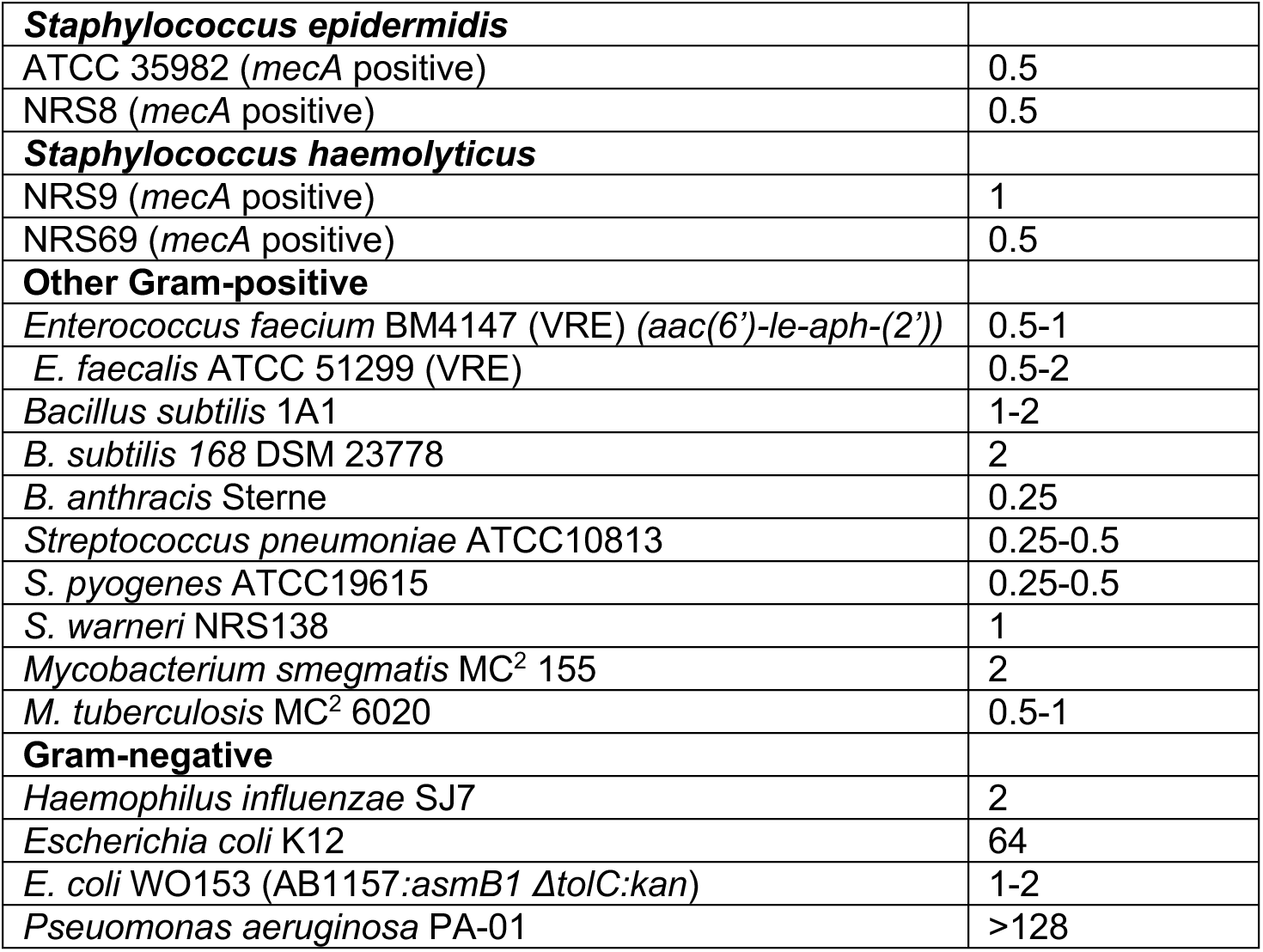
Antimicrobial activity of clovibactin. Minimal inhibitory concentrations (MIC) of clovibactin were determined by broth microdilution against selected strains and pathogenic bacteria. GISA, glycopeptide intermediate resistant *S. aureus*; MSSA, methicillin-sensitive *S. aureus*; MRSA, methicillin-resistant *S. aureus*; VISA, vancomycin intermediate resistant *S. aureus;* VRE, vancomycin-resistant enterococci.

Clovibactin is bactericidal, with *S. aureus* MBC (minimal bactericidal activity) of 2xMIC. We then examined the time-dependent killing in more detail **(Figure 2A)**. Clovibactin was more effective in killing *S. aureus* as compared to vancomycin, the first line of defense antibiotic. We noticed that clovibactin produced unusually strong lysis of the cell culture, and quantified this effect **(Figure 2B,C)**. Teixobactin was previously demonstrated to be rapidly bactericidal and to induce lysis mediated by AtlA (Ling et al., 2015), the major cell wall autolysin of *S. aureus* (Homma et al., 2016). To investigate the contribution of AtlA to the activity of clovibactin, the antibacterial activity against an Δ*atlA* deletion mutant was examined and compared to wild-type *S. aureus*. Clovibactin (2xMIC) induced strong lysis, more pronounced than teixobactin **(Figure 2B,C)**. Strikingly, lytic events were only slightly affected in the Δ*atlA* mutant, suggesting that clovibactin-induced lysis does not primarily rely on AtlA activity. Such pronounced lysis is typically observed with detergent-like compounds that rapidly destroy the cell membrane. However, lysis induced by clovibactin is not the result of rapid pore formation or membrane disruption, as evidenced by the absence of potassium efflux from staphylococcal cells in contrast to the pore-forming lantibiotic nisin **(Supplementary Figure 1)**. Similarly, clovibactin treatment did not lead to rapid membrane depolarization as determined using the membrane potential sensitive dye DisC2(5) or enable penetration of Sytox. This contrasts with the action of teixobactin that thins and depolarizes bacterial membranes (Shukla et al., 2022b). In agreement with this, clovibactin did not affect the cellular localization of the cell division protein MinD in *B. subtilis* **(Supplementary Figure 1)**. MinD is normally localized to cell poles and division sites and becomes delocalized upon dissipation of the membrane potential.

**Figure 2.**
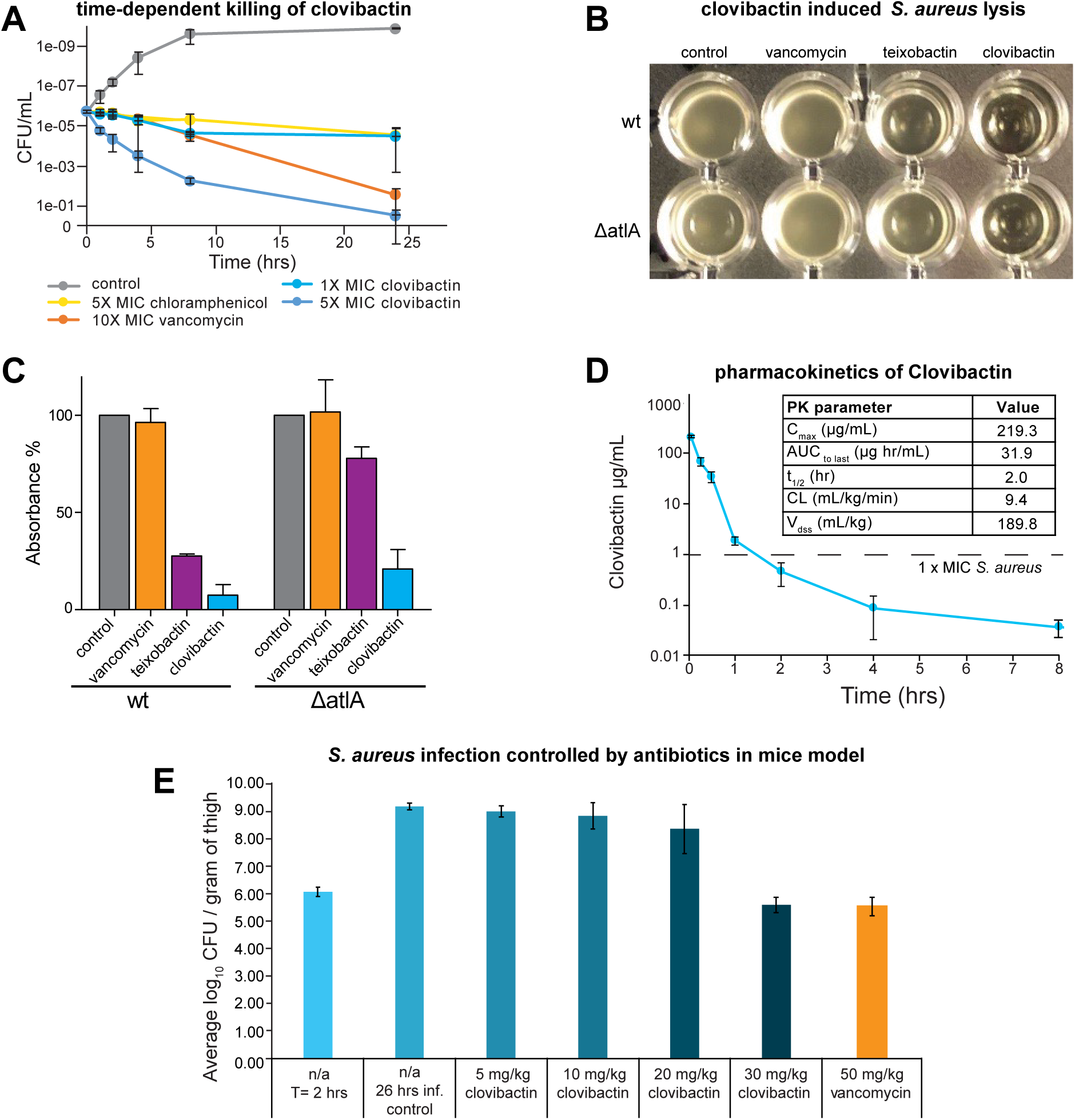
Clovibactin kills *S. aureus in vitro* and *in vivo*. **(A)** time-dependent killing of *S. aureus* (clovibactin (1xMIC), clovibactin (5xMIC), vancomycin (10xMIC) and chloramphenicol (5xMIC). **(B,C)** Clovibactin-induced lysis in *S. aureus*. Cells of *S. aureus* SA113 and a Δ*altA* mutant were incubated with each compound at 2xMIC for 24 hours as indicated. Mean values from three independent experiments are shown. Error bars represent standard deviation. **(D)** Pharmacokinetic parameters of clovibactin in mice model determined using Watson LIMS software. **(E)** The bacterial load from the thigh infection model prior to dosing and 24 hours after treatment. The infection controls demonstrated a bioload of 6.07 log10 CFUs/ gram of thigh at the time of treatment (2 hours). Clovibactin was delivered as two IV doses (2 and 4 hours post infection) of 5, 10, 20 and 30 mg/kg. Vancomycin was delivered as a single IV dose at 50 mg/kg.

Clovibactin did not show any cytotoxicity against mammalian NIH/3T3 and HepG2 cells at 100 µg/mL (highest concentration tested). Given the strong antimicrobial activity of clovibactin and low cytotoxicity, we next examined the action of this antibiotic *in vivo*. We first performed a pharmacokinetic study to evaluate the systemic exposure and blood residence time of the compound. A single dose of clovibactin was administered to mice at 20 mg/kg intravenously and was well tolerated. Blood was drawn at different times and clovibactin plasma levels were determined by LC-MSMS, and PK parameters were determined using the software package Watson LIMS **(Figure 2D)**.

Next, clovibactin was evaluated in a neutropenic mouse thigh infection model with *S. aureus*. In this model, mice are treated with cyclophosphamide to disable the immune response. Antibiotics are then evaluated for their ability to control infection without the assistance of the immune system. Clovibactin was comparable to vancomycin in diminishing the bacterial burden **(Figure 2E)**.

### Target identification

Given the novelty of the structure and the promising properties of this compound as a developmental lead, we sought to identify the molecular target of clovibactin. First, we determined the frequency of resistance, which is essential to know for advancing a compound. From a drug development standpoint, a desirable frequency of resistance is <10^-8^ (Silver, 2007) - low enough to provide extended lifetime in the clinic. The added benefit of this determination is target identification by whole-genome sequencing of resistant mutants. However, plating *S. aureus* on media containing clovibactin even at a low concentration (4xMIC) produced no resistant mutants. We therefore estimate the frequency of resistance to be <10^-10^. We then sought to determine a possible biosynthetic pathway that clovibactin might inhibit. For this, we followed the incorporation of labelled precursors into the major biosynthetic pathways of *S. aureus* – DNA, RNA, protein, and peptidoglycan. Clovibactin specifically interfered with the incorporation of radiolabeled GlcNAc into the cell wall, whereas DNA, RNA and protein biosynthesis remained unaffected **(Figure 3A)**. In a parallel approach, we used pathway-selective bioreporter strains of *Bacillus subtilis* treated with clovibactin (Harms et al., 2018). Expression of LacZ was specifically induced by clovibactin in *B. subtilis* P*_ypuA_*-*lacZ,* indicative of interference with cell wall biosynthesis **(Figure 3B)**. Treatment of *B. subtilis* with clovibactin induced cell-shape deformations as visualized by phase-contrast microscopy. This blebbing phenotype is characteristically induced by many cell-wall acting antibiotics and was similarly observed with teixobactin (Ling et al., 2015), hypeptin (Wirtz et al., 2021), or vancomycin, but not the protein synthesis inhibitor clindamycin used as a control **(Figure 3C)**, further supporting direct interference with cell wall biosynthesis. To narrow down the molecular target within the cell wall biosynthesis pathway, *liaI-lux* induction was monitored over time. LiaRS is a two-component system, known to respond to antibiotics that interfere with lipid II biosynthesis. Clovibactin strongly induced P*_liaI_-lux*, suggesting that it may directly interact with lipid II **(Figure 3D)**.

**Figure 3.**
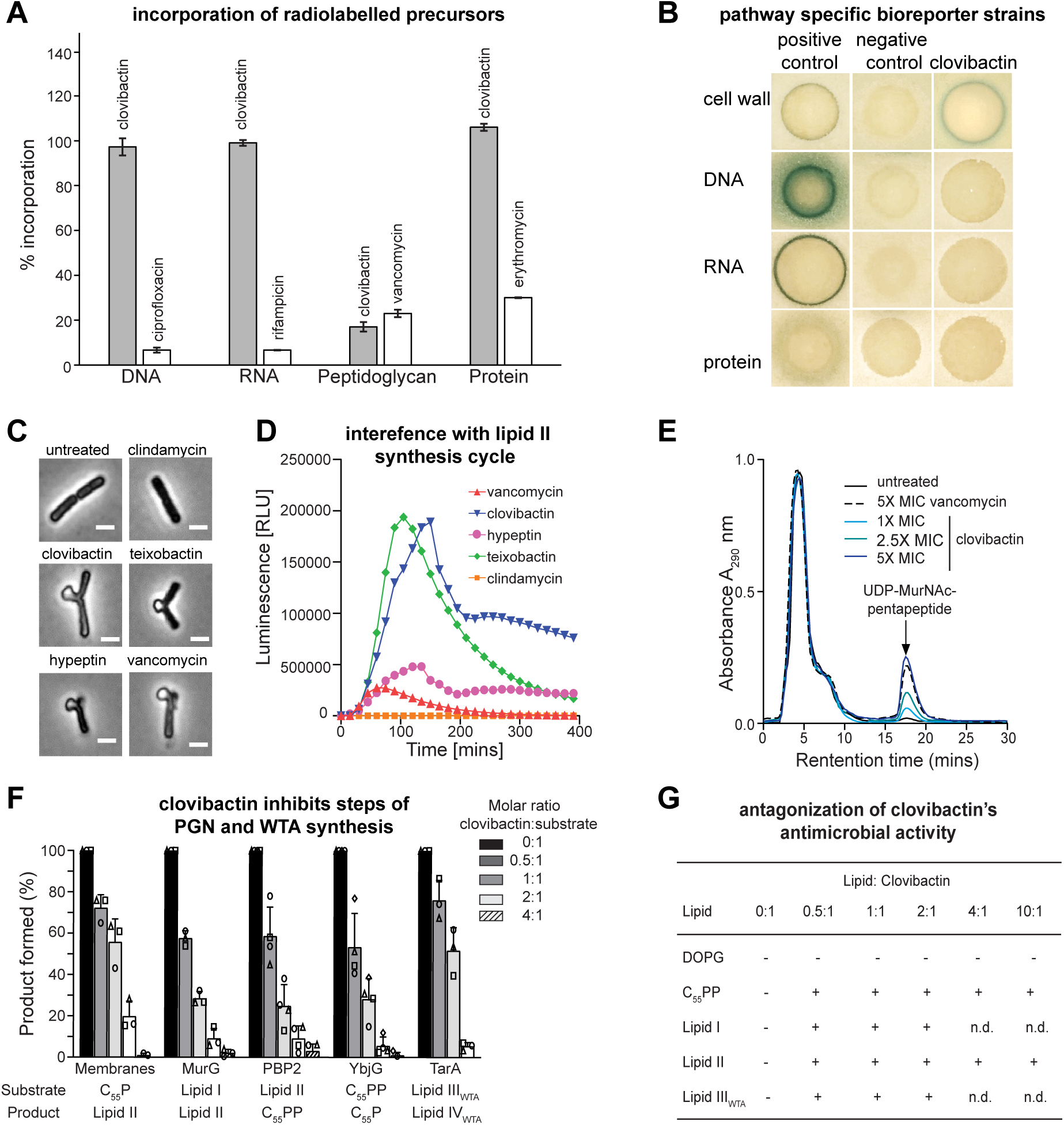
Clovibactin targets cell wall biosynthesis. **(A)** Effect of clovibactin on macromolecular biosyntheses in *S. aureus*. Incorporation of ^3^H-thymidine (DNA), ^3^H-uridine (RNA), ^3^H-leucine (protein), and ^3^H-glucosamine (peptidoglycan) was determined in cells treated with clovibactin at 2xMIC (grey bars). Ciprofloxacin (8xMIC), rifampicin (4xMIC), vancomycin (2xMIC) and erythromycin (2xMIC) were used as positive controls (white bars). Data are averages of two independent experiments. **(B)** *B. subtilis* bioreporter strains with selected promotor-*lacZ* fusions were used to identify interference with major biosynthesis pathways. β-galactosidase (*lacZ*) is fused to promotors P*_ypuA_* (cell wall), P*_yorB_* (DNA), P*_yvgS_* (RNA), and P*_yhel_* (protein) and induction of a specific stress response is visualized by a blue halo at the edge of the inhibition zone. Antibiotics vancomycin, ciprofloxacin, rifampicin, and clindamycin were used as positive controls. **(C)** Clovibactin treatment results in cell-shape deformations and characteristic blebbing as observed by phase-contrast microscopy of *B. subtilis*. Cell wall active antibiotics teixobactin, hypeptin, vancomycin and protein synthesis inhibitor clindamycin were used as controls. Scale bar = 2 µm. **(D)** Clovibactin (1xMIC, blue) strongly induced P*_lial_* as observed by expression of the *lux* operon in *B. subtilis* P*_liaI_*-*lux.* Teixobactin, hypeptin, vancomycin and clindamycin were used as control antibiotics. **(E)** Intracellular accumulation of the soluble cell wall precursor UDP-MurNAc-pentapeptide after treatment of *S. aureus* with different concentrations of clovibactin. Untreated and VAN-treated (5xMIC) cells were used as controls. Experiments are representatives of 3 independent experiments. **(F)** Clovibactin inhibits membrane-associated steps of PGN and WTA synthesis *in vitro*. The antibiotic was added in molar ratios from 0.5 to 4 with respect to the amount of the lipid substrate C_55_P, C_55_PP, lipid II, or lipid III_WTA_ used in the individual test systems. Reaction product synthesized in the absence of antibiotic was taken as 100%. Mean values from three independent experiments are shown. Error bars represent standard deviation. **(G)** Antagonization of the antimicrobial activity of clovibactin by cell wall precursors. *S. aureus* was incubated with clovibactin (8×MIC) in nutrient broth in microtiter plates, and growth was measured after a 24 h incubation at 37 °C. Putative HPLC-purified antagonists (undecaprenyl-pyrophosphate [C_55_PP], lipid I, lipid II, and lipid III_WTA_) and 1,2-dioleoyl-sn-glycero-3-phospho-glycerol (DOPG) were added in at molar ratios with respect to the antibiotic. Experiments were performed with biological replicates. +antagonization; - no antagonization.

Synthesis of the peptidoglycan precursor lipid II occurs in two different compartments of the bacterial cell. In the cytoplasm, the soluble sugar building blocks UDP-*N*-acetylmuramic acid pentapeptide (UDP-MurNAc-pp) and UDP-N-acetylglucosamine (UDP-GlcNAc) are synthesized and transferred to the lipid carrier undecaprenyl phosphate (C_55_P) to build lipid II that is flipped across the membrane to the exterior of the cell and incorporated into the growing peptidoglycan network **(Supplementary Figure 2)**. Antibiotics that block late stages of peptidoglycan biosynthesis, such as vancomycin, are known to trigger the intracellular accumulation of the last soluble peptidoglycan precursor UDP-MurNAc-pp. As observed for vancomycin, treatment of *S. aureus* with clovibactin at increasing concentrations led to an accumulation of UDP-MurNAc-pp in the cytoplasm **(Figure 3E)**. A fluorescent clovibactin-Bodipy-FL derivative bound preferentially to the septum of dividing staphylococcal cells, a site enriched in lipid II **(Supplementary Figure 3)**. Preincubation of cells with teixobactin almost completely blocked clovibactin-FL binding, suggesting that both compounds interact with the same targets **(Supplementary Figure 3)**.

Based on the results obtained with whole cells, the impact of clovibactin on the late steps of cell wall biosynthesis was analyzed *in vitro* to identify the molecular target – clovibactin was thus tested in individual biosynthesis assays using purified enzymes and substrates **(Figure 3F)**. Clovibactin inhibited all cell wall biosynthesis reactions that consume lipid I, lipid II, lipid III_WTA_ or undecaprenyl-pyrophosphate (C_55_PP) as a substrate in a dose-dependent fashion, suggesting binding to these lipid intermediates, rather than inhibiting enzyme function. Corroborating this result, the addition of purified cell wall lipid intermediates antagonized the antimicrobial activity of clovibactin and restored growth of *S. aureus* **(Figure 3G).** Interestingly, similar molar concentrations of C_55_PP and lipid II were required to fully antagonize clovibactin activity, while antagonization of teixobactin activity required a 10-fold higher concentration of C_55_PP compared to lipid II, suggesting differences in the binding mode. In agreement with this, higher concentrations of teixobactin (Ling et al., 2015) were needed to completely block the YbjG-catalyzed dephosphorylation compared to clovibactin **(Figure 3F)**.

### Oligomerization upon target binding

Next, we studied the interaction between clovibactin and lipid II in lipid bilayers using solid-state nuclear magnetic resonance (ssNMR), which allows the investigation of molecular mechanism of antibiotics binding to membrane targets under near-native conditions (Hong, 2006; Medeiros-Silva et al., 2018; Shukla et al., 2022b). To make the drug amenable to a comprehensive ssNMR characterization, we produced uniformly ^13^C,^15^N-labelled clovibactin by fermentation of *E. terrrae spp. carolina Δbat* in ^13^C,^15^N-enriched media (see Methods section). Co-assembly of clovibactin and lipid II in membranes resulted in high-quality 2D ssNMR correlation spectra that demonstrate the formation of a well-defined complex. We fully assigned the chemical shifts of clovibactin in the complex using 2D CC, CN, and NH spectra **(Supplementary Figure 4)**. Large signal shifts of the backbone amide protons show that clovibactin undergoes a major conformational change upon lipid II binding **(Figure 4A)**. An analysis of the ^13^C chemical shifts (Wang, 2002), suggests that the short linear N-terminus of clovibactin does not does not adopt a classical secondary structure but seems to contain elements of β-structuring. **(Supplementary Figure 4)**. Subsequently, we investigated the site-resolved dynamics of clovibactin in the lipid II bound state using ssNMR relaxation measurements (Lewandowski et al., 2011) **(Figure 4B,C)**. Globally, clovibactin is immobilised in the complex, indicative of the formation of a larger supramolecular structure. Strikingly, the N-terminus (Leu2 and Lys3) strongly rigidifies upon complex formation, which is reminiscent of the antimicrobial action of teixobactin, whose N-terminus drives the self-assembly into large clusters upon target binding (Shukla et al., 2022b; Shukla et al., 2020).

**Figure 4:**
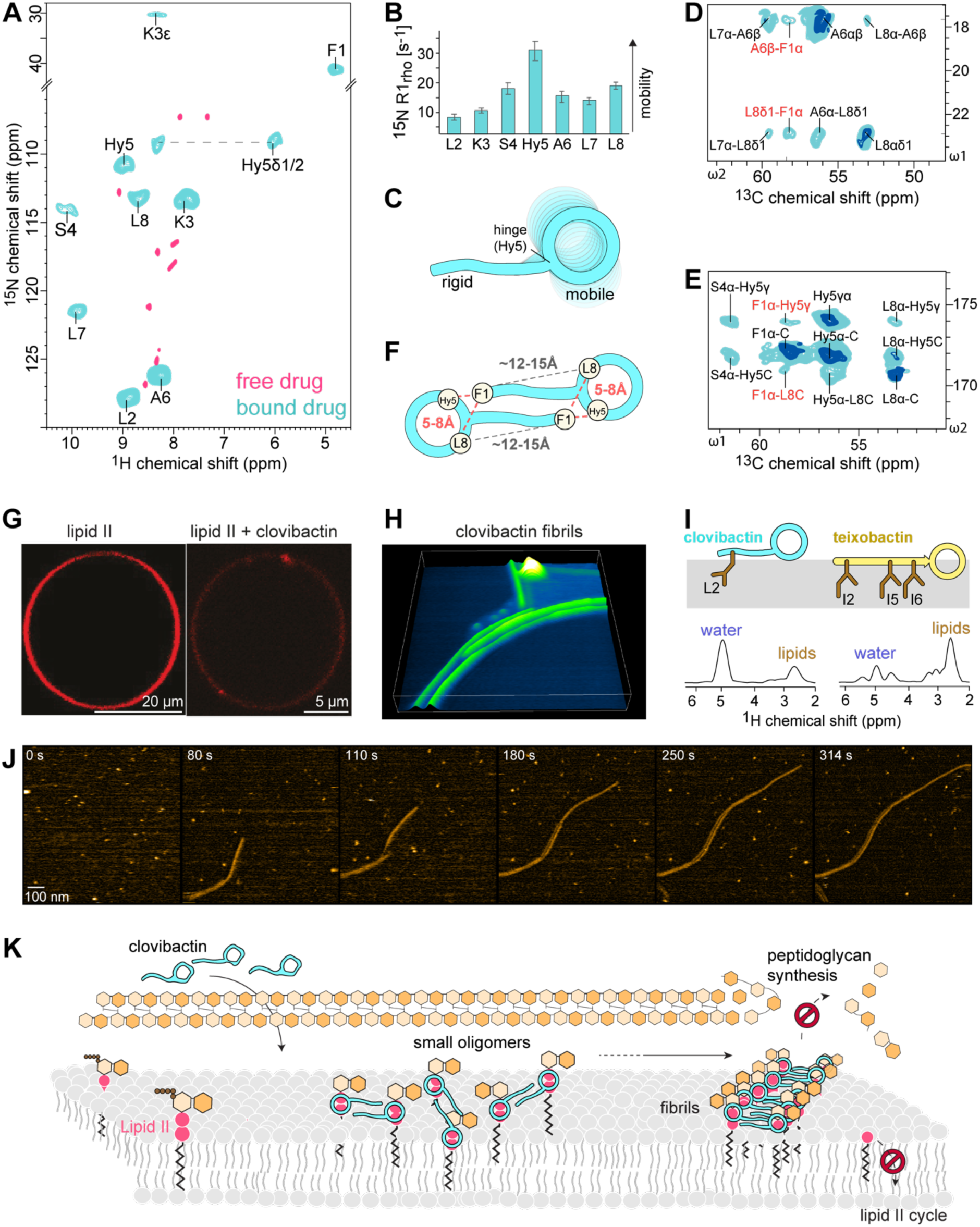
High-resolution ssNMR structure and oligomerization of the clovibactin/lipid II complex in membranes. **(A)** 2D NH ssNMR spectrum of lipid II bound clovibactin in membranes (cyan), superimposed on free clovibactin in aqueous solution (rose). **(B)** Site-resolved ^15^N R_1rho_ dynamics of lipid II bound teixobactin and clovibactin in DOPC membranes. **(C)** Illustration of the NMR-derived dynamics. While clovibactin’s N-terminus is rigid, the depsi-cycle shows elevated dynamics. **(D, E)** Zooms into a 2D CC spectrum of the complex show head-to-tail contacts in clovibactin, suggesting a dimeric (supramolecular) arrangement of clovibactin in the complex. Data acquired at 1200 MHz magnetic field using 50 (cyan) and 250 ms (dark blue) CC mixing time. **(F)** Schematic illustration of the head-to-tail contacts between Phe1-Hyn5 and Phe1-Leu8 seen in **(D, E)** a dimeric model. **(G)** Confocal microscopy of DOPC GUVs doped with Atto-labelled lipid II and treated with clovibactin show domain/cluster formation. **(H)** 3D rendered high-speed AFM image show the formation of clovibactin – lipid II supramolecular fibrils after 10 minutes of interactions. **(I)** left: Mobility-edited (Doherty and Hong, 2009) ssNMR experiments show that clovibactin is much more water-accessible than teixobactin in the lipid II-bound state. right: ssNMR-derived topology and membrane insertion of clovibactin. **(J)** Snapshots of a timelapse HS-AFM video (Supplementary Video 1) following the assembly of clovibactin–lipid II fibrils. Images were obtained on a supported lipid bilayer containing 4% (mol) lipid II in the presence of 5 µM clovibactin, added at 0 s. Image acquisition rate of 0.5 frames per second. **(K)** Model of the mode of action of clovibactin. At the membrane surface, clovibactin binds lipid II and forms small oligomers that serve as nuclei for the formation of fibrils. Fibril formation enables a stable binding of lipid II and other cell wall precursors, blocking cell wall biosynthesis.

We used confocal microscopy to probe the assembly of the complex on the surface of giant unilamellar vesicles (GUVs) doped with Atto-tagged lipid II. Microscopy images clearly show the formation of large clovibactin – lipid II patches, while such accumulation was not observed without clovibactin **(Figure 4G)**. These data confirm the formation of large supramolecular clovibactin – lipid II domains.

Next, we investigated at high-resolution how clovibactin molecules arrange in the supramolecular assemblies. We acquired 2D ssNMR PARISxy (Weingarth et al., 2010) CC spectra of the complex, in which we observed numerous clear head-to-tail contacts between N-terminal (Phe1) and C-terminal (Hyn5 and Leu8) clovibactin residues. In agreement with the formation of a supramolecular structure, these are likely intermolecular contacts between clovibactin molecules, as the intramolecular distances between N-terminal and C-terminal residues appear too far for CC magnetization transfer with a distance threshold of approximately 8 Å **(Figure 4D,E, Supplementary Figure 5)**. The contacts that we observe are consistent with an antiparallel dimeric organization of clovibactin molecules **(Figure 4F, Supplementary Figure 5)**.

### Flexible fibrils

To examine the supramolecular nature and the membrane interactions of clovibactin – lipid II complex in more detail, we first used high-speed atomic force microscopy (HS-AFM), a dynamic technique that provides biomolecular-scale structural resolution in real time (Kodera et al., 2010; Maity et al., 2020). Within minutes after the addition of 5 μM of clovibactin to membranes doped with lipid II, HS-AFM data show the formation of fibrils on the membrane surface **(Figure 4H, J and Supplementary Figure 6A)**. These fibrils only formed in the presence of both clovibactin and lipid II **(Supplementary Figure 6B)**. The observed fibrils showed a limited extend of lateral nucleation, forming thin, flat strings of associated fibrils.

The clovibactin concentrations required to observe supramolecular structures by HS-AFM requirements are comparable to the MIC, suggesting the oligomerization upon target binding is important or even critical for the mode of action of clovibactin **(Table 1 and Supplementary Table 3)**. The differential nature of the supramolecular structures of clovibactin and teixobactin (Shukla et al., 2022) is also in line with mobility measurements in supported bilayers using single-molecule tracking. These measurements agree with higher concentration requirements for the formation of clovibactin fibrils and show that lipid II remains more mobile in assemblies formed by clovibactin **(Supplementary Figure 8C)**.

To better understand the nature of clovibactin – lipid II assemblies, we next looked at its membrane topology at high-resolution using ssNMR experiments that monitor the exposure of the antibiotic to water and phospholipid phases (Doherty and Hong, 2009) **(Figure 4I and Supplementary Figure 7)**. ssNMR data show that lipid II-bound clovibactin localizes close to membrane/water interface and is in contact with both the water and lipid phases. Since the exposure to the water phase is much more pronounced, the data imply that the supra-structure formed by clovibactin lies on top of the membrane surface **(Figure 4K)**, sharply contrasting with teixobactin-supra structures, which show very strong interactions with the phospholipid tails and embeds deeply into the membrane. The differential membrane insertion can be rationalized by the presence of three long hydrophobic anchors (Ile2, Ile5, Ile6) in the N-terminus of teixobactin, while the much shorter N-terminus of clovibactin features only a single hydrophobic anchor (Leu2). The shallow membrane insertion of the clovibactin supramolecular structure is in agreement with the high localization of the fibrils above the membrane surface observed by HS-AFM (1.2 ± 0.2 nm above the membrane surface for clovibactin fibrils, 0.8 ± 0.1 nm above the membrane surface for teixobactin fibrils) **(Supplementary Figure 6A)**, and also appears in line with the increased mobility of lipid II in clovibactin assemblies **(Supplementary Fig 6C)**. Given that samples of the clovibactin complex yielded similar ssNMR spectra after weeks of storage at 278 K, the supra-structures apparently form irreversibly on biological timescales.

### The complex interface

Next, we sought to determine precisely how clovibactin targets lipid II, a complex lipid with a conserved pyrophosphate (PPi) group, a headgroup composed of the sugars MurNAc and GlcNAc and a pentapeptide whose variation confers resistance to antibiotics such as vancomycin (Münch and Sahl, 2015) **(Figure 5A)**.

**Figure 5:**
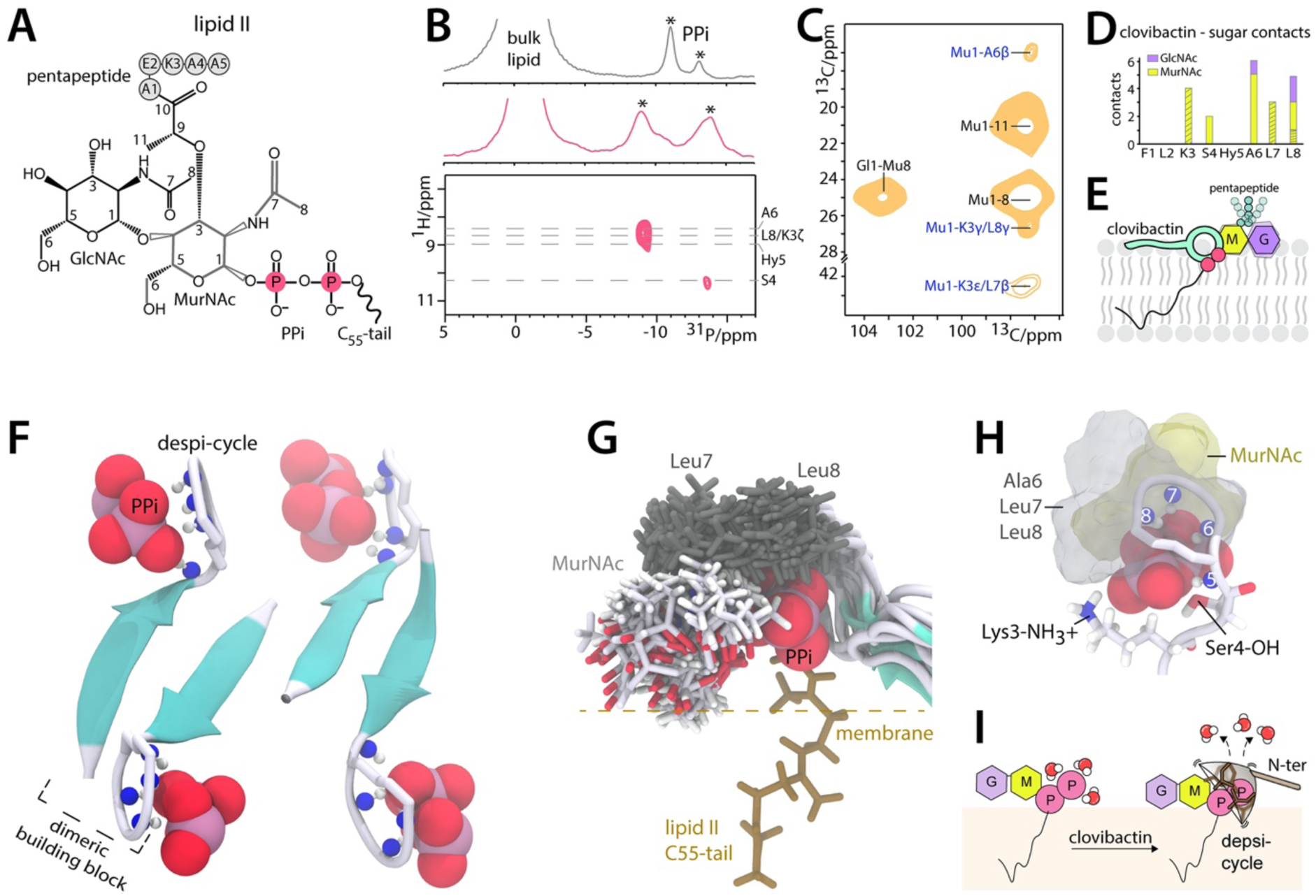
The interface and supramolecular structure. **(A)** Chemical structure of lipid II. **(B)** 1D ^31^P ssNMR data in liposomes show marked changes of the lipid II PPi signal upon addition of clovibactin. 2D ^1^H^31^P ssNMR spectrum establishes direct interactions between the backbone of the depsi-cycle and PPi. **(C)** 2D CC ssNMR data of the ^13^C,^15^N-clovibactin – ^13^C,^15^N-lipid II complex in liposomes show interfacial contacts with the MurNAc sugar and the hydrophobic sidechains of the depsi-cycle. Interfacial contacts in blue. Data acquired at 950 MHz using 300 ms CC mixing time. **(D)** Sum of interfacial contacts with the sugars of lipid II. Shaded bars show ambiguous contacts of MurNAc with either K3 or L7. **(E)** Illustration of the interface: PPi and MurNAc are in direct proximity, GlcNAc is distal, the pentapeptide is flexible and not involved in the interface. **(F)** ssNMR-derived structural model of the clovibactin – lipid II complex. **(G)** Calculated interfaces of ssNMR structures superimpose very well and show that the hydrophobic depsi-cycle sidechains (Ala6, Leu7, Leu8) wrap like a glove around lipid II-PPi group, interacting with the hydrophobic side of MurNAc. **(H)** The cationic K3 and the polar S4 sidechains favorably interact with the lipid II PPi group. **(I)** Hydrophobic residues of clovibactin embrace the PPi-group like a glove, which appears entropically favourable by the release of boundary water.

First, we investigated how clovibactin targets the PPi group. Upon addition of clovibactin, 1D ^31^P ssNMR data showed marked changes in the pyrophosphate signals of lipid II, suggesting a direct coordination **(Figure 5B and Supplementary Figure 8)**. A similar signal shift was observed for lipid I (which lacks the GlcNAc sugar). Furthermore, the emergence of intense sidebands in 1D ^31^P ssNMR data demonstrate that clovibactin also binds and immobilises C_55_PP (which lacks the sugars and the pentapeptide), in line with our biochemical binding analysis. Together, these data confirm that clovibactin is a multi-targeting antibiotic that blocks cell wall biosynthesis at several distinct, indispensable stages. To pinpoint the role of the lipid II PPi group in complex formation, we acquired a 2D ^31^P^1^H ssNMR spectrum which monitors magnetization transfer from amino protons of clovibactin to PPi **(Figure 5B)**. These data demonstrate that the backbone amino-protons of clovibactin directly coordinate the PPi group with the amino protons of the depsi-cycle (Hyn5, Ala6, Leu8). Furthermore, a weaker, but clearly discernible, signal shows that Ser4 of the N-terminus-depsi-cycle junction is in proximity close to the PPi group. These direct contacts with the PPi group agree with the stark signal changes observed upon addition of the drug.

Subsequently, to resolve the role of the lipid II sugars and the pentapeptide for target binding, we prepared a complex of ^13^C,^15^N-clovibactin and ^13^C,^15^N-lipid II to measure a series of 2D PARISxy (Weingarth et al., 2010) ^13^C^13^C spectra at ultra-high magnetic fields of 950 and 1200 MHz **(Figure 5C and Supplementary Figure 9A)**. We observed a total of 12 unambiguous interfacial contacts between clovibactin and the lipid II sugars, 10 of which relate to Ala6 and Leu8 **(Figure 5D)**, confirming that the depsi-cycle directly interacts with the lipid II headgroup. This is also in line with additional ambiguous interfacial contacts that we observed between Leu7 and the sugars. Together, our data show that all hydrophobic residues of the depsi-cycle are in direct proximity to the lipid II sugars. Almost all interfacial contacts are with MurNAc, the sugar that is covalently attached to the pyrophosphate, while we observed only three weak contacts with the GlcNAc sugar. This means that MurNAc is directly present at the interface with clovibactin, while the GlcNAc sugar is distal to this interface.

Strikingly, most residues of the lipid II pentapeptide were not observable in dipolar-based 2D CC spectra, in which we could only detect the first two residues (Ala*1 and *ψ*Glu*2). Since only rigid residues are observable in dipolar ssNMR spectra, our data strongly suggest that the pentapeptide is mobile and not part of the complex interface. This was confirmed with a complimentary scalar spectrum (Baldus and Meier, 1996), which gives a read-out on the mobile residues, in which we exclusively detected the last four residues (*ψ*Glu2*, Lys*3, Ala*4, Ala*5) of the pentapeptide **(Figure 5E and Supplementary Figure 10B)**.

### Structural model of the complex

We next calculated a structural model of the complex using HADDOCK (van Zundert et al., 2016). While the dimeric (2 x 2) clovibactin is likely the minimal binding arrangement to stably bind lipid II, we calculated a (4 x 4) arrangement to get a better idea about the supramolecular structure. The structure calculations were based on intermolecular clovibactin – clovibactin distance restraints and interfacial clovibactin – lipid II distance restraints. Hydrogen bonding restraints were applied among and between the dimeric units **(Supplementary Figure 11)**.

The obtained structures superimposed well (2.50±0.85 Å average backbone RMSD for clovibactin in the complex) **(Figure 5F and Supplementary Figure 12)** and show antiparallel dimeric units of clovibactin that could elongate to fibre-like supramolecular structures, in agreement with supramolecular structures observed by HS-AFM. A secondary structure analysis (Heinig and Frishman, 2004) shows about 60-70% β-strand propensity for clovibactin’s short N-terminus, an unusual molecular conformation that could foster oligomerization and fibril formation **(Supplementary Figure 12)**. Enabled by the sequence of alternating D- and L-amino acids, hydrophobic (Leu2) and hydrophilic (Lys3, Ser4) sidechains are divided below and above the sheet-like arrangement, positioned to face lipid and water phases, respectively **(Supplementary Figure 12).**

The clovibactin – lipid II interface is well defined by several unambiguous distance restraints (1.47±0.40 Å interfacial RMSD defined by residues Ala6, Leu7, Leu8 of clovibactin, PPi and MurNAc of lipid II) and shows that the backbone amino-protons of the depsi-cycle directly coordinate the PPi group of lipid II. Strikingly, the hydrophobic residues of the despi-cycle (Ala6, Leu7, Leu8) neatly wrap around the PPi-moiety, reminiscent of a hydrophobic glove structure, in agreement with ssNMR distance measurements **(Figure 5G)**. In this configuration, the hydrophobic depsi-cycle sidechains of clovibactin do not form specific interactions with the sugars. Instead, they form a plastic, adjustable interface with the hydrophobic side of the MurNAc sugar, in agreement with unambiguous contacts between Leu8δ1 – MurNAcC3, Ala6Cβ - MurNAcC11, or ambiguous contacts between Leu7Cβ - MurNAcC3 that are all within the hydrophobic patch of MurNAc. The lack of specific interactions with the MurNAc sugar also matches well with the increased dynamics of clovibactin’s depsi-cycle in the complex observed by ssNMR relaxation data, and the relatively weak interfacial clovibactin - sugar contacts observed in 2D CC ssNMR spectra. Residue 5 (3-hydroxyasparagine) that attaches the depsi-cycle to the linear N-terminus, shows by far the strongest dynamics, and hence acts as a sort of hinge that uncouples the dynamics of the PPi-binding depsi-cycle from the N-terminal oligomerization domain **(Figure 4B,C)**.

The structures consistently show favourable closer-distance interactions between the anionic lipid II PPi group and the cationic (Lys3) and the polar (Ser4) sidechains of the N-terminus **(Figure 5H)**. This agrees with clear interfacial ssNMR distance restraints Ser4Cβ - MurNAcC1 and Ser4Cβ - MurNAcC3 that show that Ser4-hydroxyl group is in proximity of the lipid II headgroup, and it matches well with the high rigidity of the Lys3 sidechain that is observable in dipolar 2D NH spectra **(Figure 4A and Supplementary Figure 4)**. 2D NH spectra also show that the protons of the cationic N-terminal amino-group of Phe1 are in fast exchange with water, which implies that they are not bound in a tight interaction.

The absence of specific interactions with the hydrophobic depsi-cycle sidechains (Ala6, Leu7, Leu8) is one of the reasons why clovibactin is able to efficiently (K_d_ for lipid II = 0.086 +/− 0.007 μM) bind a broad spectrum of cell wall precursors with a PPi group (C_55_PP, lipid I, lipid II) **(Figure 3E, 3F, and Supplementary Figure 13)**. The lack of specific interactions is likely compensated by favourable entropic contributions of the hydrophobic cage, which remains flexible but strips off boundary water molecules that coordinate the PPi group (**Figure 5I**). Critically, we did not observe binding of clovibactin to soluble PPi (Na-pyrophosphate) in isothermal titration calorimetry (ITC) experiments **(Supplementary Figure 13)**, demonstrating that clovibactin selectively binds lipid-anchored pyrophosphate groups. This suggests that settling on the membrane surface and the irreversible formation of supramolecular structures contribute to the effective binding of precursors.

## Discussion

Traditional screening platforms have failed to introduce new antibiotics in the last decades, in part due to overmining of Actinomycetes, the traditional source of antibiotics. Novel antibiotics are likely to be discovered by accessing silent operons, uncovering compounds masked by abundantly produced antimicrobials, and by growing previously uncultured bacteria (Cook and Wright, 2022; Lewis, 2020).

Clovibactin was isolated from an uncultured soil Gram-negative β-proteobacterium *E. terrae ssp. carolina*. It is a new cell wall acting antibiotic with an unusual structure and mode of action. The extract from the producing organism was active against *S. aureus* but isolating the major compound responsible for this activity led to kalimantacin, a known antibiotic (Fage et al., 2020). Kalimantacin is produced by *Pseudomonas* and *Alcaligenes* and inhibits the FabI enoyl-acyl carrier protein reductase of the fatty acid biosynthesis. Kalimantacin is inactive against *B. subtilis* carrying a different biosynthetic enzyme, but the extract was active against *B. subtilis* as well. Upon further examination of the extract, we isolated clovibactin, which was a relatively low abundant compound relative to kalamantacin. In order to simplify the isolation of clovibactin, we inactivated the kalimantacin operon. Inactivating the biosynthesis of abundant compounds to help detect and isolate minor compounds from extracts of interesting producers should be generally applicable to antibiotic discovery. In a similar approach, we recently reported isolation of a novel prodrug antibiotic aminodeoxyguanosine from a silent operon of *Photorhabdus luminescens* by first fractionating and concentrating an extract, and then screening (Shahsavari et al., 2022).

Clovibactin binds to the pyrophosphate moiety of multiple essential cell wall precursors C_55_PP, lipid II and lipid III_WTA_, from different cell wall biosynthetic pathways. Clovibactin binds to the PPi moiety of these precursors. In general, PPi seems as an unsuitable target for an antibiotic, since it will be released from dead cells together with PPi-containing nucleoside phosphates, and commonly present in the environment. At the same time, pyrophosphate is both an essential and immutable moiety of cell wall lipid intermediates, unlike other portions of the molecules, such as the pentapeptide of lipid II or sugars that can be modified in, e.g., mycobacteria (Mahapatra et al., 2005) or mutated. Moreover, a D-Ala-D-Lac substitution in the lipid II pentapeptide is a common mechanism of resistance to vancomycin that binds to the terminal D-Ala-D-Ala motif. Binding to immutable PPi would explain the lack of detectable resistance to this compound. How clovibactin manages to bind PPi of the central peptidoglycan precursor lipid II tightly *and* selectively is a fascinating question that we address in this study by performing a detailed structural analysis.

From the external medium, clovibactin settles at the membrane surface where it binds cell wall precursors. The leucine sidechain on the N-terminus helps partition clovibactin into the membrane at the site of its targets. Backbone amino-protons of clovibactin’s depsi-cycle directly coordinate the PPi, while the hydrophobic depsi-cycle sidechains (Ala6, Leu7, Leu8) surround the PPi group like an adjustable glove, an unusual interaction which is presumably entropically favorable by replacing boundary water, and which enables efficient binding to distinct, indispensable cell wall precursors. Attacking a highly polar target (PPi) with a hydrophobic warhead is a striking, counter-intuitive mode of action. Selective binding to the PPi moiety of lipid precursors is achieved by the formation of a supramolecular complex and subsequent oligomerization into an irreversible higher-order fibrillar assembly of interacting clovibactin molecules bound to the targets on the surface of the membrane. This is presumably enabled by an antiparallel arrangement of clovibactin molecules, in which the short N-terminus acts as oligomerization domain. The comparable concentrations for antibacterial action and for the formation of clovibactin fibrils suggests that supra-structures are an important or even essential part of the killing mechanism. The unique presence of readily accessible PPi-carrying molecules on bacterial membranes enables the formation of this structure and accounts for the lack of clovibactin toxicity against mammalian cells. A surprising feature of clovibactin is its superior ability to cause cell lysis. An intriguing possibility is that clovibactin fibrils, floating on top of the membrane, have an additional function of displacing autolysins from wall teichoic acids, resulting in lysis. The formation of an irreversible supramolecular structure will have favorable consequences for the in vivo activity of this compound, concentrating it onto bacterial surfaces where it belongs, and continuing to act long after the soluble compound has been cleared from the body. Clovibactin’s action against the PPi of cell wall precursors, a simple immutable target, expands our understanding of antibiotics evolved to avoid resistance, and points the way to rationally designing compounds with a long clinically useful life.

## Acknowledgements

This work was funded by the Netherlands Organisation for Scientific Research (NWO, grant numbers 723.014.003 & 711.018.001 to MW, and 718.015.001 to AMJJB). This project has received funding from the European Union’s Horizon Europe grant and innovation programme under grant agreement No. 101045485 (to MW). Funding for TS, KCL, FG, SDB was provided by the Deutsche Forschungsgemeinschaft (DFG, German Research Foundation), Project-ID 398967434 - TRR 261, and the German Center for Infection Research (DZIF). KL is supported by NIH grants P01AI118687 and RO1AI170962. Funding was also provided by NIH grant AI136137 to L.L.L., and NIH grant AI091224 to A.L.S. NMR experiments at the 950 and 1200 MHz instruments were supported by uNMR-NL, an NWO-funded Roadmap NMR Facility (no. 184.032.207). Support by Instruct-ERIC (to ML and MW) is acknowledged. This work has been supported BioExcel, grant numbers 675728 and 823830, funded by the Horizon 2020 program of the European Commission. The NRS strains were provided by the Network on Antimicrobial Resistance in *Staphylococcus aureus* (NARSA) for distribution by BEI Resources, NIAID, NIH. The authors also wish to acknowledge the help and expertise of the University of Maryland Core Genome Sequencing Facility, Micromyx LLC, Kalamazoo MI, NeoSome Life Sciences, Billerica MA. We would like to thank the following scientists for their work on clovibactin at NovoBiotic Pharmaceuticals, LLC Alysha Desrosiers, William Millett, Kelly Demeo, Ashley Zullo, and Cintia Felix. The plasmid pIJ12738 was a gift from the John Innes Centre.

## Author contributions

KL, TS, DH, LLL and MW designed the study. RS, AJP, MGN, AN, ML, and MW did NMR experiments. AN isolated clovibactin. KCL and SDB did mode of action studies. RS and FG did fluorescence microscopy. AMK and UK did single molecule mobility measurements in supported bilayers. SM and WHR did HS-AFM studies. ALS performed the fermentations, gene knockout work, and isolated the DNA for sequencing. CA performed susceptibility and cytotoxicity assays. CJS sequenced and annotated the genome. RS did calorimetric studies. RS, MGND, and EB prepared ssNMR samples and Lipid II. FL, BJAV, RVH, AMJJB, and MW did structure calculations. All authors contributed to data analysis and manuscript writing.

## Data availability

The NMR assignments of unbound clovibactin and the clovibactin – Lipid II complex have been deposited in the BMRB database (accession numbers 51629 and 51630). Experimental solid-state NMR raw data have been deposited in an open repository (DOI: 10.5281/zenodo.7075976).

## Material & Methods

### Isolation of strain producing clovibactin

The isolate producing clovibactin, P9846, was isolated from a sandy soil collected in North Carolina using techniques previously described (Buerger et al., 2012a, b). Briefly, 1 gram of heat-treated soil (dry heat at 65° C for 30 minutes) was mixed with 9 mL of sterile water, vortexed, and then diluted into molten SMS agar (0.125 g casein, 0.1 g potato starch, 1 g casamino acids, 20 g bacto-agar in 1 L of water). This mixture was then dispensed in 100 µL aliquots per well of a flat bottom 96-well plate. The 96-well plates were incubated for 16 weeks at room temperature in a humidified chamber and observations of growth were made over time. The isolate P9846 grew to a size detectable under a dissecting microscope (50X magnification) at week 12 of the incubation.

Isolates from long-term incubation experiments were sub-cultured from their original incubation plates to individual plates of SMSR4 (0.125 g casein, 0.1 g potato starch, 1.5 g casamino acids, 1 g glucose, 0.1 g yeast extract, 0.3 g proline, 1 g MgCl_2_-6H2O, 0.4 g CaCl_2_-2H_2_O, 0.02 g K_2_SO_4_, 0.56 g TES free acid (2-[[1,3-dihydroxy-2-(hydroxymethyl) propan-2-yl]amino]ethanesulfonic acid) and 20 g bacto-agar in 1 L of water agar). Monocultures of isolates were grown in a seed medium (15 g glucose, 10 g malt extract, 10 g soluble starch, 2.5 g yeast extract, 5 g casamino acids, and 0.2 g CaCl_2_-2H_2_O per 1 litre of deionized H_2_O, pH 7.0) to promote biomass production. Grown cultures were diluted 1:20 into 4 different fermentation broths. After 11 days of agitation at 28°C, the fermentations were dried and resuspended in an equal volume of 100% DMSO. Then 5 µL of extracts were spotted onto a lawn of growing *S. aureus* NCTC8325-4 cells in Mueller Hinton agar (MHA) plates. After 20 hours of incubation at 37°C, visible clearing zones indicated antibacterial activity. The extract from P9846 produced a large clearing zone. Although it produced antibacterial activity under several fermentation media, the best activity (that is, largest clearing zone) was seen with R4 fermentation broth (10 g glucose, 1 g yeast extract, 0.1 g casamino acids, 3 g proline, 10 g MgCl_2_-6H_2_O, 4 g CaCl_2_-2H_2_O, 0.2 g K_2_SO_4_, 5.6 g TES free acid (2-[[1,3-dihydroxy-2- (hydroxymethyl) propan-2-yl]amino]ethanesulfonic acid) per 1 litre of deionized H_2_O, pH adjusted to 7.0 using KOH).

### Clovibactin Isolation Procedure

n-Butanol (0.5V) was added to fermentation broth. The mixture was shaken vigorously, then left over night at room temperature until 2 distinct phases developed. The butanol phase was dried to completeness on a Buchi rotary evaporator. The resulting residue was reconstituted in a solution of 25% acetonitrile in water with 0.1% trifluoroacetic acid (~240 mL). This concentrated extract was centrifuged, and the supernatant decanted to remove any undissolved material. The supernatant was divided into 6 equal portions (~40 mL each) and successively purified on a preconditioned C18 flash chromatography column (Biotage, Sfar C18 60g, gradient of 25% - 100% acetonitrile in water (w/0.1% TFA), 50 mL/min). The compound of interest (clovibactin) was found at an adequate level of purity in two fractions as verified by mass spectrometry. The fractions containing clovibactin were concentrated under reduced pressure and lyophilized to yield approximately 125 mg from 6L of fermentation.

### Clovibactin Mass Spectrometry

Clovibactin was obtained as a white amorphous powder and its molecular formula (C_43_H_70_N_10_O_11_) was determined by LC-MS (*m/z* of 903.5291 [M+H]^+^, calculated 903.5298). This is consistent with ^1^H and ^13^C NMR data. Furthermore, ^13^C and ^15^N labeled clovibactin had an observed mass of 956.6472 [M+H]^+^ (calculated 956.6460).

### Clovibactin 1D and 2D solution NMR Experiments

About 10 mg of Clovibactin was dissolved in ~600 µL of DMSO-*d*6. The following NMR experiments were performed on this sample: ^1^H, ^13^C, ^1^H-^1^H COSY, ^1^H-^1^H COSY, ^1^H-^1^H NOESY, ^1^H-^1^H TOCSY, ^1^H-^13^C HSQC, ^1^H-^13^C HMBC, ^1^H-^15^N HSQC.

### Whole genome sequencing of P9846

Isolated gDNA was sequenced on a multiplexed PacBio Sequel II SMRT Cell 8M (Pacific Biosciences) using Sequel II 2.0 chemistry at the University of Maryland Core Genome Sequencing Facility yielding 836,139 total reads for this sample with a mean read length of 11,416 bp. Reads were assembled with SMRT8.0.0_HGAP4 (N50 = 5,455,962). The completed genome for P9846 contained a single circular chromosome and 3 extrachromosomal contigs (5.5 Mb chromosome, 7.0 Mb total) with 69% GC content.

### Biosynthetic analysis of P9846 genome

Assembled contigs were concatenated into a single nucleotide FASTA file and analyzed by AntiSMASH (https://antismash.secondarymetabolites.org/#!/start) version 5.1.1 using relaxed detection strictness and all extra features enabled. 10 putative biosynthetic gene clusters (BGCs) were identified on the chromosome, as well as 4 and 5 on two of the three extrachromosomal contigs for a total of 19 BGCs. Of these, two were found to have high similarity to known compounds in the Minimum Information about a Biosynthetic Gene cluster (MIBiG) database, the PKS compounds SGR-polycyclic tetramate macrolactam and malleilactone.

### Annotation of clovibactin biosynthetic gene cluster

The BGC responsible for cyclic peptide clovibactin was identified in the extrachromosomal genetic material where the majority of the NRPS BGCs are located. Eight complete NRPS modules corresponding to the amino acids found in the compound were identified and predictions of adenylation domain specificity from AntiSMASH were checked against Non-Ribosomal Peptide Synthase Substrate Predictor (NRPSsp, http://www.nrpssp.com/). Presence of dual-function condensation domains were used to identify sites of epimerization. Dioxygenase function was assigned based on homology to *cucE* from cupriachelin biosynthesis (MIBiG BGC0000330). The annotated sequence for clovibactin has been deposited into the MIBiG repository under the accession number #####.

### Determination of MICs

MIC values were determined using a broth microdilution method as recommended by the CLSI. The test medium for most species was cation-adjusted Mueller-Hinton broth (MHB). The same test medium was supplemented with 3% lysed horse blood (Cleveland Scientific, Bath, OH.) for growing Streptococci. *Haemophilus* Test Medium was used for *H. influenzae* (Teknova, Hollister, CA, or HTM; Remel; Lot No. 903401), Middlebrook 7H9 broth (Difco) was used for mycobacteria. For *N. gonorrhoeae*, a modified broth medium described by the ATCC was used. The medium contained 15 g Oxoid Special Peptone (Oxoid, Hampshire, UK; Lot No. 1280296), 1 g corn starch (Ward’s Science; Rochester, NY; Lot No. AD-13344-14), 5 g NaCl (VWR, Radnor, PA; Lot No. 57897), 4 g K_2_HPO_4_ (Sigma, St. Louis, MO; Lot No. 052K0147), and 1 g KH_2_PO_4_ (Sigma; Lot No. SLBC1921V). After autoclaving, the medium was centrifuged at 5,000 x g for 10 min, the supernatant was passed through a 0.45 µm filter, and IsoVitaleX supplement (BD; Lot No. 6309687) was added at 1% (v/v).

All test media were supplemented with 0.002% polysorbate (Tween) 80 to prevent drug binding to plastic surfaces, and cell concentration was adjusted to approximately 5X10^5^ cells/mL. After 20 hours of incubation at 37°C (2 days for *M. smegmatis,* 7 days for *M. tuberculosis*), the MIC was defined as the lowest concentration of antibiotic with no visible growth.

Fetal bovine serum (ATCC) was added to MHB (1:10) to test the effect of serum for *S. aureus* NCTC8325-4 and *S. aureus* ATCC29213.

Experiments were performed with three biological replicates.

### Mammalian cytotoxicity

The CellTiter 96® AQueous One Solution Cell Proliferation Assay (Promega) was used to determine the cytotoxicity of clovibactin on NIH 3T3 mouse embryonic fibroblast (ATCC CRL-1658, in Dulbecco’s Modified Eagle’s medium supplemented with 10% bovine calf serum), and HepG2 cells (ATCC HB-8065^TM^, in Dulbecco’s Modified Eagle’s medium supplemented with 10% fetal calf serum). Exponentially growing cells were seeded into a 96-well flat bottom plate, and incubated at 37°C. After 24 hours, the medium was replaced with fresh medium containing test compounds (0.5 µL of a two-fold serial dilution in DMSO to 99.5 µL of media, tested up to 100 µg/mL clovibactin). After 72 hours of incubation at 37°C, reporter solution was added to the cells and after 2 hours, the OD_490_ was measured using a Spectramax Plus Spectrophotometer. Experiments were performed with three biological replicates.

### Minimum bactericidal concentration (MBC)

Cells from the wells from an MIC microbroth plate for *S. aureus* NCTC8325-4 and *S. aureus* ATCC29213 that had been incubated for 20 hours at 37°C were pelleted. An aliquot of the initial inoculum for the MIC plate was similarly processed. The cells were resuspended in fresh media, plated onto MHA, and the colonies enumerated after incubating for 24 hours at 37°C. The MBC is defined as the first drug dilution which resulted in a 99.9% decrease from the initial bacterial titer of the starting inoculum. Experiments were performed with three biological replicates.

### Time-kill curves

Macrobroth MIC was determined for S. aureus ATCC 29213 in MHB II supplemented with 0.002% polysorbate 80 as 1 mL culture in polystyrene culture tubes after 20 hours of incubation at 37°C with aeration at 225 rpm. The MIC was 16 µg/mL for chloramphenicol, 1 µg/mL for vancomycin and 2 µg/mL for clovibactin under these conditions. To conduct time dependent killing, exponentially growing *S. aureus* ATCC 29213 cells were challenged with antibiotics at 37°C with aeration at 225 rpm - chloramphenicol (5X MIC, 80 µg/mL), vancomycin (10X MIC, 10 µg/mL), clovibactin (1XMIC, 2 µg/mL) or clovibactin (5X MIC, 10 µg/mL) in culture tubes at 37°C and 225 rpm. At intervals, 100 µL aliquots were removed, and 100 µL of ten-fold serially diluted suspensions were plated on MHA plates. Colonies were counted after 20 hours of incubation at 37°C, and CFU/mL was calculated. Experiments were performed with three biological replicates.

### Macromolecular synthesis assay

*S. aureus* NCTC8325-4 cells were cultured in minimal medium (0.02 M Hepes, 0.002 M MgSO_4_, 0.0001 M CaCl_2_, 0.4% succinic acid, 0.043 M NaCl, 0.5% (NH_4_)_2_ SO_4_) supplemented with 5% Tryptic Soy Broth (TSB). Cells were pelleted and resuspended into fresh minimal medium supplemented with 5% TSB containing test compounds and radioactive precursors to a density of 10^8^ cells/mL. The radioactive precursors were glucosamine hydrochloride, D- [6-3H(N)] (1 mCi/mL), leucine, L-[3,4,5-3H(N)] (1 mCi/mL), uridine, [5-3H] (1 mCi/mL), or thymidine, [methyl-3H] (0.25 mCi/mL) to measure cell wall, protein, RNA, and DNA synthesis, respectively. After 20 minutes of incubation at 37°C, aliquots were removed, added to ice cold 25% trichloroacetic acid (TCA), and filtered using Multiscreen Filter plates (Millipore Cat. MSDVN6B50). The filters were washed twice with ice cold 25% TCA, twice with ice cold water, dried and counted with scintillation fluid using Perkin Elmer MicroBeta TriLux Microplate Scintillation and Luminescence counter. Experiments were performed with two biological replicates.

### Resistance studies

No spontaneous resistant mutants of *S. aureus* ATCC 29213 were obtained when plating 1.2X10^10^ cells on agar media with 4xMIC clovibactin. No colonies grew up after 3 days of incubation at 37°C.

### Single dose PK in mice

Clovibactin was tested in a PK study to evaluate the systemic exposure and blood residence time of the compound. CD-1 female mice (n=3 per timepoint) were administered a single dose of compound at 20 mg/kg IV bolus. Blood was drawn at different times and clovibactin plasma levels determined by LC-MSMS. PK parameters were determined using the software package Watson LIMS.

### Neutropenic mouse thigh infection study

Clovibactin was evaluated in a neutropenic mouse thigh infection model. Female CD-1 mice (n=4 per group) received two doses of cyclophosphamide 4 days (150mg/kg), and 1 day (100 mg/kg) prior to infection. S. *aureus* ATCC 33591 (MRSA) was injected into the thighs. The infection controls demonstrated a bioload of 6.07 log_10_ CFUs/ gram of thigh at the time of treatment (2 hours). Clovibactin was delivered as two IV doses (2 and 4 hours post infection).

### Producing ^13^C-^15^N labeled clovibactin

P9846 makes multiple antibacterial compounds. Under the typical fermentation conditions (R4 broth) clovibactin is made in sufficient quantities. However, under labelling conditions we found that a known antibiotic, kalimantacin (Kamigiri et al., 1996), was produced in large amounts and interfered with clovibactin purification. Using the whole genome sequence, we identified the BGC with 55% identity to the kalimantacin gene(Mattheus et al., 2010). Production of kalimantacin was reduced below detectable levels by interrupting the bat genes and generating the strain P9846m01. The bat genes were interrupted using homologous recombination previously described (Fernández-Martínez and Bibb, 2014). Following procedures from Martinez et al. ~500 bp fragments from upstream and downstream of the P9846 bat1 gene were made by pcr using primers in supplemental table 2. Fragments were digested with XbaI or KpnI and BamHI (New England Biolabs) and cloned into plasmid pIJ12738 digested with XbaI and KpnI. This created a 4605 bp plasmid, pBat1ko. pBat1ko was transformed into mobilization strain ET12567. Using conjugation conditions, the pIJ12738 was transferred into P9846 and selected with 50 ug/ml apramycin on SMSR4 agar. The insertion in mutant, P9846m01 was confirmed using pcr with primers bat1downstream and bat1upstream (supplemental table 2). Disruption was further confirmed by fermentation in R4 medium and subsequent loss of kalimantacin derived active zones against a lawn of *S. aureus* followed by a lack of detectable kalimantacin by LC-MS in the appropriate fraction.

To label clovibactin, P9846m01 was grown from a frozen stock on SMSR4 agar supplemented with 50 ug/ml apramycin for 48 hours at room temperature. Biomass was scraped into 20ml flask of Celtone-R4 broth (10 g D-glucose; Cambridge Isotope Laboratories (CLM) #CLM-1396-5, 1 g Celtone base powder; CLM #CGM-1030P-CN-1, 0.5 g L-proline; CLM #CNLM-436-H-0.5, 10 g MgCl2-6H2O, 4 g CaCl2-2H2O, 0.2 g K2SO4, 5.6 g TES free acid (2-[[1,3-dihydroxy-2-(hydroxymethyl) propan-2-yl]amino]ethanesulfonic acid) per 1 liter of deionized H2O, pH 7 C). After 3 days of incubation with shaking at 28°C the culture was split between 2 flasks of 500ml Celtone-R4 broth. This culture was incubated with shaking at 28°C for 6 days. ^13^C and ^15^N labeled clovibactin was isolated in a similar manner as above, yielding 5.7 mg of ^13^C^15^N - labeled clovibactin from a 1 L fermentation.

### β-galactosidase reporter assays

*B. subtilis* β-galactosidase reporter assays were performed as previously described (Harms et al., 2018). In short, reporter strains were grown in MHB containing 5 μg/ml chloramphenicol at 30 °C to an OD_600_ of 0.5. Subsequently, cells were poured at 1 × 10^7^ CFU/ml in MHA plates supplemented with 5 μg/ml chloramphenicol, 75 µg/ml (cell wall reporter), 125 µg/ml (DNA reporter), and 250 µg/ml (RNA and protein reporters) X-gal, respectively. After solidification of the plates, 6 µg of clovibactin and control antibiotics selectively inducing the promotors were spotted (6 µg vancomycin for cell wall, 0.3 µg ciprofloxacin for DNA, 6 µg rifampicin for RNA, 3 µg clindamycin for protein). Results were documented after incubation overnight at 30°C.

### Quantification of intracellular UDP-*N*-acetylmuramic acid-pentapeptide

To analyze the cytoplasmic nucleotide pool we adapted the protocol of Kohlrausch and Höltje (Kohlrausch and Höltje, 1991). *S. aureus* SG511 was grown in 20 ml MHB at 37°C to an OD_600_ of 0.6 and incubated with 130 µg/ml chloramphenicol for 15 min. Clovibactin was added at 0.5×, 1×, 2.5×, and 5×MIC and incubated for another 30 min. Vancomycin (5× MIC) was used as positive control. Extraction and analysis of nucleotide-linked peptidoglycan precursors was performed as described previously (Schneider et al., 2009). Corresponding fractions were confirmed by mass spectrometry.

### Luciferase reporter antagonization assays

*B. subtilis* luciferase reporter assays were conducted as previously described (Tan et al., 2019; Umbreit and Strominger, 1972). Briefly, luciferase reporters were grown in MHB containing 5 μg/ml chloramphenicol at 30°C to an OD_600_ of 0.5. Cells were added to 96-well white wall chimney plates containing serially diluted antibiotics. For antagonization assays, purified cell wall lipid intermediates (C_55_PP, lipid I, lipid II, lipid III_WTA_) and phospholipids (phosphatidylglycerol [PG], phosphatidylcholine [PC], cardiolipin [CL]) were added in 0.5 to 4-fold molar excess with respect to clovibactin (0.5× MIC) and were pre-incubated for 10 min prior to addition of the reporter strain. Luminescence measurements were performed at 30 °C in a microplate reader Spark 10M (Tecan). At least three independent biological replicate experiments were conducted.

### Bacterial cell wall integrity assay

Bacterial cell wall integrity assays were adapted from previous work (Schneider et al., 2010). *B. subtilis* 168 cultures were grown in MHB at 30°C to an OD_600_ of 0.3. Subsequently, cells were treated with 0.5 µg/ml clovibactin, 1 µg/ml vancomycin, 32 µg/ml clindamycin, or DMSO for 30 min. Cells were immediately fixed in a 1 ml 1:3 (v:v) mixture of acetic acid and methanol and immobilized on thin 1% w/v agarose slides. Imaging was performed by phase contrast microscopy on a Zeiss AxioObserver Z1 equipped with an HXP 120 C lamp, an αPlan-APOCHROMAT 100×/1.46 oil objective and an AxioCam MRm camera and further processed with Zen 2 (Zeiss) and analyzed and postprocessed using ImageJ v1.52p(Schneider et al., 2012).

### MIC-type antagonization assays

Antagonization of the antibiotic activity of clovibactin by potential target molecules was performed by an MIC-based setup in microtiter plates. Clovibactin (8× MIC) was mixed with HPLC-purified antagonists (C_55_PP, lipid I, lipid II, lipid III_WTA_ and DOPG) in 0.5 to 4-fold molar excess with respect to the antibiotic. *S. aureus* SG511 (5 × 10^5^ CFU/ml) was added and samples were examined for visible bacterial growth after incubation at 37°C overnight. Experiments were performed with at least three biological replicates.

### Synthesis and purification of lipid intermediates

Large scale synthesis and purification of the PGN precursors lipid I and lipid II were performed as previously described (Ling et al., 2015). UDP-*N*-acetyl-muramic acid pentapeptide (UDP-MurNAc-pp) was purified according to the protocol elaborated by Kohlrausch and Höltje (Kohlrausch and Höltje, 1991). Undecaprenyl phosphate (C_55_P) and undecaprenyl diphosphate (C_55_PP) were purchased from Larodan Fine Chemicals AB (Malmö, Sweden). The phospholipids 1,2-dioleoyl-*sn*-glycero-3-phosphocholine (DOPC), 1,2-dioleoyl-*sn*-glycero-3-phosphoglycerol (DOPG) and 1′,3′-bis[1,2-distearoyl-*sn*-glycero-3-phospho]-glycerol (DOCL, cardiolipin) were purchased from Avanti Polar Lipids (Alabaster, AL, USA). The concentration of purified PGN and wall teichoic acid precursors was quantified on the basis of their phosphate content as described (Rouser et al., 1970).

### *In vitro* lipid II synthesis with isolated membranes

*In vitro* lipid II synthesis was performed using membranes of *M. luteus* as previously described (Brötz et al., 1998) (Umbreit and Strominger, 1972). Briefly, synthesis was assayed by incubating membrane preparations (200 µg protein) with 5 nmol C_55_P, 50 nmol UDP-MurNAc-pentapeptide, 50 nmol UDP-*N*-acetylglucosamine (UDP-GlcNAc) in 60 mM Tris-HCl, 5 mM MgCl_2_, and 0.5 % Triton X-100, at pH 7.5 in a total volume of 50 µl at 30°C for 1 h. C_55_P-containing products were extracted with an equal volume of *n*-butanol/pyridine acetate, pH 4.2 (2:1, v/v) and analyzed by TLC using chloroform/methanol/water/ammonia (88:48:10:1, v/v/v/v) as the solvent(Rick et al., 1998) and phosphomolybdic acid staining (Schneider et al., 2004). The quantitative analysis of lipids extracted to the butanol phase was carried out by phosphorimaging in a StormTM imaging system (GE Healthcare) or PMA staining and analysis performed using Image Quant TL. Clovibactin was added in molar ratios of 0.5 to 4 with regard to C_55_P.

### *In vitro* PGN synthesis reactions using purified proteins and substrates

To determine the enzymatic activity of MraY-His_6_ the assay was carried out in a total volume of 50 µl containing 5 nmol C_55_P, 25 nmol of UDP-MurNAc-pp in 100 mM Tris-HCl, 10 mM MgCl_2_, at pH 7.5, and 0.6% Triton X-100. The reaction was initiated by the addition of 2 µg of MraY-His_6_ and incubated for 1.5 hours at 30°C.

The *in vitro* MurG reaction was performed in a 30 μL reaction containing 2 nmol of purified lipid I and 25 nmol of UDP-*N*-acetyl glucosamine (UDP-GlcNAc) in 200 mM Tris-HCl, 5.7 mM MgCl_2_, at pH 7.5, and 0.8% Triton X-100 with 2 μg of purified, recombinant MurG-His_6_ enzyme. Reaction mixtures were incubated for 30 min at 30°C.

The PBP2 activity assay was performed in a 50 μL reaction containing 2 nmol of purified lipid II, in 20 mM MES, 2 mM MgCl_2_, and 2 mM CaCl_2_, at pH 5.5 with 2 μg of purified, recombinant PBP2-His_6_ enzyme. Reaction mixtures were incubated for 2 h at 30 °C. Dephosphorylation of C_55_PP was determined using purified YbjG-His_6_ enzyme (SA0415). A total of 20 nmol of C_55_PP was incubated with 3 μg of YbjG-His_6_ in 20 mM Tris-HCl, 150 mM NaCl, and 0.8% Triton X-100 at pH 7.5 in 50 μL for 30 min at 30°C.

In all *in vitro* assays, clovibactin was added in molar ratios from 0.5 to 2 (and to 4 for MraY and MurG) with respect to the respective substrate. C_55_P-containing products were extracted, analyzed by TLC and quantified as described above. The quantitative analysis of lipids extracted to the *n*-butanol phase was carried out using ImageJ v1.53p software (National Institutes of Health) (Schneider et al., 2012). Experiments were performed at least in triplicates.

### Complex formation clovibactin

Binding of clovibactin to C_55_PP, lipid I, lipid II, lipid III_WTA_ and DOPG was analyzed by incubating 2 nmol (lipid I, lipid II, lipid III_WTA_) or 5 nmol (C_55_PP and DOPG) with 8-fold molar excess of clovibactin in 50 mM Tris-HCl, at pH 7.5 for 30 min at room temperature. Complex formation was analyzed by extracting unbound precursors from the reaction mixture followed by TLC analysis as described above. Experiments were performed with biological replicates.

### Lysis assay

Visual analysis of lysis was conducted in microtiter plates, exponential and stationary phase cells (OD_600_ 0.5 and 1, respectively) cells of *S. aureus* SA113 and the AtlA-deficient mutant *S. aureus* SA113 Δ*altA* (supplemented with 150 µg/ml spectinomycin) were treated with clovibactin and vancomycin at concentrations of 0.5×, 1× and 2× the MIC and were photographed after 24 hours. Experiments were performed with three biological replicates.

### Bacterial cell wall integrity “blebbing” *assay*

Bacterial cell wall integrity assays were performed as described previously (Schneider et al., 2010). *B. subtilis* 168 cultures were grown in MHB at 30°C to an OD_600_ of 0.3. Subsequently, cells were treated with clovibactin, teixobactin, hypeptin and vancomycin (0.25 x MIC each), 256 µg/ml ampicillin, and further incubated at 30°C for 90 min. Cells were immediately fixed with in a 1 ml 1:3 (v:v) mixture of acetic acid and methanol and immobilized on thin 1 % w/v agarose slides. Imaging was performed by phase contrast microscopy on a Zeiss Axio Observer Z1 microscope (Zeiss, Jena, Germany) equipped with HXP 120 V light source and an Axio Cam MR3 camera. Images were acquired with ZEN 2 software (Zeiss) and analyzed and postprocessed using ImageJ v1.45s software (National Institutes of Health).

### Potassium efflux

The potassium release assay was performed as described previously. Briefly, tryptic soy broth (TSB)-grown *S. simulans* 22 cells were harvested at an OD_600_ of 1.0 to 1.5, washed with cold choline buffer (300 mM choline chloride, 30 mM MES, 20 mM Tris, pH 6.5), and resuspended to an OD_600_ of 30. The concentrated cell suspension was kept on ice and used within 30 min. For each measurement, the potassium electrode (Mettler Toledo) was calibrated with potassium chloride. Cells were diluted in choline buffer at RT to an OD_600_ of 3, and antibiotic-induced potassium release was monitored in 15 sec intervals for 5 min at RT. Clovibactin was added in a concentration corresponding to 1 and 5× MIC. Potassium concentrations were calculated from the measured voltage according to (Orlov et al., 2002) and plotted relative to the total amount of potassium released after the addition of 1 µM of the pore-forming lantibiotic nisin (set 100 % efflux). Results show mean values of three independent experiments.

### MinD delocalization studies

*B. subtilis* 1981 *erm spc minD*:*ermC amyE*::P*_xyl_*-*gfp*-*minD*, a strain with a *gfp*-*minD* fusion under control of the P*_xyl_* promotor, was grown in MHB supplemented with 0.1 % w/v xylose and 50 µg/ml spectinomycin at 30 °C to an OD_600_ of 0.6. Imaging was carried out within 2, 5, and 30 min after addition of antibiotic at 2× and 10× MIC. The proton ionophore carbonyl cyanide *m*-chlorophenylhydrazone (CCCP, 100 µM) was used as positive control and imaging was carried out within 2 min. Samples were immobilized on microscope slides covered with 1 % w/v agarose. Fluorescence microscopy and analysis was performed using the same microscope and software as described for phase contrast microscopy.

### Fluorescence labeling and fluorescence microscopy

Clovibactin was dissolved in anhydrous DMF (5 mg/ml) and BODIPY™ FL succimidyl ester (Invitrogen) was dissolved in DMSO (10 mg/ml). For labeling, 1 vol of BODIPY™ FL was added to 0.5 vols of DIPEA and 5 vols clovibactin solution and incubated at room temperature with gentle agitation. After 2 hours, ice-cold diethylether was added to the reaction mixture leading to precipitation of clovibactin. After centrifugation, the supernatant was carefully aspirated, and the pellet was dissolved in anhydrous DMF. Separation of labeled and non-labeled clovibactin was achieved by HPLC purification on a Poroshell 120 EC-C18, 2.7 µm, 3 × 150 mm column (Agilent), column temperature 50 °C, with gradient of (A) 79.9% H_2_O + 20% acetonitrile (MeCN) + 0.1% trifluoroacetic (TFA) and (B) 49,9% H_2_O + 50% MeCN + 0.1% TFA: 0 min, %B = 0, 10 min, %B = 60, 34 min, %B = 80. Clovibactin-FL was collected and confirmed by LC-MS.

To study localization of clovibactin, cultures were grown in MHB at 37°C until an OD_600_ of 0.5 and were pre-incubated for 10 min with teixobactin (2× MIC) or the corresponding amount of DMSO. Cells were then washed four times with MHB prior to 10 min incubation with a 1:3 mixture of unlabeled and labelled clovibactin (2× MIC) and Tween 80 at a final concentration of 0.002%. To visualize cell membranes, nile red was added to a final concentration of 1 µg/ ml. Subsequently, cells were washed four times in phosphate-buffered saline (PBS) and mounted on microscope slides covered with a thin film of 1% v/v agarose. Images were acquired using a Zeiss AxioObserver Z1 equipped with an HXP 120 C lamp, an αPlan-APOCHROMAT 100×/1.46 oil objective and an AxioCam MRm camera. Standard filter sets were used for BODIPY™ FL (450–490 nm excitation and 500–500 nm emission). Images were further processed with Zen 2 (Zeiss) and analyzed and postprocessed using ImageJ v1.52p (Schneider et al., 2012).

### Analysis of single molecule mobility in supported lipid bilayers

Lipid II-Atto565 was obtained by labeling of lipid II with a 10-fold excess of Atto565-NHS ester (Sigma-Aldrich, Taufkirchen, Germany) in presence of 0.6 vol% diisopropylethylamine (Sigma-Aldrich, Taufkirchen, Germany). The reaction was carried out in water-free chloroform for 2 hours at room temperature. Lipid II-Atto565 was purified via thin layer chromatography.Supported bilayers were formed by DOPC 0.2 mol% lipid II/lipid II-Atto565. The required lipid composition was prepared in chloroform to reach a lipid concentration of 1.3 mM. Then, chloroform was removed in a nitrogen stream and the dried lipid film was solved in HEPES/Triton buffer (20 mM HEPES, 150 mM NaCl, 20 mM Triton X-100, pH 7,4) in order to reach a lipid concentration of 5 mM. The stock solution was divided into 20 µL aliquots and stored at −20°C. To enable the preparation of planar lipid bilayers on coverslips, a suspension containing very small unilamellar vesicles (VSUV) was prepared by addition of 200 µL HEPES buffer (20 mM HEPES, 150 mM NaCl) and 200 µL of a 4 mM heptakis(2,6-di-O-methyl)-ß-cyclodextrin to each of the aliquoted lipid-detergent solutions. Finally, the vial was vortexed for 2 minutes.

In order to increase the hydrophilicity of the glass support, coverslips with a diameter of 18 mm were placed for at least 1 hour in freshly prepared Piranha-solution (one-part H_2_O_2_ 30% and three parts H_2_SO_4_). Then, the coverslips were washed with milliQ water, dried in a nitrogen stream and inserted into custom-built sample chambers. The VSUV suspension resulting from one aliquot was immediately added on top of the cleaned coverslip. After an incubation time of 6 minutes, unfused vesicles were removed by washing of the coverslip with HEPES buffer. During the washing steps care was taken to not dry out the lipid bilayer. Bilayers were kept in HEPES buffer and the respective amounts of teixobactin and clovibactin were added from stock solutions in DMSO to freshly prepared bilayers.

After an incubation time of 30 minutes the samples were imaged using a custom-built setup enabling single-molecule microscopy (Ruland et al., 2021). An α-plan-apochromat 63x/1.46 objective (Zeiss) and a sCMOS camera (Prime BSI, Teledyne Photometrics, Tucson, AZ, USA) were used for data acquisition resulting in a pixel size of 103.2 nm. Images were acquired with a field of view of 30.96 µm x 30.96 µm and a single frame integration time of 10 ms. Excitation of Atto565 was performed using a DPL 561 nm laser (Hübner Photonics GmbH, Kassel, Germany). For every sample 27 movies consisting of 1000 frames were acquired at different sample locations. For every experimental condition three independently prepared samples were measured. Single-particle tracking was performed with the ImageJ 1.52p (Schindelin et al., 2012) (Schindelin et al., 2012) plugin TrackMate (Tinevez et al., 2017) after background subtraction using a rolling ball radius of 50. For spot detection, the LoG-based detector was used, the parameter “estimated blob diameter” was set to 0.75 µm, the threshold was set to 2.2. Tracking was performed by the “Simple LAP Tracker”. Gap-closing was allowed with a maximum closing distance of 1 µm and a maximum frame gap of two frames. A maximum linking distance of 1 µm was chosen. From the resulting tracking data, the mean square displacement as function of time was determined and plotted with the Software OriginPro 2021b (OriginLab Co., Northhampton, MA, USA). A linear fit was performed and according to the Einstein equation for two-dimensional diffusion, the diffusion coefficient of lipid II-Atto565 molecules was determined by dividing the slope of the linear fit by 4.

### Synthesis and purification of isotopically labelled Lipid II

Lipid II was produced according to published methods(Breukink et al., 2003) based on enzymatic lipid reconstitution using the Lipid II precursors UDP-GlcNAc, UDP-MurNAc-pentapeptide and polyisoprenolphosphate as substrates.(Breukink et al., 2003) Lysine-form UDP-MurNAc-pentapeptide was extracted from *Staphylococcus simulans* 22. ^13^C,^15^N-labelled UDP-GlcNAc and UDP-MurNAc-pentapeptide (lysine form) were extracted from *S. simulans* 22 grown in [^13^C/^15^N]-labelled rich medium (Silantes) and supplemented with [U-^13^C]-D-glucose and [^15^N]-NH_4_Cl. Polyisoprenolphosphate was synthesized via phosphorylation of polyisoprenol obtained from *Laurus nobilis*(Danilov et al., 1989). The head-group precursors were extracted from bacteria and polyisoprenol was extracted from leaves as described (Kohlrausch, 1991). After synthesis, Lipid II was extracted with 2:1 BuOH:(Pyr/Acetate, 6 M) and then purified with a DEAE cellulose resin using a salt gradient of 0 - 600 mM NH_4_HCO_3_ with 2:3:1 CHCl_3_:MeOH:[H_2_O+salt]. Fractions containing pure Lipid II were pooled, dried, and dissolved in 2:1 chloroform/methanol. Lipid II concentration was estimated through an inorganic phosphate determination(Rouser et al., 1970).

### Solid-State NMR sample preparation

Multi-lamellar vesicles (MLVs) of DOPC doped with 4 mol% Lysine-Lipid II in buffer (30mM Citrate, 300mM NaCl, pH 5.5) were collected by centrifugation (60,000×*g*) and loaded into ssNMR rotors. For 3.2 mm rotors, we used 800 nmol of clovibactin with unlabelled Lipid II, while we used 400 nmol with labelled Lipid II. For 1.3 mm rotors, samples contained 200 nmol of antibiotic for unlabelled Lipid II.

### Solid-state NMR spectroscopy

^1^H-detected ssNMR experiments were performed at 60 kHz magic angle spinning (MAS) using magnetic fields of 700, 950, and 1200 MHz (^1^H frequency) with dipolar transfer steps and using low-power PISSARRO (Weingarth et al., 2009a) decoupling in all dimensions. ^1^H-detected ^15^N T_1rho_ relaxation experiments (Lewandowski et al., 2011) were acquired with a ^15^N spin lock-field of 18 kHz and spin-lock durations of 0, 10, 20, 40, 70, and 100 ms. T_1rho_ trajectories were fit to single exponentials. 2D CC experiments were acquired with PARIS (Weingarth et al., 2009b) PARISxy (Weingarth et al., 2010) recoupling (m=1) at 950 and 1200 MHz magnetic field and 15 to 18 kHz MAS. To probe interfacial contacts between ^13^C,^15^N-clovibactin and ^13^C,^15^N-Lipid II, we used CC magnetization transfer times of 50, 150, and 300 ms. 2D CaN and CON experiment were acquired at 800 MHz, 15 kHz MAS, and 5 to 7 ms N to C cross-polarization transfer time. To characterize Lipid II-bound teixobactin, we used CC magnetization transfer times of 50 and 300 ms. The scalar 2D CC TOBSY (Baldus and Meier, 1996) experiment was acquired at 700 MHz using 8 kHz MAS with 6 ms CC mixing time. The mobility edited(Doherty and Hong, 2009) H(H)C experiment was measured at 1200 MHz with 16.5 kHz MAS at 300 K temperature using a T_2_ relaxation filter of 2.5 ms. 1D MAS ^31^P experiments were acquired at 500 MHz magnetic field and 12 kHz MAS. 2D HP experiments were acquired at 800 MHz and 60 kHz MAS using 1 and 2 ms ^1^H to ^31^P cross-polarization contact time.

### Fluorescence Microscopy

*GUVs preparation*: We used a self-assembled GUV cell, aligned with two titanium electrodes in a closed Teflon chamber (volume = 500 mL). 1 mL of 0.5 mM DOPC doped with Atto 550-labelled Lipid II (0.1 mol%) was brushed on the titanium electrodes. The GUV cell was dried under vacuum. Next, the chamber was filled with 350 mL 0.1M sucrose solution, the electrodes dipped in and connected to a power supply of a sine wave (2.5V; 10 Hz; 90 minutes). Each microscopy slide (m-slide 8 well, Ibidi) was incubated with 350 mL BSA solution (1 mg/mL) for 1 hour. To detach the GUVs, the power supply was changed to square wave (2V; 2Hz; 15 minutes). The slides were washed once with water and 0.1M glucose solution. The slides were immersed in 300 mL of 0.1M glucose solution to which 50mL of GUVs were added. These were incubated for 3 hours with 1 mM clovibactin and later observed under confocal microscope Zeiss LSM 880. GUVs were imaged using Zeiss LSM 880 with 63x/1.2NA glycerol and 100x/1.2NA oil objective lenses. The Atto 550 label appeared red upon excitation by the 560 nm laser. The brightfield was used for detection and location of the GUVs and to observe their shape. ImageJ software(Schindelin et al., 2012) was used for the analysis of the images.

### Isothermal Titration Calorimetry

For ITC measurements LUVs (Large unilamellar vesicles) containing Lys-Lipid II were prepared by incorporating 2 mol% of Lys-Lipid II in DOPC from the stock solution. The lipids were dried under a nitrogen stream and hydrated with buffer (30mM Citrate, 300mM NaCl, pH 5.5) to a lipid-phosphate concentration of 20mM determined by Rouser’s method(Rouser et al., 1970). Finally, unilamellar vesicles were obtained after 10 rounds of extrusion through 200nm membrane filters (Whatman Nuclepore, Track-Etch Membranes). ITC experiments were performed with the Affinity ITC (TA Instruments-Waters LLC, New Castle, DE, USA) to determine interaction between LUVs and clovibactin. Clovibactin was diluted in the buffer, to a final concentration of 30 mM. The samples were degassed before use. The chamber was filled with 177 mL of clovibactin, and the LUVs were titrated into the chamber at a rate of 1.96 mL/150 seconds with a constant syringe stirring rate of 125 rpm. Number of injections = 21. Experiments were performed at 37°C and analyzed using the Nano Analyze Software (TA instruments – Water LLC). All experiments were performed in triplicates. Control experiments were performed with Lipid II-free DOPC LUVs. The independent model was used to determine the interaction between clovibactin and lipid II. ITC data for teixobactin was recently published(Shukla et al., 2022a).

### High Speed-Atomic Force Microscopy (HS-AFM) Imaging

The HS-AFM images were acquired in amplitude modulation tapping mode in liquid using a high-speed atomic force microscope (RIBM, Japan). Short cantilevers (~7 μm) with a nominal spring constant of 0.15 N/m were used (USC-F1.2-k0.15, NanoWorld, Switzerland). A minimal imaging force was applied by using a small set-point amplitude of 0.8 nm (for a 1 nm free amplitude). The HS-AFM results showing the assembly of clovibactin filaments were obtained from imaging of supported lipid bilayers on mica. The lipid bilayer was obtained by incubating LUVs containing 0.5 mg/ml of DOPC and 4 mol% Lipid II (prepared as mentioned above) mixture (or 0.5 mg/ml DOPC without lipid II) on top of a freshly cleaved mica for 20-30 minutes. After the incubation period, the mica was cleaned gently using recording buffer (10 mM Tris-Cl, 100 mM NaCl, pH 8.0). Imaging was started on the lipid bilayer surface in recording buffer. Next, a concentrated clovibactin solution was added and pipetted to reach the desired final concentration in the AFM liquid chamber of 40 µl. Images were primarily processed using built-in scripts (RIBM, Japan) in Igor Pro (Wavemetrics, Lake Oswego, OR, USA) and analyzed using ImageJ software. The images/movies were corrected minimally for tilt, drift, and contrast. Unless otherwise mentioned, the times reported in AFM images are relative to clovibactin addition into the imaging chamber. Reported image acquisition rate is 0.5 frames/second, and the line rate is 150 lines/second. Stated errors are standard deviation.

### Permeabilization assay

The bacterial cultures were grown overnight at 37°C in LB media for *B. subtilis*. Secondary cultures were grown for 3 hours until the OD_600_ = 0.5 was reached. The bacterial cells were then centrifuged at 1500x g for 10 minutes at 4°C and washed twice with 10 mL of buffer (10 mM Tris, 100 mM NaCl, 1 mM MgCl_2_, 0.5% glucose, pH=7.2). The bacterial cells were resuspended to an OD_600_ = 10 in the buffer and used for the experiment. All permeability experiments were performed with a Cary Eclipse (FL0904M005) fluorometer. All samples (1.0 mL) were continuously stirred in a 10 × 4-mm quartz cuvette and kept at 20°C. For the assay, 1 mL of the bacterial suspension was added to 1mL of buffer. For ion leakage assays, 1mL of the DISC-2 probe from a 1mM stock was added to the cuvette and the fluorescence was measured between 650nm to 670nm wavelength (bandwidth = 5mm) for 2 minutes before the addition of the antibiotic and 6 minutes after. For Sytox green leakage assays, 1 mL of the Sytox green probe from a 0.25mM stock was added to the cuvette and the fluorescence was measured between 500nm to 520nm wavelength (bandwidth = 5mm) for 2 minutes before the addition of the antibiotic and 6 minutes after. All experiments were performed in triplicates. The concentrations of antibiotics used are 10 nM Nisin (1 x MIC) and 2 mM clovibactin (1 x MIC) and 0.2 mM teixobactin (10 x MIC) for *B. subtilis*.

### Structure Calculations

#### Parametrization of Clovibactin

Clovibactin parametrization was started from a linear peptide topology of L-amino acids. D-amino acids were generated by inverting relevant dihedral and improper torsion angles and hydroxyasparagine (Hyn) topology was based on asparagine, where −OH group parameters were derived from threonine −OH group parameters. Depsi-cycle formation between Hyn5 and Leu8 was performed as previously described for [R4,L10]-teixobactin (Shukla et al., 2020). Parameters for lipid II were taken from ref. (Hsu et al., 2004).

#### Structure calculation protocol

We used HADDOCK version 2.4 (van Zundert et al., 2016) for the structure calculations. An eight-body docking (four lipid II and four clovibactin molecules) was performed using ssNMR-derived distance and dihedral restraints. Five thousand models were generated in the rigid-body docking stage of HADDOCK, of which the best-scoring 500 were subjected to the flexible refinement protocol of HADDOCK. The resulting models were energy minimized. Default HADDOCK settings were used except for doubling the weight of the distance restraints during all stages of the structure calculation. The final models were further filtered based on the topological requirements (that is, the lipid tails of all lipid II molecules must point in the same direction as the membrane-exposed hydrophobic residue Leu2). This resulted in a final ensemble of 22 structure models. See Supporting Information for detailed analysis of the calculated structure models.

