## Supplemental data for "A new antibiotic from an uncultured bacterium binds to an immutable target"

| Primer Name | Primer sequence 5'-3' |
|---|---|
| bat1kofrag2F | agactaGGATCCctgcgggtggaccccgt |
| bat1kofrag2R | acagtaTCTAGAgcgcacactccctactcggtc |
| bat1kofrag1F | agactaGGATCCgacgggccttacgcagg |
| bat1kofrag1R | agactGGTACCactccacagcagtgttgaaggt |
| Bat1downstream | gcgctggccctcggaaag |
| Bat2upstream | gcgccgcttggacacttat |

**Supplementary Table 1:** Primer DNA sequences used in the interruption of bat1 in P9846. Capitalized letters highlight the restriction enzyme cuts sites used in cloning fragments into pIJ12738.

| | **HN** | **Hα** | **Hβ** | **Others** |
|---|---|---|---|---|
| **L-Phe (1)** | 8.21 | 4.11 | 3.01 | $H\delta_{(1,2)}$: 7.27, $H\varepsilon_{(1,2)}$: 7.33, Hζ: 7.28 |
| **D-Leu (2)** | 8.62 | 4.30 | 1.34/1.52 | Hγ: 1.21, $H\delta_1$: 0.75, $H\delta_2$: 0.79 |
| **D-Lys (3)** | 8.22 | 4.33 | 1.53/1.68 | Hγ: 1.28/1.33, Hδ: 1.52, Hε: 2.74, Hζ: 7.78 |
| **L-Ser (4)** | 7.91 | 4.41 | 3.41/3.59 | Hγ: N.A. |
| **3-OH-Asn (5)** | 8.10 | 5.03 | 5.28 | Hδ: 7.40/7.88 |
| **L-Ala (6)** | 8.32 | 3.99 | 1.34 | - |
| **L-Leu (7)** | 7.62 | 4.37 | 1.51 | Hγ: N.A., $H\delta_1$: N.A., $H\delta_2$: N.A. |
| **L-Leu (8)** | 7.76 | 4.35 | 1.51/1.66 | Hγ: N.A., $H\delta_1$: N.A., $H\delta_2$: N.A. |
| | **N** | **Cα** | **Cβ** | **Others** |
| **L-Phe (1)** | 39.20 | 53.63 | 37.52 | Cγ:135.32, $C\delta_{(1,2)}$: 129.94, $C\varepsilon_{(1,2)}$: 128.98, Cζ: 127.63 |
| **D-Leu (2)** | 123.10 | 51.26 | 41.43 | Cγ: 23.98, $C\delta_1$: 21.78, $C\delta_2$: 23.51 |
| **D-Lys (3)** | 118.00 | 52.54 | 31.45 | Cγ: 22.57, Cδ: 26.77, Cε: 38.97, Nζ: 33.30 |
| **L-Ser (4)** | 113.30 | 56.14 | 61.71 | - |
| **3-OH-Asn (5)** | 109.60 | 53.46 | 73.74 | Cγ: N.A., Nδ: 106.1 |
| **L-Ala (6)** | 123.00 | 52.31 | 17.06 | - |
| **L-Leu (7)** | 110.00 | 52.72 | 41.89 | Cγ: N.A., $C\delta_1$: N.A., $C\delta_2$: N.A. |
| **L-Leu (8)** | 113.60 | 50.96 | 40.32 | Cγ: N.A., $C\delta_1$: N.A., $C\delta_2$: N.A. |

**N.A.: Not Assigned**

**Supplementary Table 2**: $^{1}$H and $^{13}$C solution NMR data of clovibactin in DMSO-*d*6 (500 MHz, δ in ppm).

| Organism (Gram-positives) | # Isolates | Clovibactin MIC range (µg/mL) | Vancomycin MIC range (µg/mL) |
|---|---|---|---|
| *Staphylococcus aureus* (MSSA) | 3 | 1 | 0.5 - 1 |
| *Staphylococcus aureus* (MRSA) | 3 | 1 | 0.5 - 1 |
| *Staphylococcus aureus* (VISA) | 3 | 1 - 4 | 4 |
| *Staphylococcus aureus* (DAP$^{NS}$) | 3 | 1 - 4 | 4 - 8 |
| *Staphylococcus aureus* (LZD$^{R}$) | 3 | 0.5 - 2 | 0.5 - 1 |
| *Staphylococcus epidermidis* (MSSE) | 1 | 1 | 1 |
| *Staphylococcus epidermidis* (MRSE) | 2 | 0.5 - 1 | 1 |
| *Streptococcus pneumoniae* (PSSP) | 3 | 0.25 – 0.5 | 0.25 |
| *Streptococcus pneumoniae* (PISP) | 4 | 0.5 - 1 | 0.12 - 0.25 |
| *Streptococcus pneumoniae* (PRSP) | 3 | 0.25 – 0.5 | 0.25 |
| *Streptococcus pyogenes* | 4 | 0.5 - 1 | 0.25 |
| *Streptococcus agalactiae* | 3 | 1 | 0.25 |
| *Streptococcus anginosus* | 1 | 0.5 | 0.5 |
| *Streptococcus mutans* | 1 | 1 | 0.5 |
| *Streptococcus salivarius* | 1 | 0.25 | 0.25 |
| *Enterococcus faecium* (VSE) | 3 | 2 | 0.5 - 1 |
| *Enterococcus faecium* (VanA VRE) | 2 | 1 - 2 | ≥ 32 |
| *Enterococcus faecium* (VanB VRE) | 2 | 1 - 2 | ≥ 32 |
| *Enterococcus faecalis* (VSE) | 3 | 4 - 8 | 0.5 - 4 |
| *Enterococcus faecalis* (VanA VRE) | 2 | 2 - 4 | > 32 |
| *Enterococcus faecalis* (VanB VRE) | 2 | 4 | > 32 |
| **Organism (Gram-positives)** | **# Isolates** | **Clovibactin MIC range (µg/mL)** | **Ciprofloxacin MIC range (µg/mL)** |
| *Escherichia coli* ATCC 25922 | 1 | > 64 | 0.008 |
| *Escherichia coli* (ESBL) | 1 | > 64 | > 32 |
| *Klebsiella pneumoniae* | 1 | 64 | 2 |
| *Klebsiella pneumoniae* (KPC) | 1 | > 64 | > 32 |
| *Proteus vulgaris* | 1 | > 64 | 0.015 |
| *Proteus mirabilis* | 1 | > 64 | 8 |
| *Enterobacter cloacae* | 2 | ≥ 64 | 0.008 – 0.015 |
| *Enterobacter aerogenes* | 2 | > 64 | 0.03 – 0.06 |
| *Serratia marcescens* | 2 | > 64 | 0.06 |
| *Pseudomonas aeruginosa* | 2 | > 64 | 32 |
| *Acinetobacter baumannii* | 2 | > 64 | > 32 |
| *Haemophilus influenzae* | 3 | >64 | 0.06 – 0.25 |
| *Moraxella catarrhalis* | 2 | 4 - 16 | 0.03 |
| *Neisseria gonorrhoeae* | 2 | ≥ 64 | 0.002 - 16 |

**Supplementary Table 1: Antibacterial activity of clovibactin and known drugs against contemporary clinical isolates.** Susceptibility testing determined by broth microdilution in accordance with CLSI guidelines was conducted by Micromyx LLC, Kalamazoo, MI, USA. MSSA, methicillin-sensitive *S. aureus;* MRSA, methicillin-resistant *S. aureus;* VISA, vancomycin-intermediate *S. aureus;* DAP$^{NS}$, daptomycin non-susceptible; LZD$^{R}$, linezolidresistant; MRSE, methicillin-resistant *S. epidermidis;* MRSH, methicillin-resistant *S. haemolyticus;* PSSP, penicillin-susceptible *S. pneumoniae;* PISP, penicillin-intermediate *S. pneumoniae;* PRSP, penicillin-resistant *S. pneumoniae;* VSE, vancomycin-sensitive enterococci; VRE, vancomycin-resistant enterococci, ESBL, extended spectrum β-lactamase positive; KPC, *Klebsiella pneumoniae* carbapenemase-positive

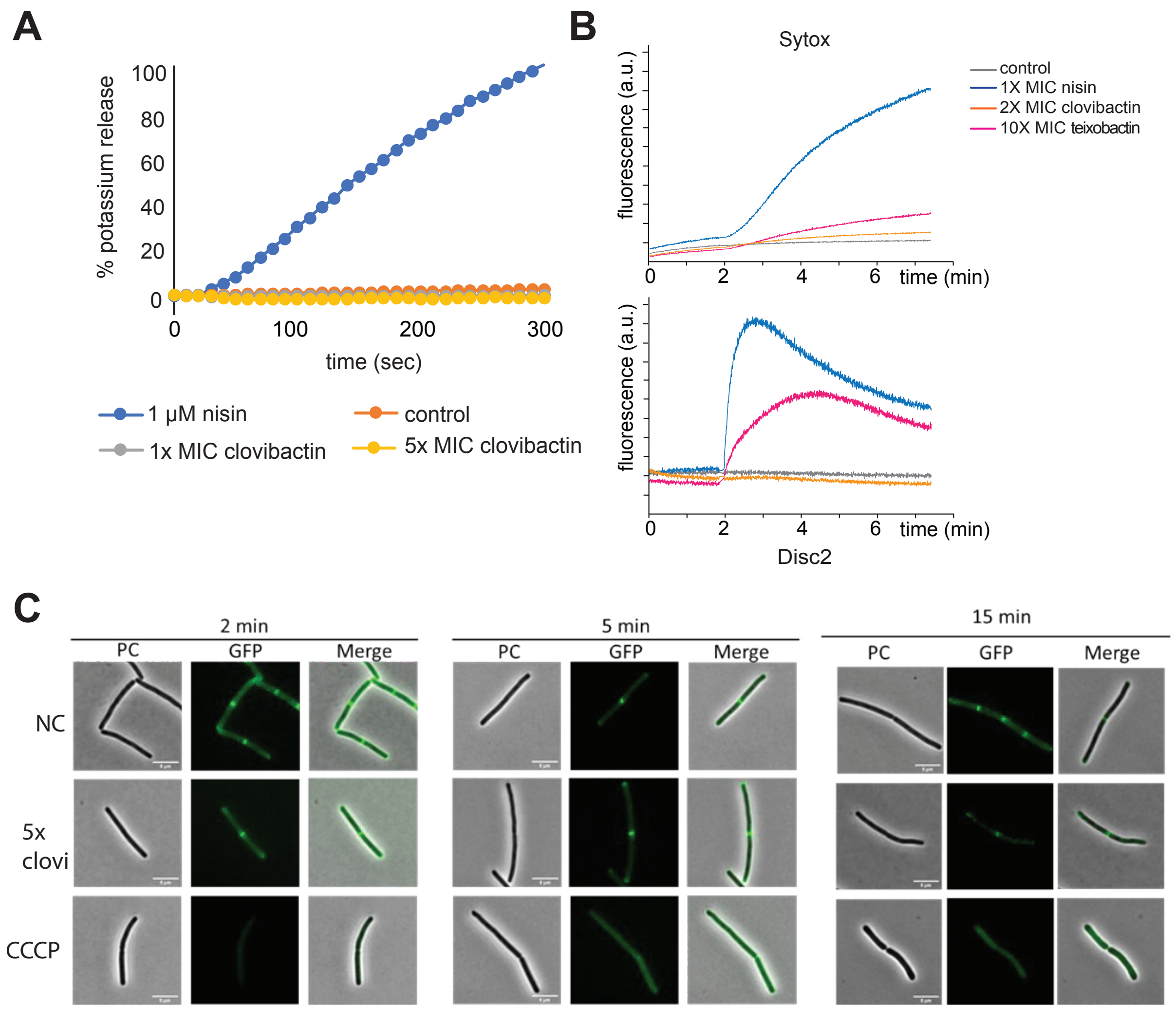

**Supplementary Figure 1: Clovibactin does not disrupt the membrane.** (A) Potassium efflux from living cells of *S. simulans* was monitored with a potassium-sensitive electrode. Ion leakage is expressed relative to the total amount of potassium released after addition of 1 µM pore-forming lantibiotic nisin (100%). (B) Permeabilization assay with DiSC(2) and Sytox reporter dye for *B. subtilis* with nisin, teixobactin, clovibactin and untreated cells. While nisin and teixobactin compromise the membrane integrity, clovibactin shows no permeabilizing activity for either of the two reporters. (C) GFP-MinD cells of *B. subtilis* were treated with clovibactin (5xMIC). The ionophore CCCP was used as a positive control. Scale bar = 5 µm.

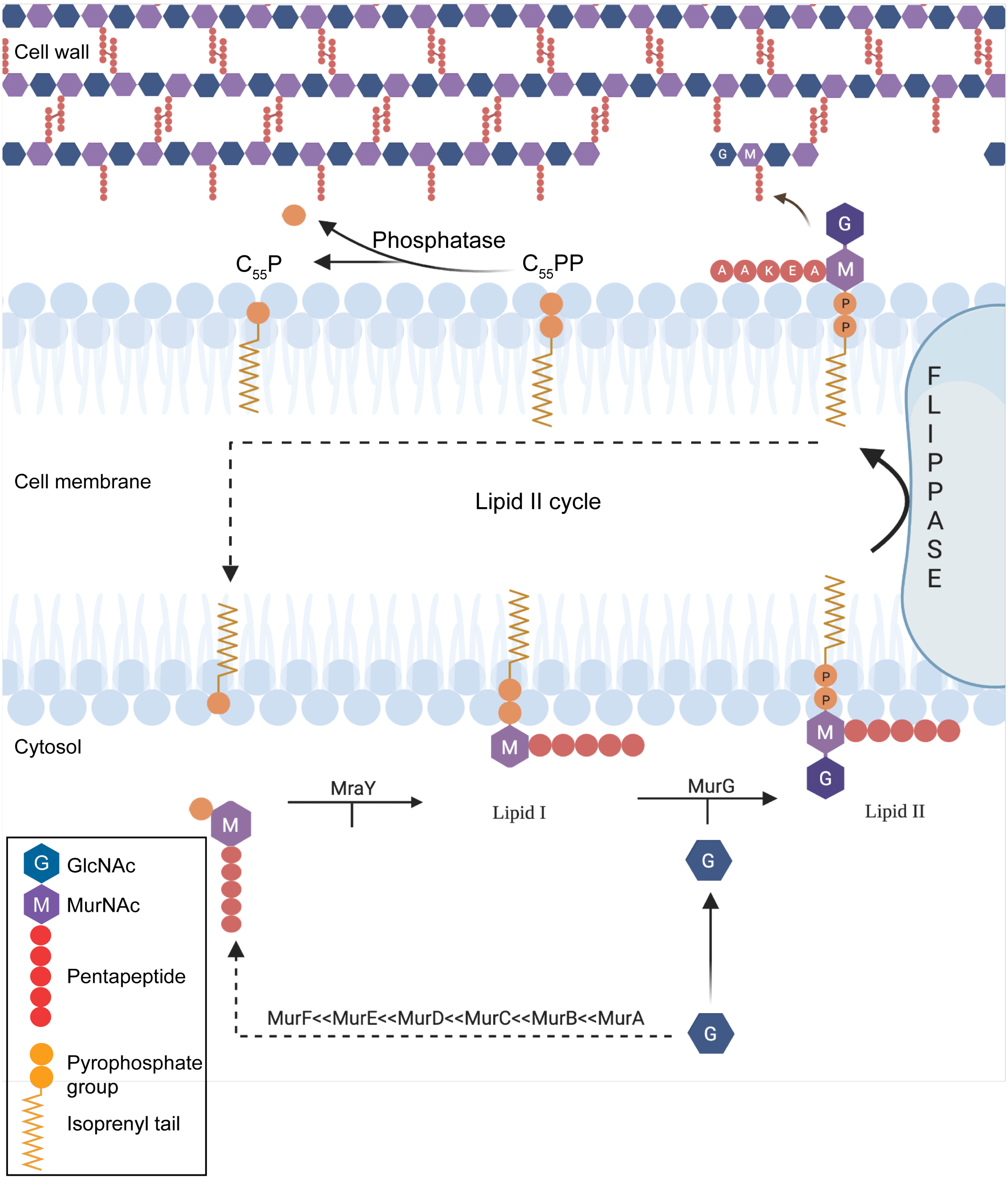

**Supplementary Figure 2: Schematic of peptidoglycan synthesis.** Peptidoglycan synthesis is dependent on the lipid II cycle which is shown in the illustration. It shows the organization of the lipid II molecule on the inner leaflet which is eventually flipped to the outer leaflet. When present in the outer leaflet lipid II deposits the disaccharide moiety (GlcNAc-MurNAc) and the attached pentapeptide to develop the peptidoglycan layer. The $C_{55}P$ moiety is then recycled to continue the cycle.

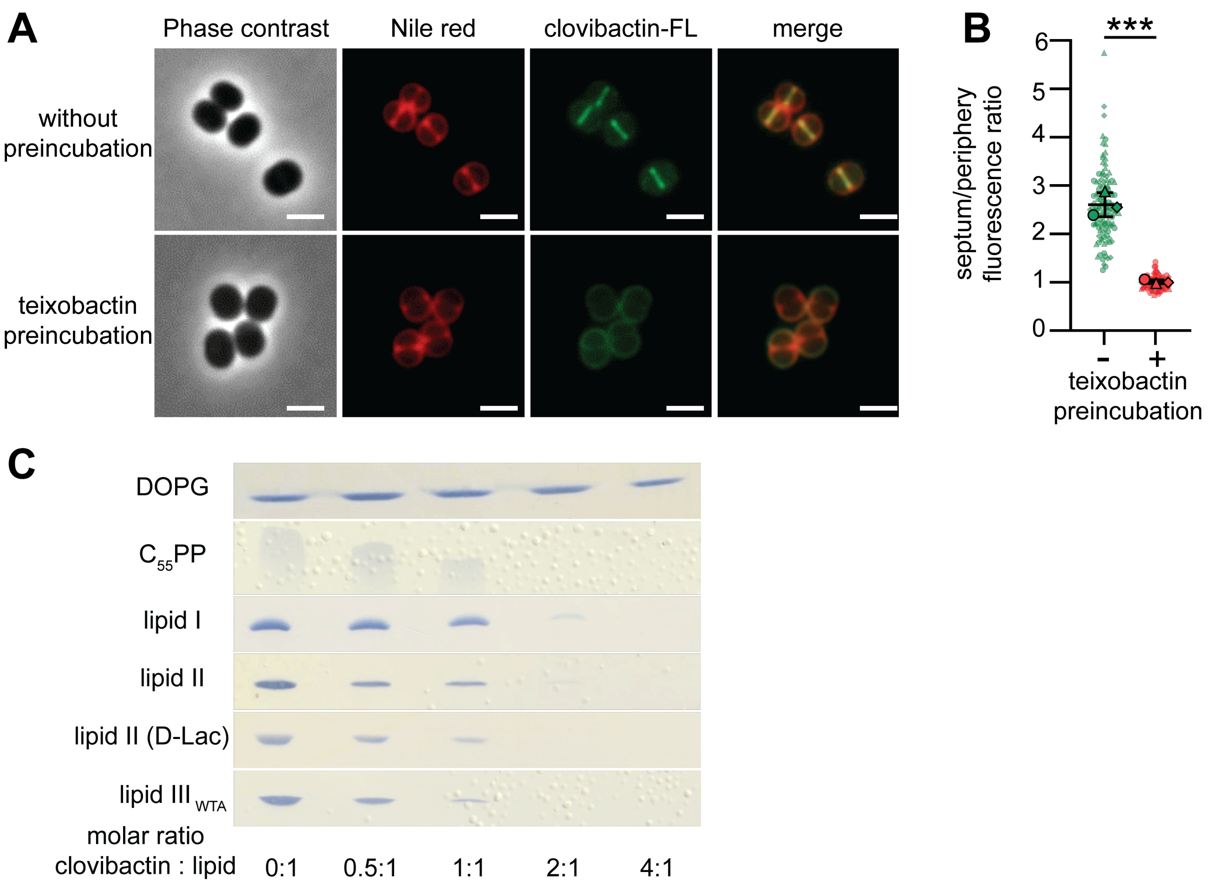

**Supplementary Figure 3: BODIPY-FL-clovibactin localizes to the septum of *S. aureus* and forms extraction-stable complexes with undecaprenyl pyrophosphate-containing cell wall precursors.** (A) In cells pre-incubated with teixobactin (2 min), septal localization of FL-clovibactin is diminished. Nile red was used for membrane staining. Scale bar 1 µm. (B) Images of individual cells were used to calculate the fluorescence ratio (FR) of the septal versus peripheral fluorescence signal for cells pre-treated with teixobactin and lacking pre-treatment. FR >2 indicates septal localization. At least $n = 30$ cells were evaluated for each condition from three biologically independent experiments. Significance was determined by unpaired Student's *t*-test with a 95% confidence interval. ****$p < 0.0004$.
(C) Complex formation is indicated by a reduction of the amount of free lipid intermediates visible on the TLC. Cell wall intermediates are fully locked in a complex at a twofold molar excess of antibiotic. No complex formation was observed with the anionic phospholipid DOPG. Note that the molar clovibactin : lipid ratios are not representative for stoichiometries. The chromatograms are representative of three independent experiments.

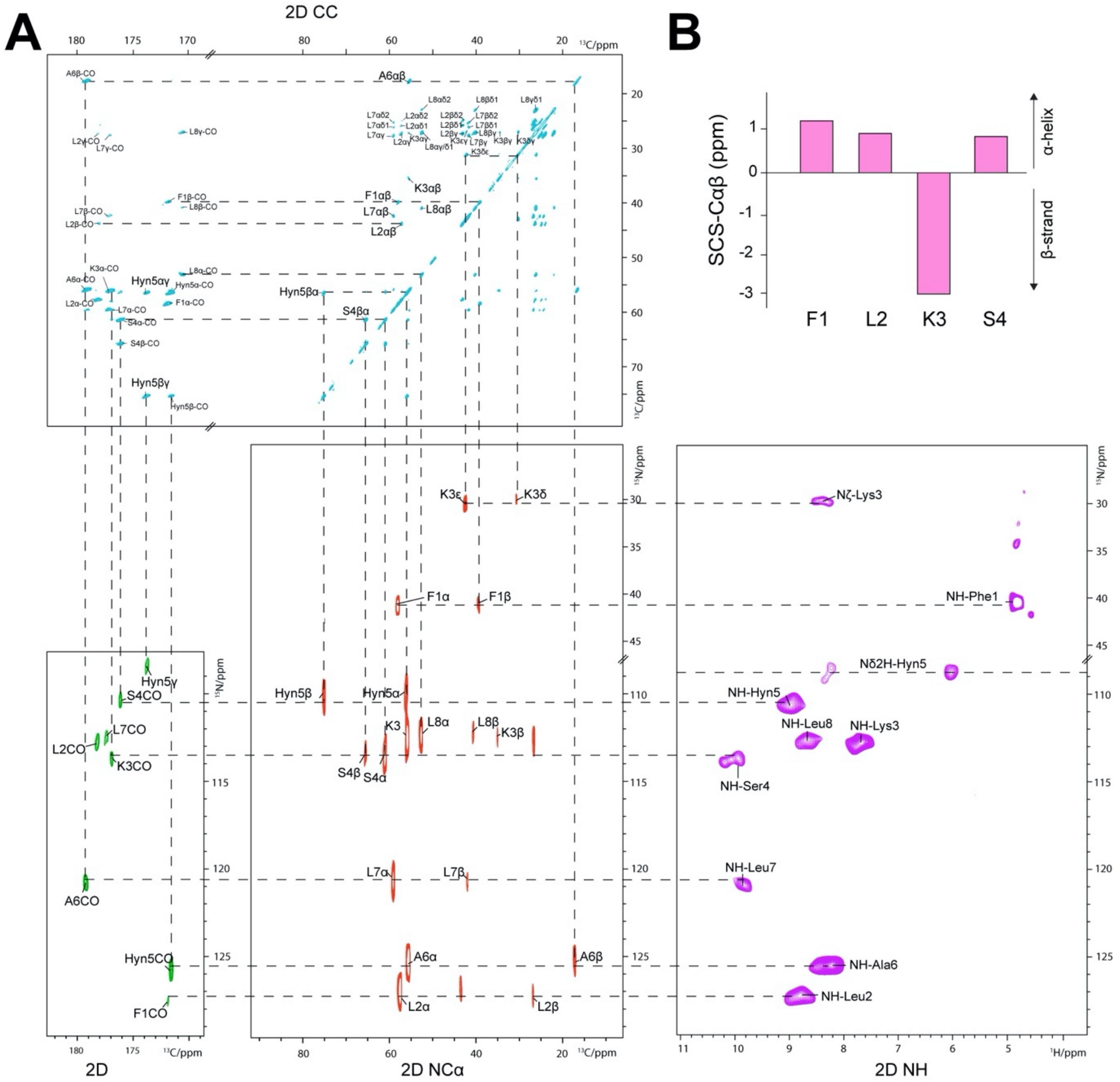

**Supplementary Figure 4: ssNMR assignments of clovibactin in the lipid II-bound state in DOPC liposomes.** Assignments were performed using a 2D PARIS CC experiments (in blue), 2D NCO(i-1) (in green), 2D NCA (in red), and 2D NH (in purple). 2D PARIS CC ssNMR experiment was recorded at 1200 MHz magnetic field at 18k Hz MAS. 2D NCO(i-1) and 2D NCA experiments were recorded at 800 MHz magnetic field at 15 kHz MAS. 2D NH experiment was recorded at 700 MHz magnetic field at 16.5 kHz MAS.

B) Cαβ Secondary chemical shifts (SCS) do not show consistent β-structuring for the linear clovibactin N-terminus. We note that it is unclear to which standard chemical shift information (Wang, 2002) extent chemical shift information applies to the very short N-terminus with its unusual structure and non-canonical residues. We note that chemical shift of teixobactin (Shukla et al., 2022), residues D-N-Me-Phe1 and Ser7, i.e., the residue connected to the depsi-cycle, also did not indicate β-structuring.

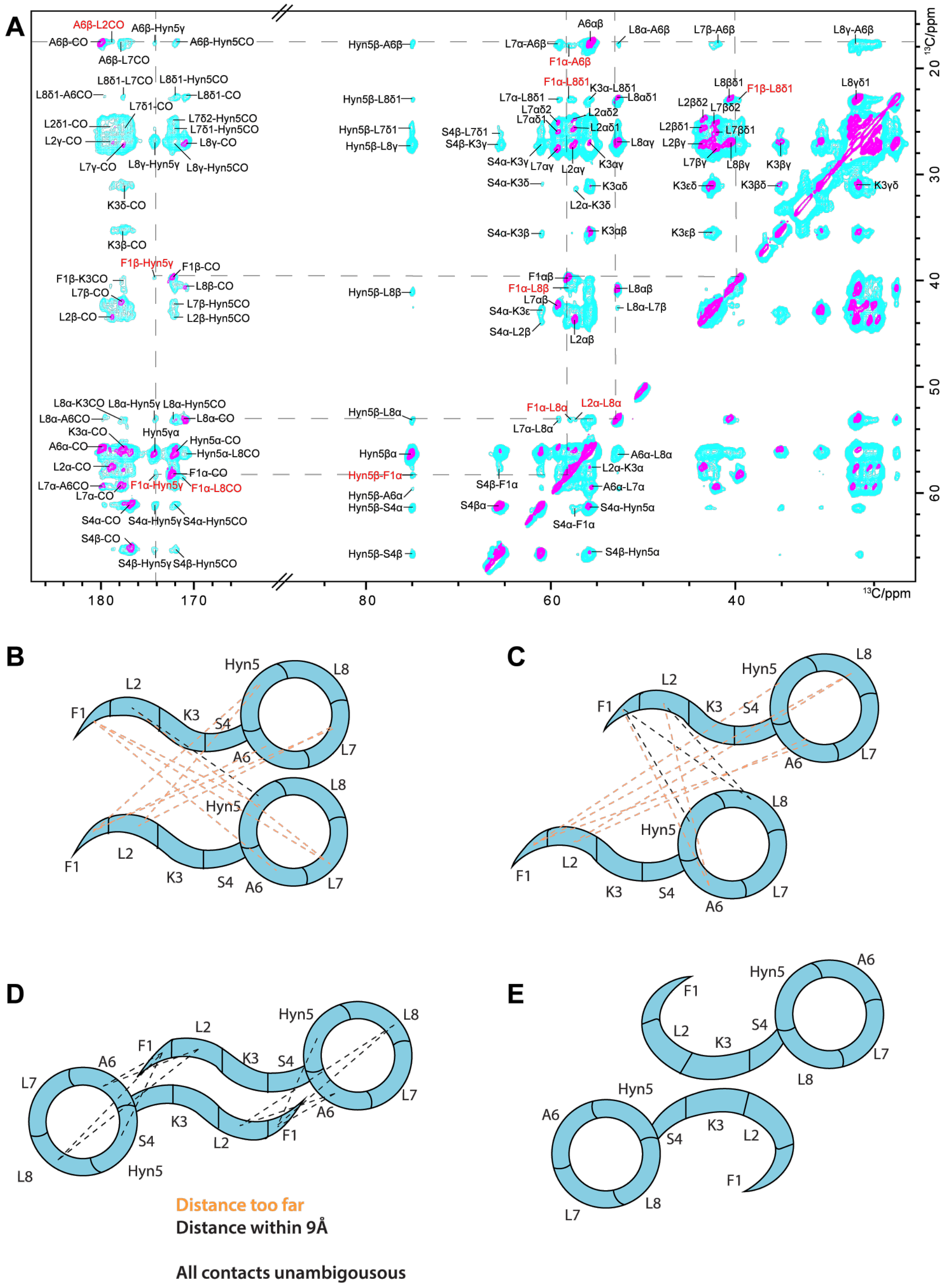

**Supplementary Figure 5: Antiparallel arrangement of the clovibactin dimer.** ssNMR, confocal microscopy, and AFM-data show that clovibactin assembled into fibrils upon lipid II binding. From ssNMR data, the nature of the fibrils can be gauged. (A) Superposition of 2D PARIS CC spectra of clovibactin in the lipid II-bound state with 50 ms (magenta) and 300 ms (cyan) mixing times measured at 1200 MHz magnetic field and 18 kHz MAS. The contacts (in red) very likely arise from intermolecular interactions between adjacent clovibactin molecules as they are beyond the 8 Å range of such an experiment. The illustrations show how these contacts are consistent with be an antiparallel arrangement (D) of the neighbouring clovibactin molecules. The parallel (B) and out of register parallel (C) arrangements still contain many contacts that would not be satisfied by the distance limit of the experiment. In the antiparallel arrangement, the N- and C-terminal residues of adjacent clovibactin molecules come close enough to justify observing contacts such as F1$\alpha$-L8$\alpha$ and F1$\alpha$-A6$\beta$. E) Due to the small size of clovibactin, we note that the ssNMR restraints could potentially also be fulfilled by alternative supramolecular arrangements in which the N-terminus bends towards the C-terminal depsi-cycle. However, such an alternative arrangement appears difficult to align with the formation of fibrils observed by AFM.

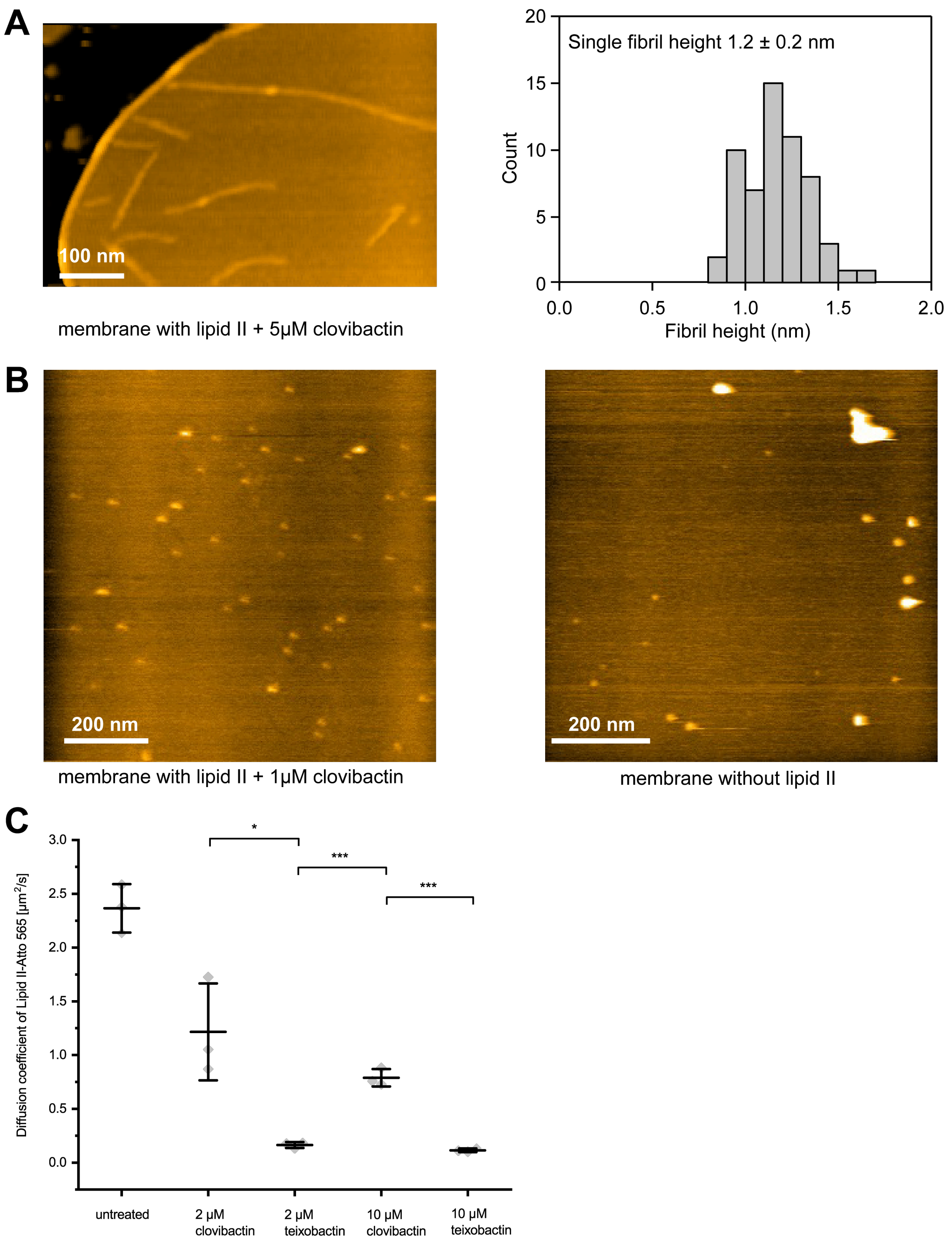

**Supplementary Figure 6: HS-AFM demonstrating the fibrilization of clovibactin and lipid II.** A) The HS-AFM experiments show the formation of the clovibactin fibrils only at high-concentrations of 5 μM and in the presence of lipid II. Also, the fibrils are loosely attached to the membrane and the average height of the fibrils is 1.2 ± 0.2 nm above the membrane. There was no visible membrane deformation, and this is in line with our permeabilization assays which also show no effect on bacterial membrane depolarization.
B) Control experiments with low clovibactin concentration (1 μM) at a membrane containing 4% lipid II (left image) and 5 μM clovibactin at a membrane containing no lipid II (right image) both show no fibrilization events.
C) Lipid II mobility was also monitored using single-molecule tracking microscopy experiments in supported lipid bilayers. These experiments also align with the requirement of higher clovibactin concentrations to immobilize lipid II in lipid bilayers. This contrasts with the low concentrations of teixobactin required for maximum lipid II immobilizations.

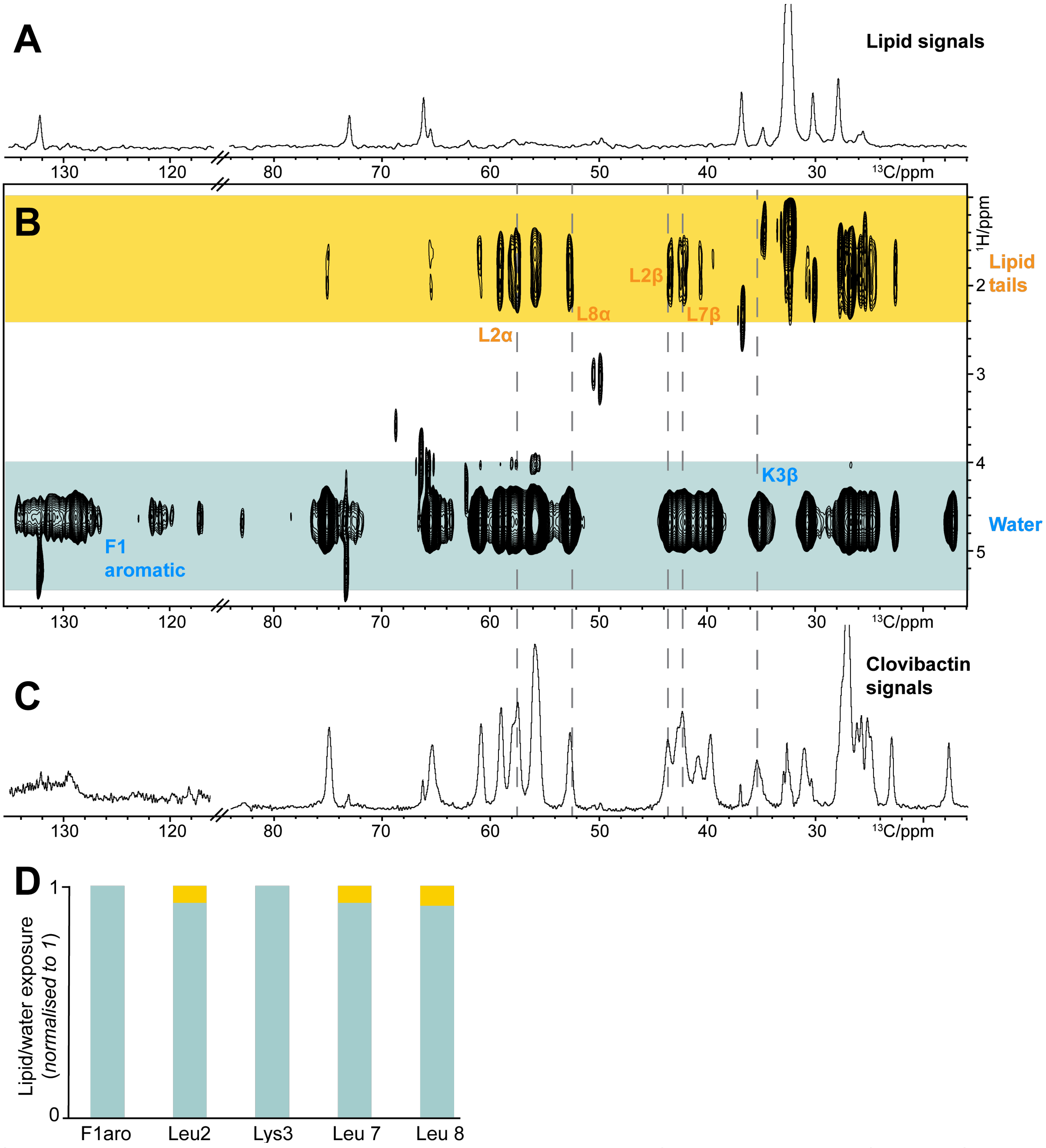

**Supplementary Figure 7: The membrane topology of the complex.** A) and B) show mobility-edited ('$T_2$-filtered') $^{1}$H($^{1}$H)$^{13}$C ssNMR experiments(Doherty and Hong, 2009) of $^{13}$C,$^{15}$N-clovibactin bound to Lipid II. Here, magnetization from mobile water and lipid molecules is transferred to the rigid clovibactin protons via $^{1}$H-$^{1}$H mixing and eventually transferred to the $^{13}$C nuclei of clovibactin via a short cross polarization step (200 µs). A 2D implementation of this experiment demonstrates that the sidechain of Leu2 is partially inserted in the membrane, whereas the sidechain of Lys3 points to the water phase. In general, as shown in Figure 4I of the main text, clovibactin fibrils do reside on the membrane surface and do not insert deeply into the membrane. Spectra were measured at 1200 MHz, 16.5 kHz MAS, and 300 K sample temperature. A) Control: 1D $^{1}$H($^{1}$H)$^{13}$C ssNMR spectrum (5120 scans) using a $T_2$-filter of 2.5 ms without transfer to clovibactin (0 ms $^{1}$H-$^{1}$H mixing). All signals relate to lipids, demonstrating the effectiveness of the $T_2$-filter. B) 2D $^{1}$H($^{1}$H)$^{13}$C ssNMR spectrum using 2.5 ms $T_2$ filter (1792 scans) and 5 ms $^{1}$H-$^{1}$H mixing. C) $^{13}$C cross-polarization spectrum (200 µs contact time) of Lipid II-bound teixobactin. D) Normalized relative signal intensities (peak heights) of the correlations with water (blue) and lipid (brown) protons for several residues. See Supplementary Figure 12 for a structural representation of the topology.

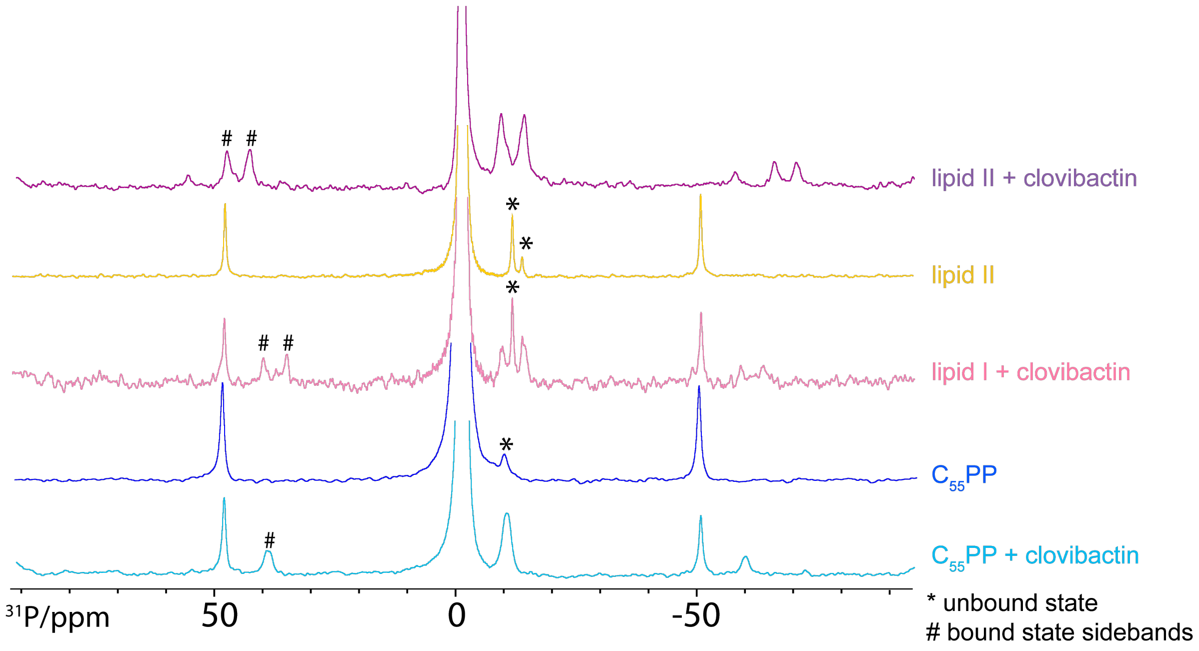

**Supplementary Figure 8: $^{31}$P magic angle spinning ssNMR of clovibactin with cell wall precursors in DOPC liposomes.** Clovibactin binds to all cell wall precursors containing the $C_{55}$PP unit ($C_{55}$PP, lipid I, lipid II). Changes in the chemical shifts of the pyrophosphate groups of lipid II and lipid I upon addition of clovibactin show the direct binding to the site. Furthermore, the emergence of strong spinning sidebands around +45/-55 ppm show that clovibactin binds and immobilises $C_{55}$PP. Experiments were performed using MLV's containing 2 mol% of the cell wall precursor with and without the treatment with clovibactin at 500MHz and 10 kHz MAS (lipid II + clovibactin was performed at 12 kHz MAS).

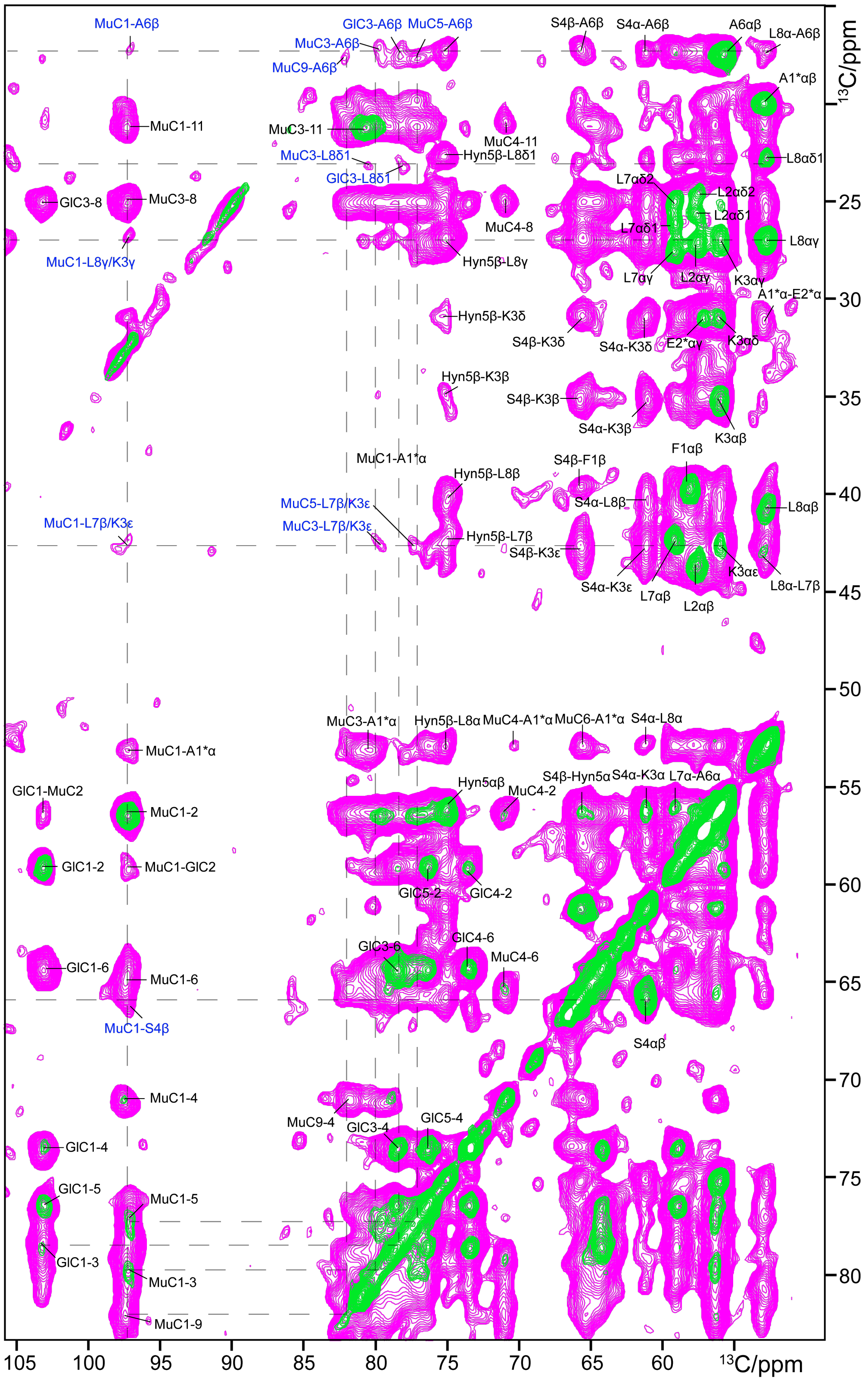

**Supplementary Figure 9: The binding interface and absence of the pentapeptide in the binding.** Spectrum showing multiple interfacial contacts illustrating the interaction between the depsi-cycle of clovibactin with the first sugar of lipid II. The experiments were performed at 950 MHz magnetic field with15.5 kHz MAS with a mixing time of 50 ms (green) and 300 ms (magenta).

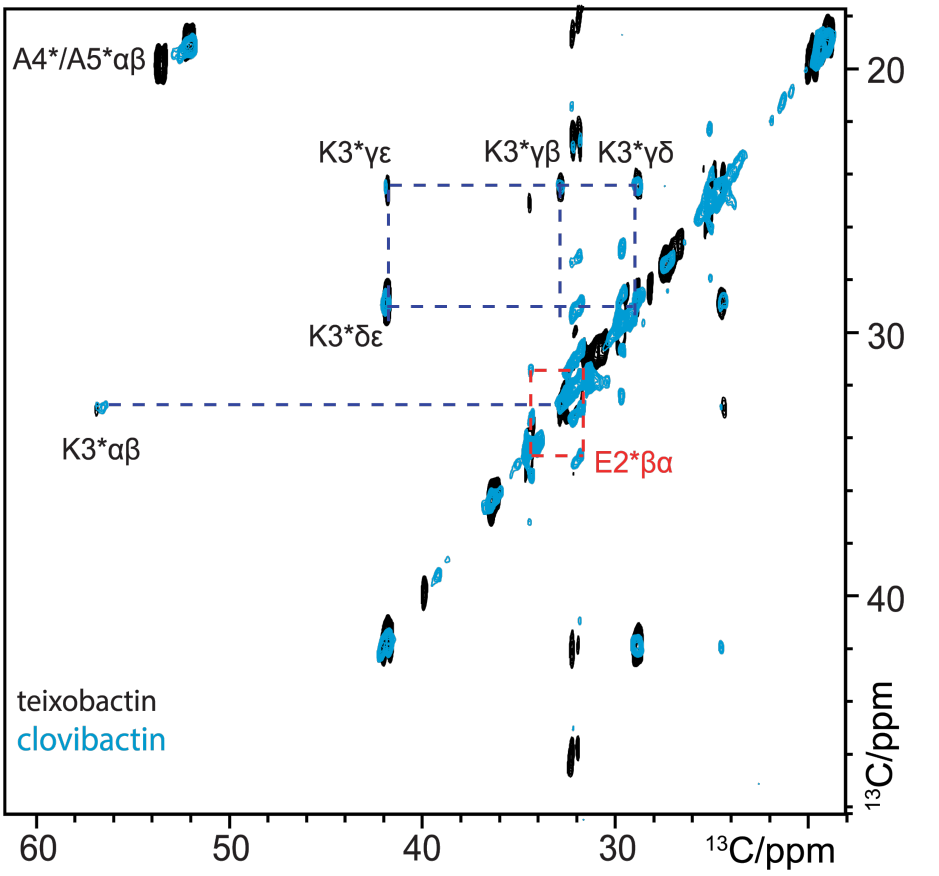

**Supplementary Figure 10: The TOBSY spectrum shows that the lipid II pentapeptide is not involved in complex formation.** The scalar based TOBSY spectrum only shows highly mobile residues in the complex. Here one can clearly see the presence of all the residues of the pentapeptide of lipid II, except for A1*. This indicates that the pentapeptide is highly mobile in the complex and not involved in the binding interface. The first A1* residue is linked to the first sugar (MurNAc) and therefore more rigid and in the dipolar based spectrum (Supp. Fig. 9). This contrasts with teixobactin's scalar spectrum (black) where residue E2 (in red) was not observed due to specific interactions of the MurNAc sugar with the enduracididine residue, leading to higher degree of rigidification of the pentapeptide.

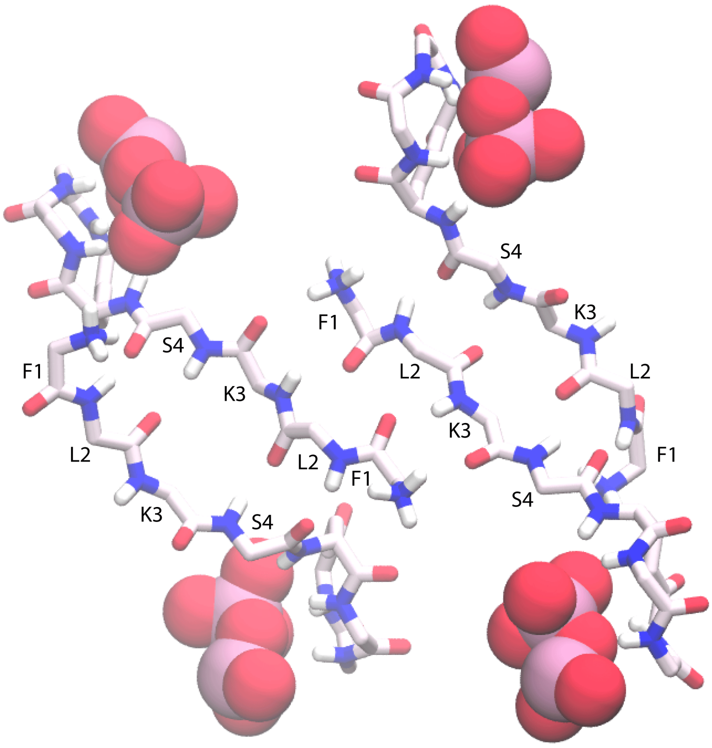

**Supplementary Figure 11: Hydrogen bonding in the structural model.** Atomic force microscopy, confocal microscopy, and ssNMR data demonstrate that clovibactin and Lipid II oligomerize into fibrils. ssNMR distance measurements are consistent with an antiparallel arrangement of clovibactin molecules, something that is also in line with the fibrils observed in AFM experiments.

Based on these experimental data, we applied hydrogen bonding restraints to foster an antiparallel arrangement in the structural model.

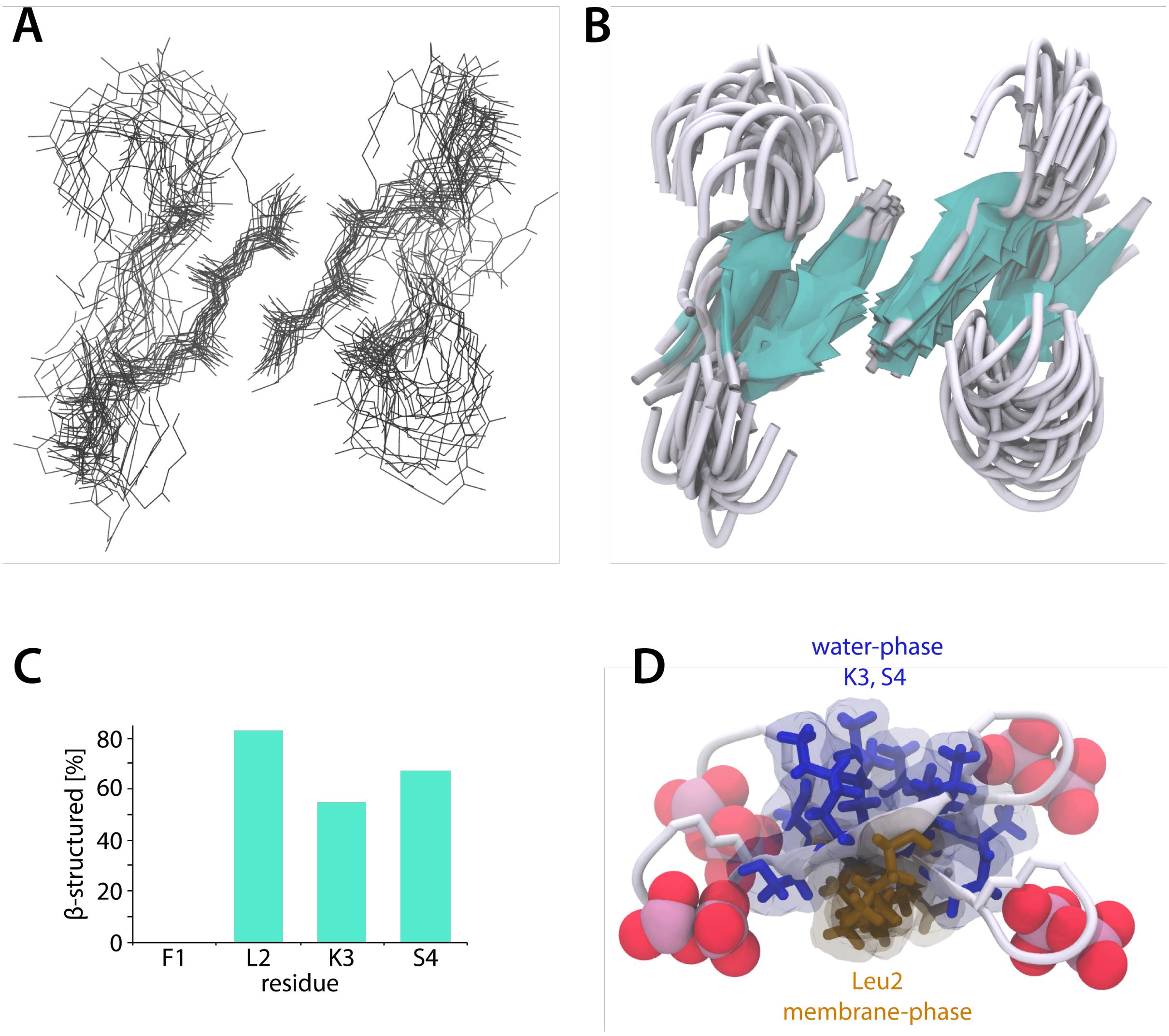

**Supplementary Figure 12: RMSD, secondary structure, and membrane topology.** (A, B) Superposition (2.50±0.85 Å average backbone RMSD for clovibactin in the complex) of 22 calculated structure models of the clovibactin – lipid II complex. Lipid II is not shown for clarity. (C) Quantification (Heinig and Frishman, 2004) of secondary structure elements of the very short linear N-terminus based on the calculated structural model. The N-terminus adopts a secondary structure that contains elements of β-structuring and an elongated loop. This is also reflected in the calculated ensemble (A, B) that does not show consistent β-structuring for the N-terminus. (D) Membrane-topology of clovibactin in the complex derived from ssNMR T2-edited H(H)C experiments (Doherty and Hong, 2009). The long-hydrophobic Leu2 sidechain (in brown) is embedded in the membrane, while the cationic (Lys3) and polar (Ser4) sidechains (in blue) of the N-terminus are water-exposed and, at the same time, favourably interact with the anionic PPi-group.

clovibactin - lipid II

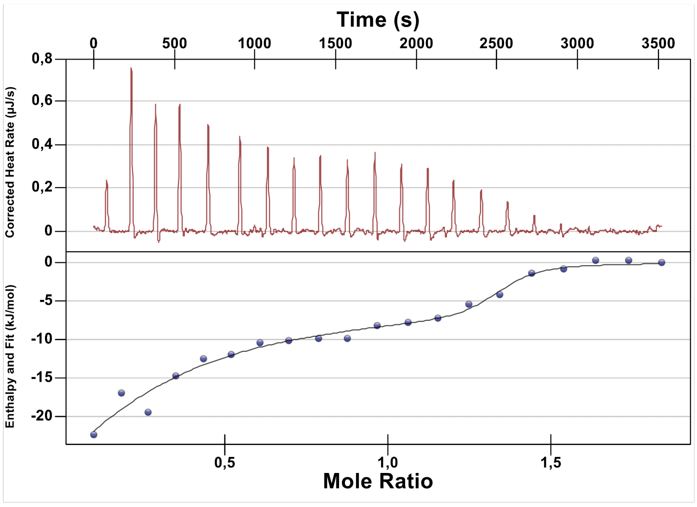

clovibactin - lipid I

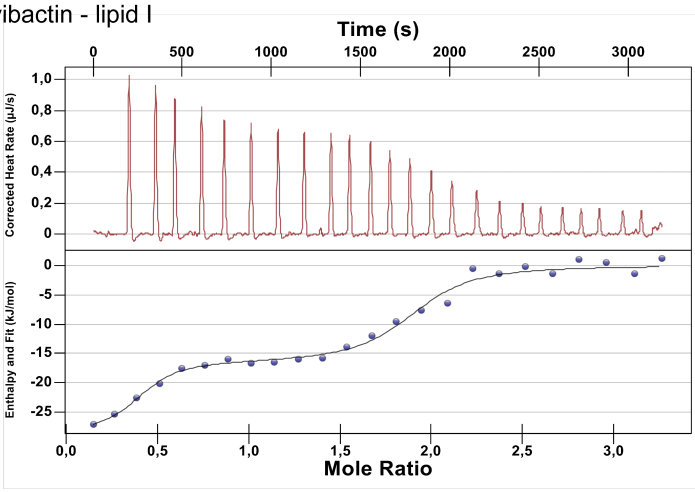

clovibactin - Na-pyrophosphate

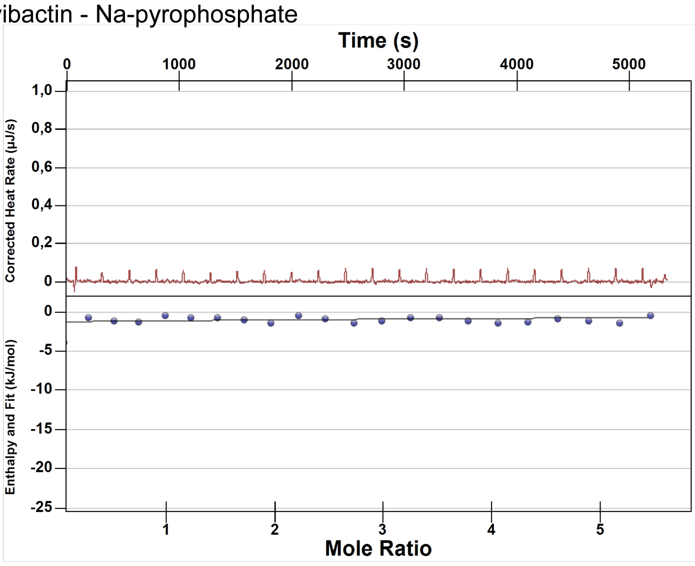

| Antibiotic | lipid II $K_d$ (µM) | lipid I $K_d$ (µM) |
|---|---|---|
| Clovibactin | 0.0862+/- 0.007 | 0.0807+/- 0.017 |

| | | |
|---|---|---|
| Teixobactin | 0.0146 +/-0.002 | 0.0805+/- 0.007 |

**Supplementary Figure 13: ITC data for clovibactin and teixobactin.** The ITC data shows a binding isotherm for interaction between clovibactin and lipid II and lipid I. When compared to teixobactin, clovibactin has lower affinity for lipid II. This is in line with the less specific interaction of the clovibactin depsi-cycle with the first sugar of lipid II. There is no binding observed between clovibactin and soluble Na-pyrophosphate.

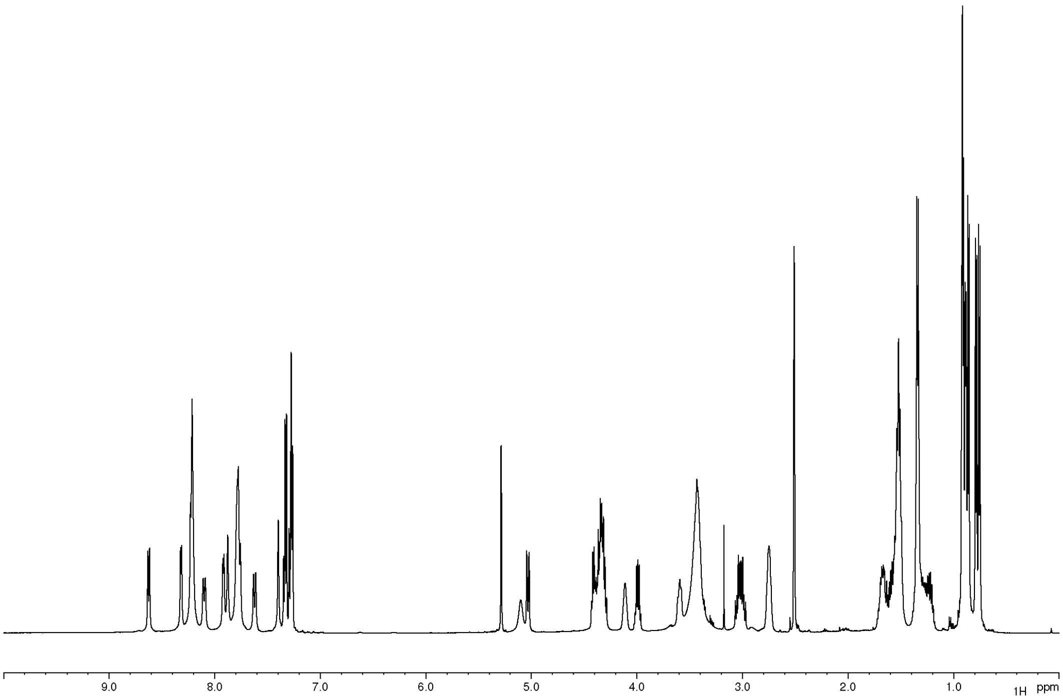

Supplementary Figure 15 $^{1}$H NMR Spectrum of Clovibactin

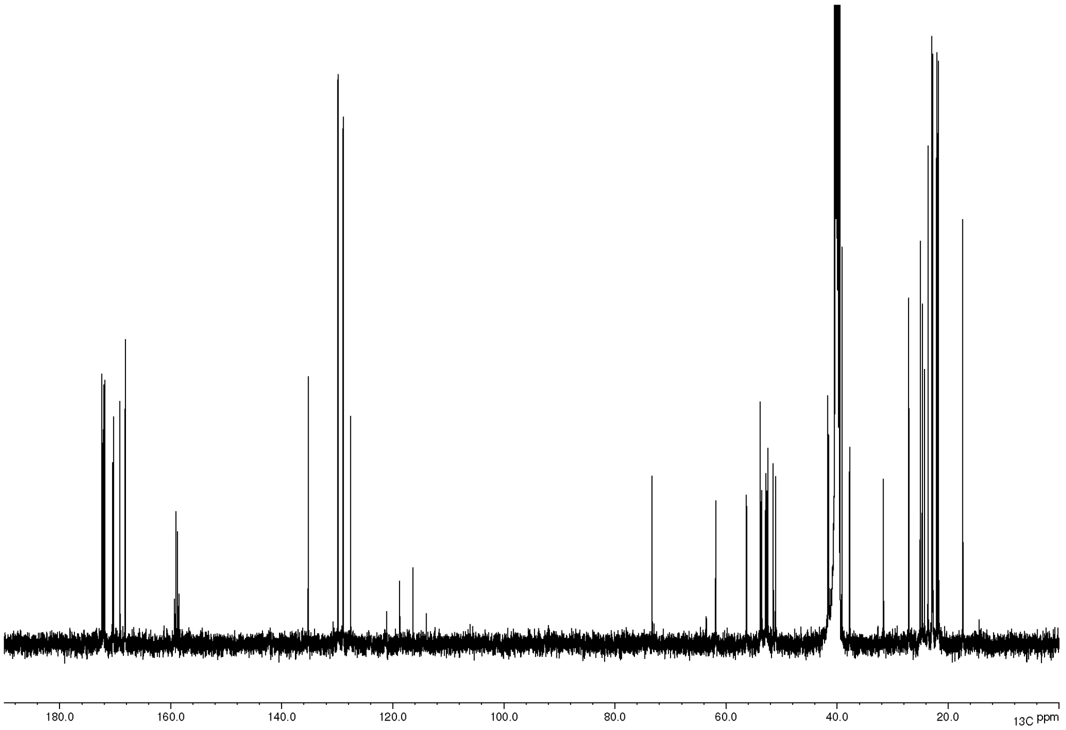

**Supplementary Figure 16.** $^{13}C$ solution NMR spectrum of clovibactin.

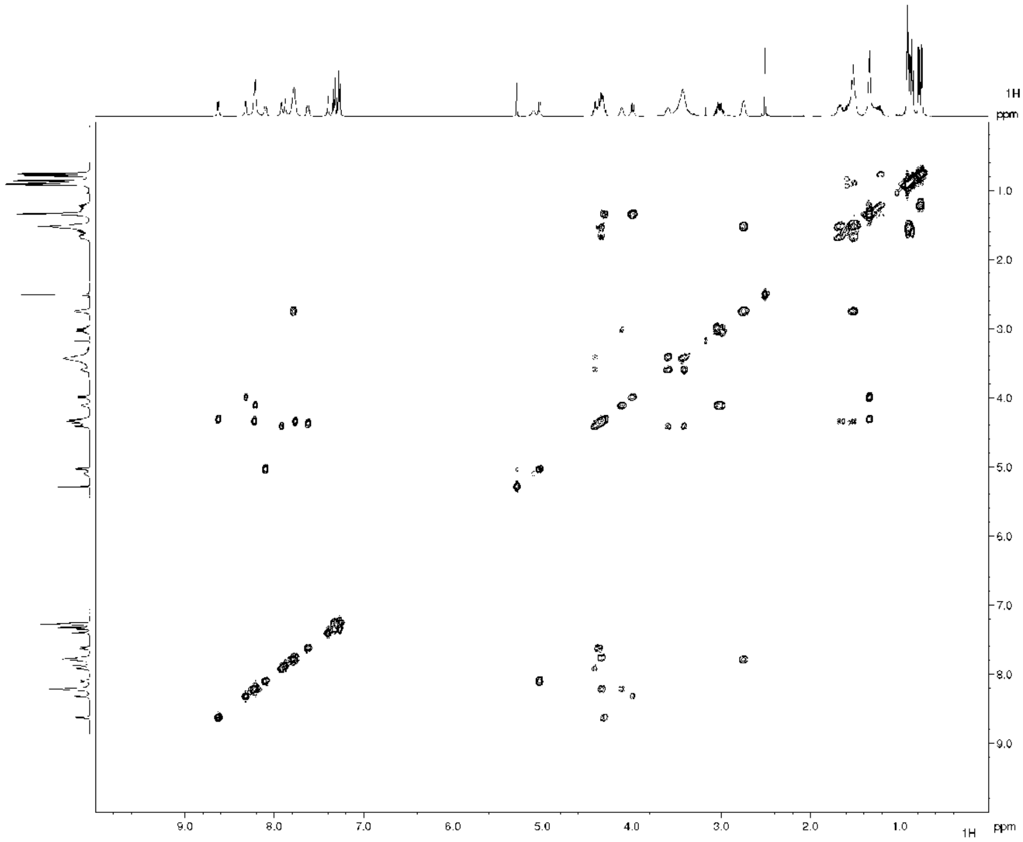

**Supplementary Figure 17.** $^{1}H$-$^{1}H$ COSY solution NMR Spectrum of clovibactin.

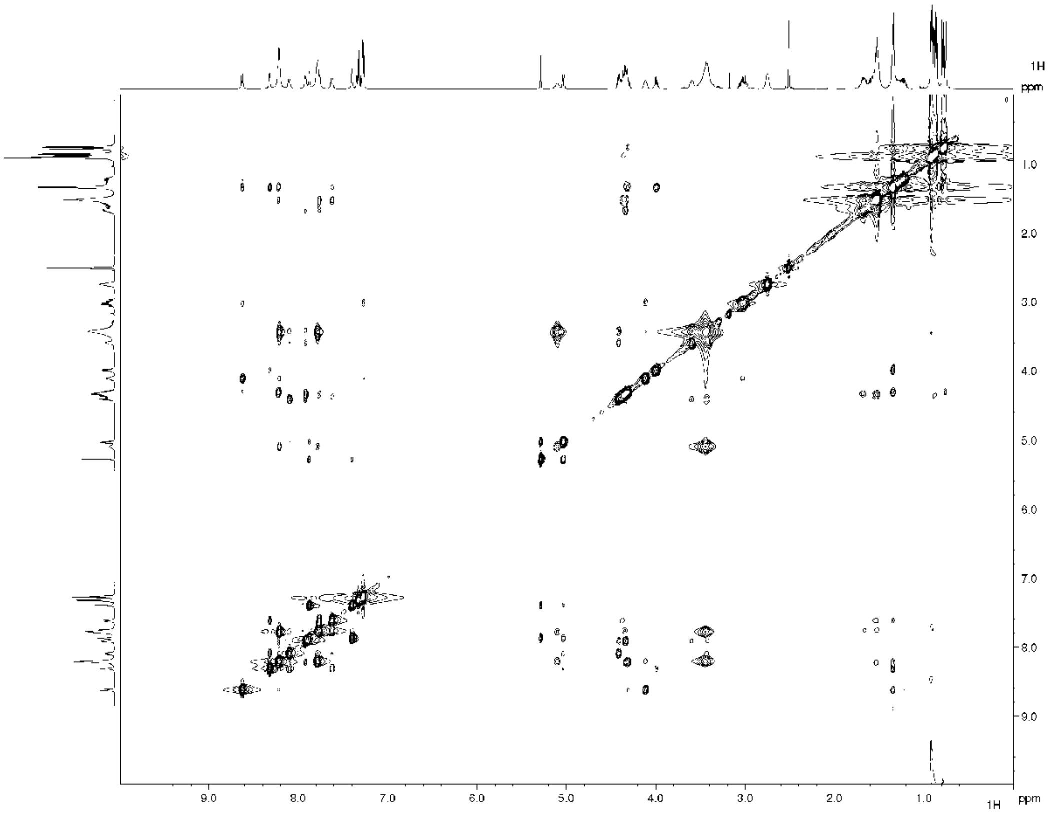

**Supplementary Figure 18.** $^{1}$H-$^{1}$H NOESY solution NMR Spectrum of clovibactin.

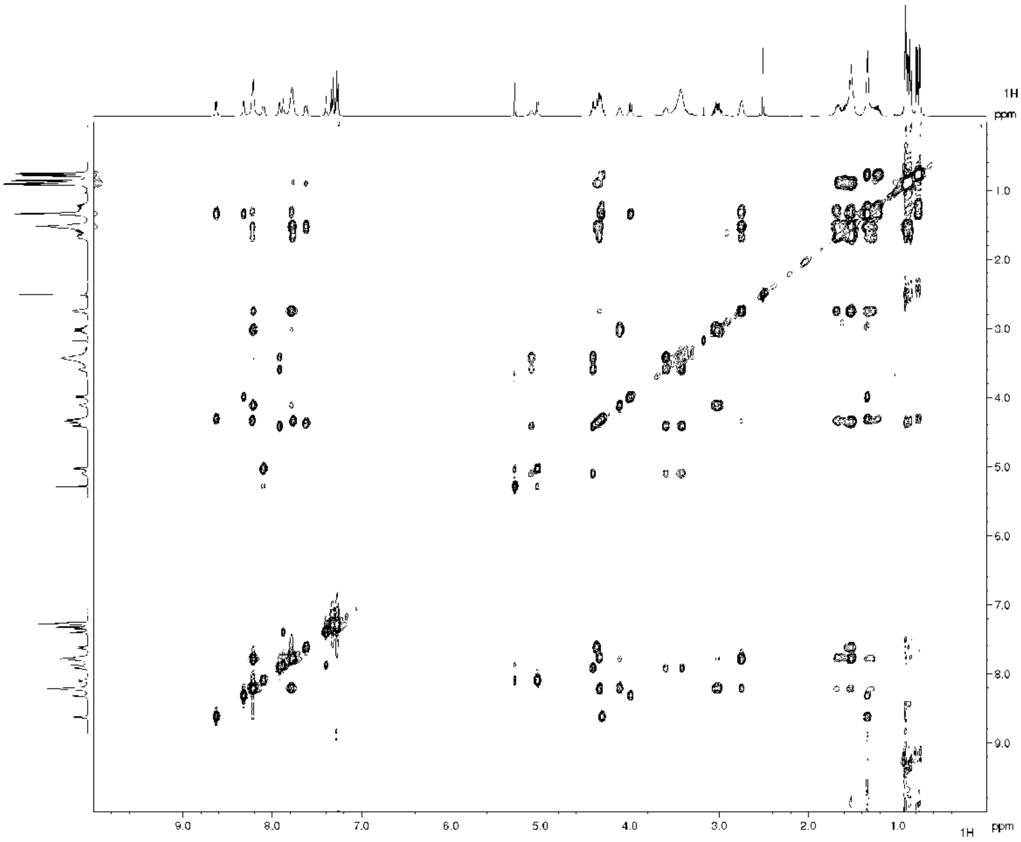

**Supplementary Figure 19.** $^{1}$H-$^{1}$H TOCSY solution NMR spectrum of clovibactin.

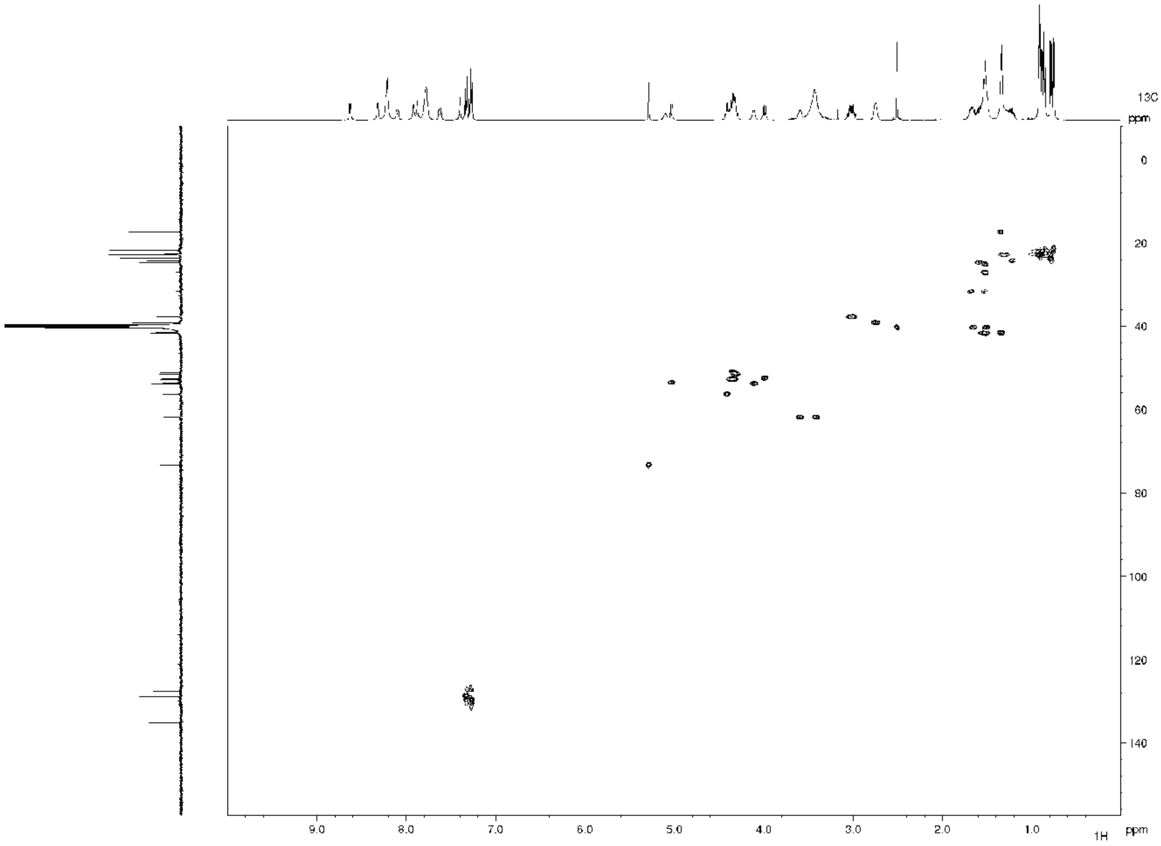

**Supplementary Figure 20.** $^{1}$H-$^{13}$C HSQC solution NMR spectrum of clovibactin.

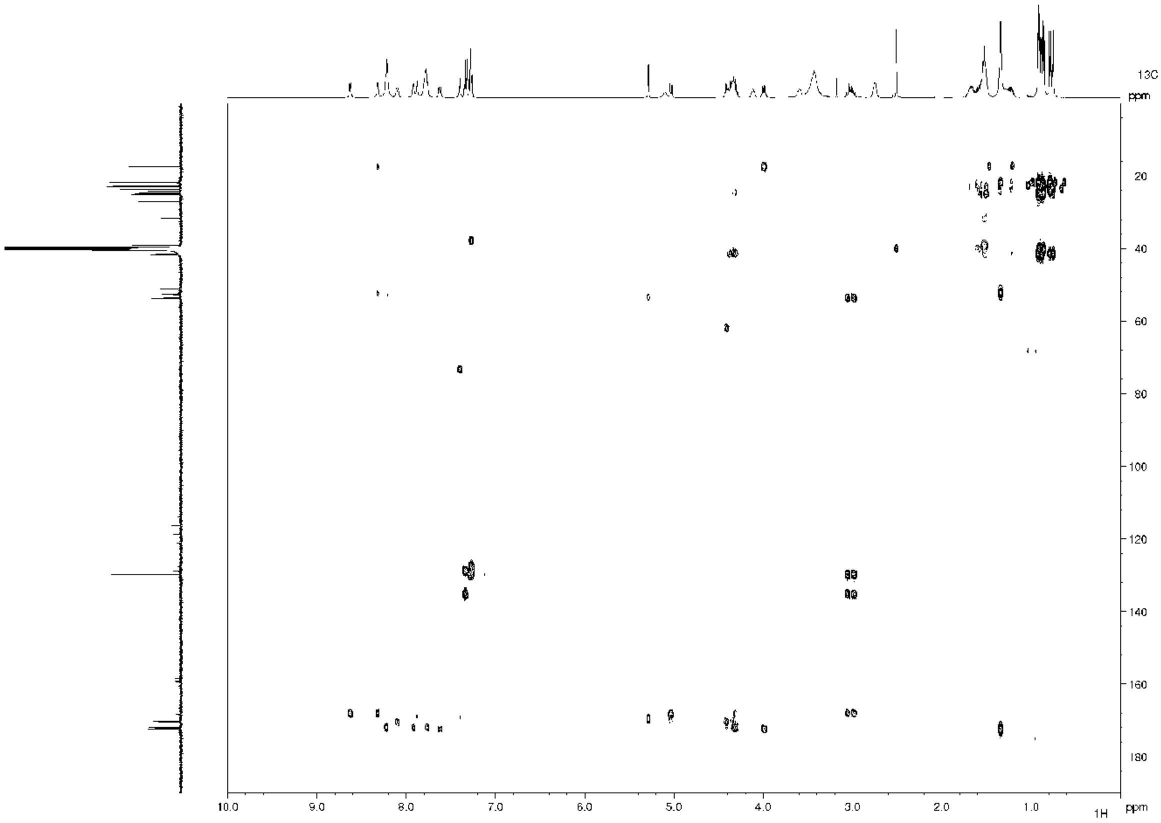

**Supplementary Figure 21.** $^{1}$H-$^{13}$C HMBC solution NMR spectrum of clovibactin.

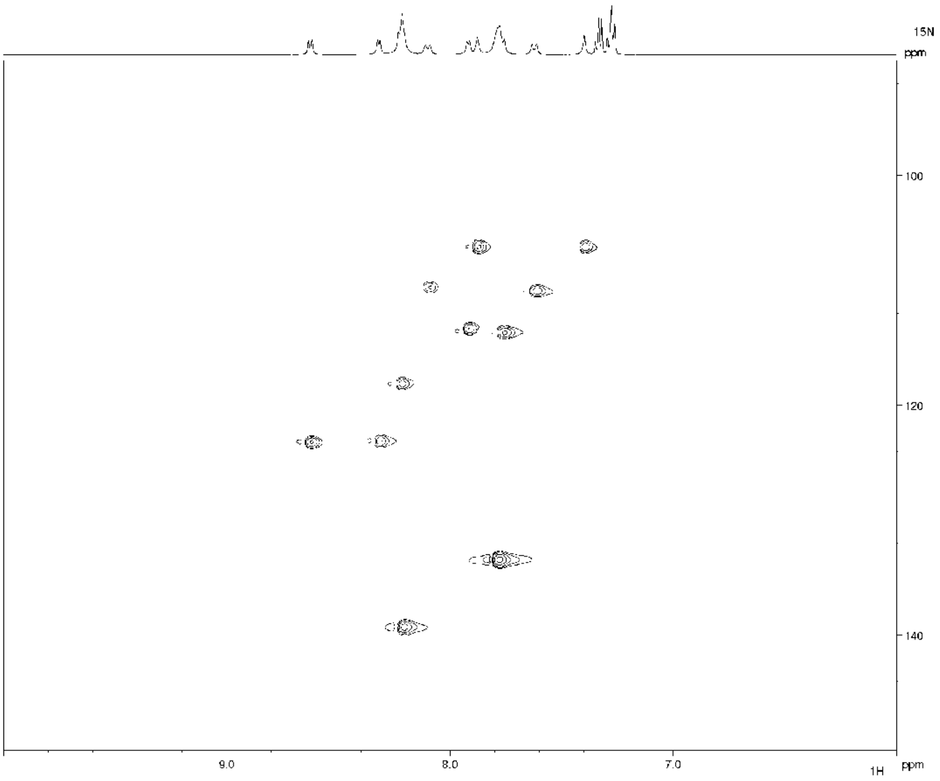

**Supplementary Figure 22.** $^{1}H$-$^{15}N$ HSQC solution NMR Spectrum of clovibactin.

### Supplementary Table 4

$^{13}C$,$^{15}N$-labelled clovibactin in the unbound state, obtained from solution NMR measurements, acquired at 600 MHz $^{1}H$-frequency in 30 mM Citrate buffer with 10% D2O, pH = 5.5. Assignments in ppm. Overlapping $^{13}C$-signals are shown in the same colour.

| Res | # | N | H(N) | Cα | Cβ | Nδ |
|---|---|---|---|---|---|---|
| Phe | 1 | | | 57.3 | 39.1 | |
| D-Leu | 2 | 127.9 | 8.4 | 55.5 | 42.2 | |
| D-Lys | 3 | 121.8 | 8.3 | 56.2 | 32.8 | |
| Ser | 4 | 117.7 | 8.1 | 58.8 | 63.9 | |
| Hyn | 5 | 112.9 | 8.9 | 56.4 | 76.1 | 107.3 |
| Ala | 6 | 126.1 | 8.1 | 55.3 | 18.4 | |
| Leu | 7 | 117.0 | 7.8 | 56.0 | 42.3 | |
| Leu | 8 | 118.6 | 7.9 | 54.2 | 42.1 | |

### Supplementary Table 5

$^{13}C$,$^{15}N$-labelled clovibactin in the lipid II bound state, obtained from ssNMR measurements in DOPC liposomes. Assignments in ppm. Overlapping $^{13}C$-signals (also with $^{13}C$,$^{15}N$-labelled lipid II) are colour coded.

| Res | # | N | H(N) | CO | Cα | Cβ | Cγ | Cδ1 | Cδ2 | Cε1/2 | Cζ | Nδ | Nζ | H(N)ζ | Hγ1+2 |
|---|---|---|---|---|---|---|---|---|---|---|---|---|---|---|---|
| Phe | 1 | 40.9 | | 172 | 58.5 | 39.7 | 140.2 | ~132 | ~132 | ~132 | ~132 | | | | |
| D-Leu | 2 | 127.7 | 8.8 | 178.6 | 57.7 | 43.7 | 27.4 | 25.7 | 24.7 | | | | | | |
| D-Lys | 3 | 113.0 | 7.7 | 177.4 | 56.1 | 35.2 | 27.0 | 31.1 | | 42.4 | | | 29.9 | 8.4 | |
| Ser | 4 | 113.8 | 9.9 | 176.6 | 61.3 | 65.9 | | | | | | | | | |
| Hyn | 5 | 110.7 | 9.0 | 171.7 | 56.3 | 75.3 | 174.1 | | | | | 108.8 | | | 8.4/6.1 |
| Ala | 6 | 125.9 | 8.2 | 179.6 | 55.9 | 17.4 | | | | | | | | | |
| Leu | 7 | 121.2 | 9.9 | 177.6 | 59.5 | 42.2 | 27.6 | 26.1 | 25.1 | | | | | | |
| Leu | 8 | 112.9 | 8.7 | 171.2 | 53.1 | 40.9 | 27.0 | 22.9 | 27.0 | | | | | | |

### Supplementary Table 6

$^{13}C$,$^{15}N$-labelled lipid II in the bound state with clovibactin, obtained from ssNMR measurements in DOPC liposomes. Assignments in ppm. Overlapping $^{13}C$-signals (also with $^{13}C$,$^{15}N$-labelled clovibactin) are colour coded.

| Sugars | # | C1 | C2 | C3 | C4 | C5 | C6 | C7 | C8 | C9 | C10 | C11 |
|---|---|---|---|---|---|---|---|---|---|---|---|---|
| MurNAc | | 97.5 | 56.5 | 80.0 | 71.2 | 77.6 | 64.6 | 176.9 | 25.2 | 81.8 | 178.3 | 20.4 |
| GlcNAc | | 103.3 | 59.3 | 78.6 | 73.7 | 76.5 | 64.5 | 175.6 | | | | |
| Pentapep | | CO | Cα | Cβ | Cγ | Cδ | Cε | | | | | |
| L-Ala | 1 | 176.9 | 53.5 | 20.0 | | | | | | | | |
| D-γ-Glu | 2 | | 34.4 | 30.9 | 57.4 | 180.7 | | | | | | |
| L-Lys | 3 | | 56.5 | 32.8 | 24.5 | 28.8 | 41.9 | | | | | |
| D-Ala | 4/5 | | 52.2 | 19.2 | | | | | | | | |

| | | |
|---|---|---|
| **PPi** | | **P** |
| **Pi** | **1** | - 8.4 |
| **Pi** | **2** | -13.0 |

### Analysis of calculated structure models

The average backbone RMSD (from the average structure) of the 22 clovibactin molecules in the complex was 2.50 ± 0.85 Å.

#### NMR restraints

Intermolecular distance restraints between clovibactin molecules

All restraints based on a series of 2D $^{13}C^{13}C$ PARIS experiments (up to 300 ms $^{13}C^{13}C$ transfer) using a target distance of 7.5 Å, and lower distance margin of 5.5 Å, and a higher distance margin of 2.0 Å; i.e., format (7.5 5.5 2.0). Note that in the arrangement of four clovibactin molecules, intermolecular clovibactin-clovibactin distance restraints for the *inner* clovibactin molecules were implemented so that a restraint could be with the clovibactin molecule on either side.

Detailed list of clovibactin – clovibactin restraints.

**Supplementary Table T4:**

**Clovibactin-Clovibactin contacts**
**Unambiguous**

| # | | $^{13}$C-CS (ppm) | | $^{13}$C-CS (ppm) | Average shortest intramolecular distance (Å) | Average shortest intermolecular distance (Å) |
|---|---|---|---|---|---|---|
| 1 | Phe1Cα | 58.5 | Hyn5Cβ | 75.3± | 11.8 ± 0.6 | 6.3 ± 0.7 |
| 2 | Phe1Cα | 58.5 | Hyn5Cγ | 174.1 | 10.4 ± 0.7 | 6.5 ± 0.9 |
| 3 | Phe1Cα | 58.5 | Ala6Cβ | 17.4 | 15.3 ± 0.5 | 6.5 ± 1.0 |
| 4 | Phe1Cα | 58.5 | Leu8Cα | 53.1 | 13.4 ± 0.8 | 8.8 ± 1.0 |
| 5 | Phe1Cα | 58.5 | Leu8CO | 171.2 | 12.6 ± 0.8 | 8.2 ± 0.9 |
| 6 | Phe1Cα | 58.5 | Leu8Cβ | 40.9 | 12.6 ± 0.9 | 8.1 ± 1.3 |
| 7 | Phe1Cα | 58.5 | Leu8Cδ1 | 22.9 | 13.6 ± 1.1 | 7.8 ± 1.6 |
| 8 | Phe1Cβ | 39.7 | Hyn5Cβ | 75.3 | 11.3 ± 0.9 | 5.4 ± 0.6 |
| 9 | Phe1Cβ | 39.7 | Hyn5Cγ | 174.1 | 9.9 ± 1.0 | 5.9 ± 0.7 |
| 10 | Phe1Cβ | 39.7 | Ala6CO | 179.6 | 15.4 ± 0.8 | 6.6 ± 0.9 |
| 11 | Phe1Cβ | 39.7 | Leu8CO | 171.2 | 12.1± 1.2 | 7.6 ± 0.6 |
| 12 | Phe1Cβ | 39.7 | Leu8Cδ1 | 22.9 | 13.2 ± 1.6 | 8.1 ± 1.7 |
| 13 | Phe1Cγ | 140.2 | Leu8Cα | 53.1 | 13.5 ± 1.6 | 8.5 ± 0.7 |
| 14 | Phe1Cγ | 140.2 | Leu8Cβ | 40.9 | 12.7 ± 1.7 | 8.3 ± 1.2 |

| | | | | | | |
|---|---|---|---|---|---|---|
| 15 | Phe1Caro | ~132 | Leu8Cβ | 40.9 | 13.5 ± 2.1 | 7.3 ± 1.3 |
| 16 | Leu2Cα | 57.7 | Leu8Cα | 53.1 | 11.2 ± 0.6 | 9.6 ± 1.0 |
| 17 | Leu2Cβ | 43.7 | Leu8CO | 171.2 | 10.9 ± 0.6 | 8.5 ±1.2 |
| 18 | Leu2CO | 178.6 | Ala6Cβ | 17.4 | 10.8 ± 0.5 | 8.6 ± 0.6 |
| 19 | Leu2CO | 178.6 | Leu8Cδ1 | 22.9 | 10.7 ± 0.7 | 9.6 ± 1.0 |

Hydrogen bond restraints (intermolecular) between clovibactin molecules: Hydrogen bonding restraints in line with experimentally determined antiparallel clovibactin arrangements were applied. While two different register shifts are possible to form antiparallel teixobactin [illegible]-sheets (i.e., Ser4NH could interact with Leu2CO or Ser4CO), only the variant with Leu2CO agrees with the intermolecular ssNMR distance restraints. Hydrogen bonding distance restraints were defined accordingly with upper and lower limits of 2.3 and 1.5 Å, respectively, i.e., format (2.0 0.5 0.3).

*clovibactin **A** with clovibactin **B***

Leu2NH - Ser4CO
Leu2CO - Ser4NH
Ser4NH - Leu2CO
Ser4CO - Leu2NH

*clovibactin **C** with clovibactin **D***

Phe1NH - Lys3CO
Phe1CO - Lys3NH
Lys3NH - Phe1CO
Lys3CO - Phe1NH

*clovibactin **C** with clovibactin **D***

Similar as in ***A*** with ***B***

Intermolecular distance restraints between clovibactin and lipid II: *Restraints involving the pyrophosphate group:* Ambiguous distance restraints were applied between the backbone amino protons of the four depsi-cycle residues (Hyn5-Leu8) with either phosphate of the pyrophosphate group using upper and lower limits of 2.4 and 1.7 Å, respectively, i.e., format (2.0 0.3 0.4). Restraints based on a 2D $^{1}H^{31}P$ experiment (Fig. 5b of the manuscript).

*Restraints involving the MurNAc/GlcNAc sugars:* Restraints were based on a series of 2D $^{13}C^{13}C$ PARIS and PARISxy experiments with 50, 250, and 300 ms magnetization transfer. Due to the mobility of the depsi-cycle in the complex (Figure 4C,D of the main text), most interfacial restraints were weak. We therefore applied all interfacial CC restraints with wider boundaries (7.5 5.5 2.0), i.e., upper and lower boundaries of 9.5 and 2.0 Å, respectively. Notwithstanding the wider boundaries, the interface is well defined (1.47±0.40 Å interfacial RMSD defined by residues Ala6, Leu7, Leu8 of clovibactin, PPi and MurNAc of lipid II)

**Supplementary Table T6:**

Detailed list of clovibactin – lipid II restraints.

| # | | $^{13}$C-CS (ppm) | | $^{13}$C-CS (ppm) | Average shortest interfacial distance (Å) |
|---|---|---|---|---|---|
| 1 | Ser4Cβ | 65.9 | Mu6C1 | 97.5 | 5.7 ± 0.9 |
| 2 | Ser4Cβ | 65.9 | Mu6C3 | 80.0 | 7.4 ± 1.0 |
| 3 | Ala6Cβ | 17.4 | Mu6C1 | 97.5 | 4.5 ± 0.6 |
| 4 | Ala6Cβ | 17.4 | Mu6C3 | 80.0 | 6.4 ± 0.6 |
| 5 | Ala6Cβ | 17.4 | Mu6C5 | 77.6 | 5.0 ± 0.9 |
| 6 | Ala6Cβ | 17.4 | Mu6C9 | 81.8 | 7.6 ± 0.8 |
| 7 | Ala6Cβ | 17.4 | Mu6C11 | 20.4 | 7.8 ± 1.1 |
| 8 | Ala6Cβ | 17.4 | Gl7C3 | 78.6 | 7.2 ± 1.9 |
| 9 | Leu7Cβ | 42.2 | Mu6C1 | 97.5 | 4.4 ± 0.6 |
| | Lys3Cε | 42.4 | Mu6C1 | 97.5 | 8.1 ± 0.9 |
| 10 | Leu7Cβ | 42.2 | Mu6C3 | 80.0 | 5.6 ± 0.5 |
| | Lys3Cε | 42.4 | Mu6C3 | 80.0 | 8.5 ± 1.3 |
| 11 | Leu7Cβ | 42.2 | Mu6C5 | 77.6 | 5.6 ± 1.0 |
| | Lys3Cε | 42.4 | Mu6C5 | 77.6 | 9.2 ± 0.9 |
| 12 | Leu8CO | 171.2 | Mu6C1 | 97.5 | 5.8 ± 0.8 |
| 13 | Leu8Cδ1 | 81.7 | Mu6C3 | 80.0 | 6.0 ± 1.1 |
| 14 | Leu8Cδ1 | 81.3 | Gl7C3 | 78.6 | 8.6 ± 1.1 |
| 15 | Leu8Cδ1 | 81.7 | Gl7C5 | 76.5 | 7.0 ± 0.7 |
| 16 | Leu8Cγ | 27.0 | Mu6C1 | 27.0 | 5.7 ± 0.8 |
| | Lys3Cγ | 27.0 | Mu6C1 | 27.0 | 7.9 ± 1.0 |

The shortest distances were averaged over the 22 structure models of the ensemble. The error shows the standard deviation.

Topological restraints: Eventually, a filtering strategy was applied to constrain the conformational space of the isoprenyl tails. Structure models were only accepted if all lipid II tails pointed into the direction of the membrane-exposed sidechain of Leu2, Ile5, and Ile6 (see Fig. 4I of the main text and Supplementary Fig. 7). Sorting of the lipid II tails was steered by imposing soft distance restraints between the sidechain of Leu2 and the lipid II isoprenyl-tail, and by imposing soft distance restraints between the isoprenyl-tails.

**Supplementary Table T7:**

Number and type of restraints used for the structure calculations. Intramolecular restraints are listed per monomers, while intermolecular/interfacial restraints are listed per pair of interacting molecules.

| ***Number of Restraints*** | | |
|---|---|---|
| ***Intermolecular*** | *Unambiguous* | *Ambiguous* |

| ***clovibactin - clovibactin*** | | |
|---|---|---|
| Distance restraints | 19 | 0 |
| Hydrogen bonds** | 4 | 0 |
| ***Intermolecular clovibactin - lipid II*** | *Unambiguous* | *Ambiguous* |
| Distance restraints with sugars | 12 | 4 |
| Distance restraints with PPI*** | 0 | 4 |

*,**,*** See Methods (NMR Structure calculations) for details

**Analysis of calculated structure models**
Structural and violation statistics of the final 22 structure models are given below:

Structure ensemble precision:

- Average backbone RMSD (from the average structure) of the 22 clovibactin molecules in the complex:  2.50 +/- 0.85 Å.

**Details of 2D ssNMR experiments**

1. 2D CC PARIS experiment with $^{13}C$,$^{15}N$-clovibactin – $^{12}C$,$^{14}N$-lipid II

Magnetic field /MAS = 1200 MHz ($^{1}H$-frequency) / 18 kHz
Mixing time (CC) = 50 ms
t1 points / AQ = 452 / 5.00 ms
Recycle delay = 2.40 s
Co-added transients = 144
Experimental time = 1d 21h

2. 2D CC PARIS experiment with $^{13}C$,$^{15}N$-clovibactin – $^{12}C$,$^{14}N$-lipid II

Magnetic field /MAS = 1200 MHz ($^{1}H$-frequency) / 18 kHz
Mixing time (CC) = 300 ms
t1 points / AQ = 452 / 5.00 ms
Recycle delay = 2.40 s
Co-added transients = 224
Experimental time = 3d 5h

3. 2D CC PARISxy (m = 1) experiment with $^{13}C$,$^{15}N$-clovibactin – $^{13}C$,$^{15}N$-lipid II

Magnetic field /MAS = 950 MHz ($^{1}H$-frequency) / 15.5 kHz
Mixing time (CC) = 50 ms
t1 points / AQ = 316 / 4.40 ms
Recycle delay = 1.90 s
Co-added transients = 192
Experimental time = 1d 10h

4. 2D CC PARISxy (m = 1) experiment with $^{13}C$,$^{15}N$-clovibactin – $^{13}C$,$^{15}N$-lipid II

Magnetic field /MAS = 950 MHz ($^{1}H$-frequency) / 15.5 kHz

Mixing time (CC) = 300 ms
t1 points / AQ = 215 / 3.00 ms
Recycle delay = 1.84 s
Co-added transients = 1024
Experimental time = 5d 14h

5. 2D CC PARIS experiment with $^{13}C$,$^{15}N$-clovibactin – $^{13}C$,$^{15}N$-lipid II
Magnetic field /MAS = 1200 MHz ($^{1}H$-frequency) / 18 kHz
Mixing time (CC) = 50 ms
t1 points / AQ = 452 / 5.00 ms
Recycle delay = 2.40 s
Co-added transients = 144
Experimental time = 1d 21h

6. 2D CC PARIS experiment with $^{13}C$,$^{15}N$-clovibactin – $^{13}C$,$^{15}N$-lipid II
Magnetic field /MAS = 1200 MHz ($^{1}H$-frequency) / 18 kHz
Mixing time (CC) = 250 ms
t1 points / AQ = 316 / 3.50 ms
Recycle delay = 2.30 s
Co-added transients = 752
Experimental time = 7d 1h

7. 2D T2-edited H(H)C experiment with $^{13}C$,$^{15}N$-clovibactin – $^{12}C$,$^{14}N$-lipid II
Magnetic field /MAS = 700 MHz ($^{1}H$-frequency) / 16.5 kHz
Mixing time (HH) = 5 ms
T2 filter = 2.5 ms
t1 points / AQ = 60 / 2.08 ms
Recycle delay = 2.15 s
Co-added transients = 1972
Experimental time = 2d 17h

8. 2D CC TOBSY experiment with $^{13}C$,$^{15}N$-clovibactin – $^{13}C$,$^{15}N$-lipid II
Magnetic field /MAS = 950 MHz ($^{1}H$-frequency) / 8 kHz
Mixing time (HH) = 6 ms
t1 points / AQ = 265 / 5.38 ms
Recycle delay = 1.61 s
Co-added transients = 2304
Experimental time = 11d 16h

9. 2D NCA experiment with $^{13}C$,$^{15}N$-clovibactin – $^{12}C$,$^{14}N$-lipid II
Magnetic field /MAS = 800 MHz ($^{1}H$-frequency) / 15 kHz
Mixing time (NC) = 5.5 ms
t1 points / AQ = 17 / 3.6td ms
Recycle delay = 1.82 s
Co-added transients = 7168

Experimental time = 2d 17h

<u>9. 2D NCO experiment with $^{13}C$,$^{15}N$-clovibactin – $^{12}C$,$^{14}N$-lipid II</u>

| | |
|---|---|
| Magnetic field /MAS | = 800 MHz ($^{1}H$-frequency) / 15 kHz |
| Mixing time (NC) | = 7 ms |
| t1 points / AQ | = 25 / 4.4 ms |
| Recycle delay | = 1.9 s |
| Co-added transients | = 8192 |
| Experimental time | = 4d 18h |

<u>10. 2D NH experiment with $^{13}C$,$^{15}N$-clovibactin – $^{12}C$,$^{14}N$-lipid II</u>

| | |
|---|---|
| Magnetic field /MAS | = 700 MHz ($^{1}H$-frequency) / 60 kHz |
| t1 points / AQ | = 87 / 5.6 ms |
| Recycle delay | = 0.72 s |
| Co-added transients | = 4096 |
| Experimental time | = 5d 4h |
| Note | = high number of co-added transients was necessary to obtain good data for the sidechains of 3 and 5. |

*For all 2D experiments, sign discrimination in indirect dimensions was achieved with the TPPI (time-proportional phase incrementation) method.*

**Supplementary Video 1: Real-time observation of the growth of clovibactin fibrils on membrane captured by HS-AFM.** Membrane containing 4 % lipid II and 5 μM of clovibactin is used. The imaging rate is 0.5 frames/second with line rate of 150 lines/second. Clovibactin is added at 0 sec.
